# scPRINT-2: Towards the next-generation of cell foundation models and benchmarks

**DOI:** 10.64898/2025.12.11.693702

**Authors:** Jérémie Kalfon, Gabriel Peyré, Laura Cantini

## Abstract

Cell biology has been booming with foundation models trained on large single-cell RNA-seq databases, but benchmarks and capabilities remain unclear. We propose an additive benchmark across a gymnasium of tasks to discover which features improve performance. From these findings, we present scPRINT-2, a single-cell Foundation Model pre-trained across 350 million cells and 16 organisms. Our contributions in pre-training tasks, tokenization, and losses made scPRINT-2 state-of-the-art in expression denoising, cell embedding, and cell type prediction. Furthermore, with our cell-level architecture, scPRINT-2 becomes generative, as demonstrated by our expression imputation and counterfactual reasoning results. Finally, thanks to our pre-training database, we uncover generalization to unseen modalities and organisms. These studies, together with improved abilities in gene embeddings and gene network inference, place scPRINT-2 as a next-generation cell foundation model.

## Introduction

For the last few years, Single-Cell Foundation Models (scFMs), also known as Virtual Cell models, have provided early approaches to modeling the cell using single-cell RNA-seq data as their primary modality^1–4^. The field has been booming with these transformer-based machine learning models trained on large databases of tens of millions of cells. The models themselves contain tens to hundreds of millions of parameters and are trained on unsupervised (or semi-supervised) tasks such as predicting masked gene expression or denoising expression. They can then be used as is to examine their learned representations or fine-tuned to transfer their knowledge across a range of everyday tasks in that modality. Many examples have now been proposed, such as predicting single-cell perturbation responses, patient drug responses, and disease states; annotating cells; correcting for batch effects; improving noise levels; imputing unseen gene expression or modality; generating gene networks; identifying cell niches; and more^5,6,6–15^.

While many AI Virtual Cell models and scFMs exist, little has been done regarding their comparison^16–20^. A crucial question remains: how to validate the impact of the different proposed methods, regardless of implementation, datasets, or model size. Indeed, reproducing results has been challenging for many, and the literature has yielded discordant conclusions about the performance and capabilities of these models. Showing they often underperform simpler approaches on classification, batch correction, and perturbation prediction^16,17,21–23^.

Much work remains to get to feature-rich, easy-to-use scFMs. Models that allow inference in minutes, along with well-crafted reproducible benchmarks that demonstrate how scFMs uniquely solve essential problems in single-cell biology. Open-sourcing not just model weights but their pre-training tasks and datasets.

On this front, scPRINT was released as part of a second batch of scFM, presenting contributions in terms of usability and reproducibility while also showcasing pre-training strategies, data encoding, and decoding^3^. scPRINT was trained on 50 million cells using a multitask pre-training strategy that included expression denoising, autoencoding, and cell-label prediction. It also presented an in-depth benchmark that examined the foundation model’s zero-shot performance on these tasks, as well as its internal gene network representation and fidelity compared to multiple ground truths.

Building on these strengths and moving towards the next generation of scFMs, we here use scPRINT (which will be referred to as scPRINT-1) as the reference to showcase an extensive additive benchmark of scFM attributes. We address several key questions about the importance of diverse architectures, datasets, and training modalities. This additive benchmark aims to understand the relative importance of these different features in our task gymnasium, examining the choice of model architecture and pre-training tasks across 42 different scenarios. In these scenarios, we propose a breadth of novel components for scFMs. In addition to those 12 distinct contributions, we also examine various pre-training datasets, compiling a 350-million-cell database—the largest to date—with over 16 organisms.

As a result of the benchmark, we derive a next-generation scFM, **scPRINT-2**. scPRINT-2 improves upon the previous generation of models by leveraging our database, the scPRINT-2 corpus, and multiple data augmentation approaches. It uses a set of updated pre-training tasks and losses, improving its accuracy in challenging and unseen contexts. Finally, it is equipped with graph-based encoders and the XPressor architecture, enabling unprecedented expression imputation, high-quality zero-shot embeddings, and counterfactual reasoning. We dive into these specific contributions by examining multiple use cases, highlighting behaviors that are often overlooked or under-assessed in classical benchmarks. scPRINT-2, its dataloader, pre-training datasets, preprocessing, task functions, pre-trained weights, as well as the additive benchmark training traces and all 42 models’ weights are fully open-sourced and available under the GPL-v3 License.

## Results

### Decoding the impact of a foundation model’s architecture through an additive benchmark

Many scFMs have been developed in single-cell genomics. They have mostly been studied in isolation, using their own benchmarks. While most of them maintained relatively similar architectures, the impact of each design’s decisions was never thoroughly assessed. For example, scPRINT-1 uses a denoising reconstruction task similar to scFoundation. Still, scFoundation uses the mean-squared-error (**MSE**) for the reconstruction loss, whereas scPRINT-1 uses the zero-inflated negative-binomial loss (**ZiNB**) (see <u>Methods</u>). scGPT and Geneformer utilize masking, but scGPT bins expression counts (**binning**), while scPRINT-1 does denoising and employs a continuous embedding with a log transform and a pseudocount of 1 (**logp1**)^1,2^. Other models, like cellPLM, instead use a contrastive learning approach, which encourages embeddings of perturbed and unperturbed cell profiles to be more similar to each other than those of different cell profiles^4^. This method is also known as InfoNCE or Contrastive Cell Embedding (**CCE**) (see <u>Methods</u>)^24^.

### Additive benchmark

To address the lack of a consistent assessment of these models, we have designed a benchmark to comprehensively evaluate the various components of scFMs, including pre-training databases, architectures, and training tasks. This benchmark is based on a gymnasium of tasks similar to those presented in Kalfon et al.^3^ (see Figure 1A; see Table 1). The scFM gymnasium assesses each model’s ability to predict labels, remove batch effects, denoise, and impute gene expression, as well as discover known gene-gene relationships at different stages of training. For embeddings and cell type classification, we use the scIB and accuracy scores over the same ground-truth test datasets as in Kalfon et al. (see <u>Methods</u>). For denoising, we evaluate the model’s ability to recover the noised expression profile of cells from a test dataset, as measured by the improvement in correlation with the ground-truth profile after denoising. For gene-network inference, we examine the Odds Ratio (OR) and AUPRC scores of the model’s ability to recover a ground-truth gene network from expression data alone (see <u>Methods</u>).

**Figure 1:**
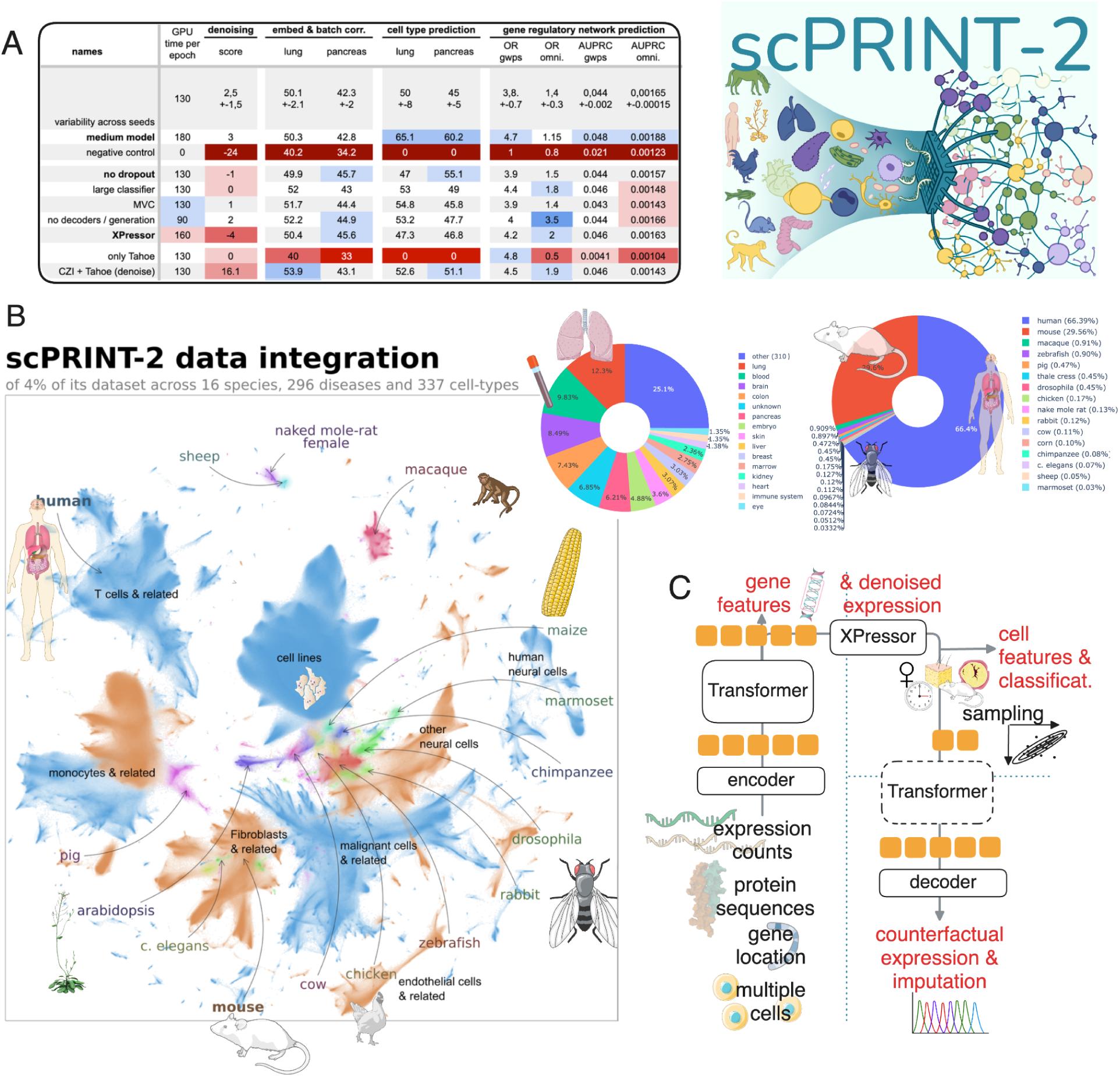
presentation of the scPRINT-2 model, pre-training dataset, and additive benchmark. (a) The additive benchmark example table with its gymnasium scores across the scFM’s features. (b) Our scPRINT-2 corpus pre-training dataset, with 16 organisms across 300+ tissues. Umap of 15 million cells from the corpus integrated using scPRINT-2. Colors represent species. (c) The scPRINT-2 model, its input data, and its different outputs. Source data are provided as a Source Data file.

**Table 1:**
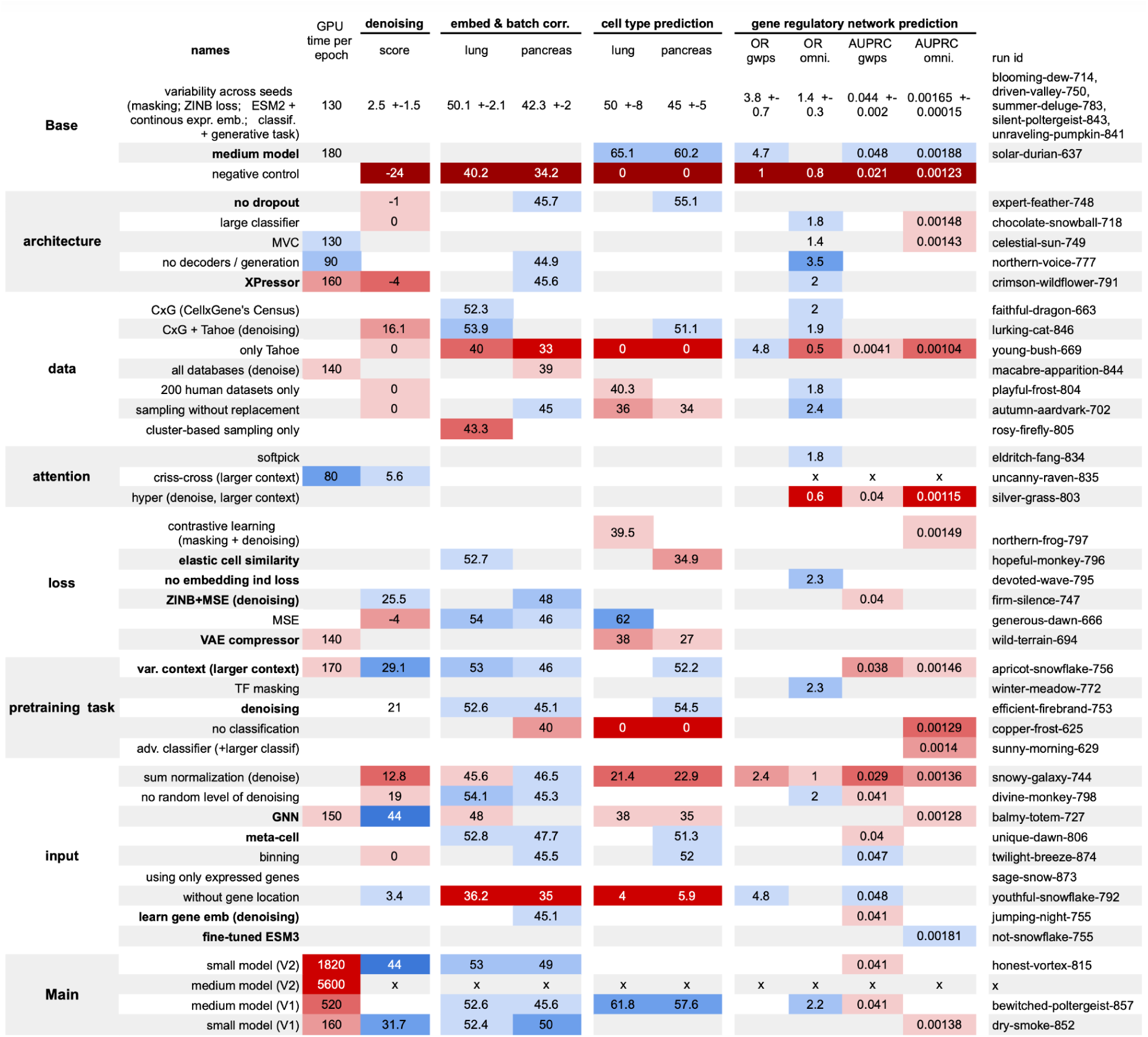
Full results of the additive benchmark. . Table representing the results of the additive benchmark on 42 models, over multiple metrics: batch correction and cell embedding quality, denoising quality, cell type prediction, and gene network inference. Additional information on the different components is available in the methods section. Bold elements are the features that are part of the scPRINT-2 foundation model.

The base model, on which the additive benchmark is performed (see Figure 1A, Table 1, and <u>Methods</u>), is trained on the CxG database, comprising 500 carefully annotated human and mouse datasets. Its training lasts for a maximum of 20 epochs, each of 20,000 steps, with a minibatch size of 64. We encode the gene expression using the scPRINT-1 approach and decode it with the MSE method. The base model’s pre-training task uses a 30% gene expression mask. We pre-train the models 6 times across multiple seeds to generate error bounds. Using Flash-Attention-3, the 20M parameters model trains on 1 H100 GPU for 2 days.

While we will not delve into the details of each feature assessed (see Methods), our benchmark broadly highlights several key points.

Regarding the tasks, we have confirmed what Kalfon et al. and De Waele et al. previously showed: that denoising is superior to masking as a pre-training task for single-cell data in classification and embedding tasks^3,25^. Similarly, un-normalized expression is better than normalizing it at the input. Classification also serves as a good supplement to the pre-training task, as without it, we observe a slight decrease in performance (see Table 1).

We also present, as part of our study, the **scPRINT-2 corpus**, which comprises more than 350 million single cells (see Figure 1B). This is the largest dataset ever assembled, consisting of data from the Chan Zuckerberg Institute’s Cellxgene (**CxG**), the **Tahoe**-100M dataset, and the scBasecount database, which contains 20,000 reprocessed datasets from the Gene Expression Omnibus^26–28^. The cells themselves are derived from 16 different eukaryotic organisms, spanning more than 1 billion years of evolution. The dataset comprises approximately 400,000 distinct genes, 4,764 different labels, and around 140,000 cell groups, totaling 25 TB of unique data^29^. Our database contains nine main classes: *cell type, disease, age, tissue of origin, assay, ethnicity, sex, cell culture,* and *organism*.

Thanks to this database, we demonstrated the growing importance of data selection in pre-training scFMs. Indeed, when using the Tahoe-100M database solely for pre-training, the model’s overall performance plummets, as the sequencing depth and diversity are low despite the large number of cells.

However, including this lower-diversity dataset with the high-diversity CxG database and carefully considering the cell-state imbalances results in only a noticeable decrease in denoising performance. Interestingly, using all available datasets did not change performance across our benchmarks. Reducing the training database to a random subset of only **200 human datasets only,** led to a minimal decrease in denoising and cell type prediction. This shows again that the benchmark fails to highlight abilities on more diverse cell types and organisms^30,31^. But it also indicates diminishing returns in adding more datasets—diversity in cell states and organisms being much more important than cell count.

We thus preprocessed each dataset by removing all duplicates, filtering for low-quality cells, aligning metadata to the CxG ontologies, and computing cell-cell similarity profiles and clusters. It allowed us to introduce multiple data augmentation techniques, such as varying the input context length (**var. context**) during training and randomly creating **meta-cells**, which are averages of similar cell expression profiles across K-nearest neighbors (K-NN) (see <u>Results</u> section 3). Interestingly, we observe that both methods tend to improve the model’s performance in most metrics, even though these models do not examine more cells overall. This highlights the importance of effective data augmentation techniques for scFM pre-training^32^.

Regarding architecture, we recomputed results from the XPressor manuscript^33^, which showed that this architecture improves the embedding quality of scFMs (see <u>Results</u> section 4; see the full table in supp Table S1). We also demonstrate that using ESM-based gene ID tokens leads to much better performance than learning gene tokens from scratch^34^. Providing each gene’s genomic location as additional input information significantly improves model convergence. However, we also noticed that when they do converge, models without gene location information can perform well. We have noticed that model size correlates with higher scores, at least for gene network inference and cell-type prediction. Using a Graph Neural Network (**GNN**) encoder shows significant improvements, with only a slight decrease in the cell-type prediction task (see <u>Results</u> Section 3; see <u>Methods</u>). Additionally, our sub-quadratic attention mechanism, Criss-cross attention, also shows substantial benefits with no reduction in performance (see <u>results</u> section 4; see <u>Methods</u>).

Moreover, MSE, on average, outperforms ZiNB as a loss function while decreasing the model’s expressivity (see <u>Methods</u>). A good proposed middle ground is the ZiNB+MSE loss (see <u>Results</u> Section 3; see <u>Methods</u>).

Some unexpected results showed that omitting the decoder part of scPRINT-1 led to stronger performance; however, this comes at the cost of generative abilities and decreased cell-embedding fidelity. Indeed, despite its importance for understanding scFMs’ behavior and feature importance, we have noted that our benchmark does not yet capture the full breadth of abilities that scFMs do or should have. For example, both scIB and classification scores are very dependent on the dataset’s quality and its labels. Scores presented here show only a facet of the model’s ability. We might be interested in the model’s performance up-to-convergence instead of stopping them at 20 epochs or looking at unseen species, or assays at training. This is a first attempt to benchmark scFMs, but more extensive efforts will be needed.

### scPRINT-2

Overall, we have examined the performance improvements driven by our 12 distinct contributions across 42 training runs. Based on these results and our own considerations, we have elected a set of features to create scPRINT-2, a next-generation cell foundation model (see supp. Figure S1; see <u>Methods</u>). We highlight its architecture in Figure 1C; scPRINT-2 is currently available in a small version with only 20M active parameters. Its encoder-compressor-decoder architecture produces cell- and gene-level outputs at multiple levels, working on one or more cells at a time.

Furthermore, to aid in the exploration of this largest-ever cross-organism single-cell dataset, we release all of the 350 million cells in the scPRINT-2 corpus, aligned into an atlas by scPRINT-2, of which 1% are directly accessible through an interactive visualization (see Figure 1B, see <u>Data</u> <u>availability</u>) along with scPRINT-2 cell label predictions for all classes. This should enable never-before analysis and exploration of single-cell RNA-seq data.

But the additive benchmark leaves some questions unanswered about the effect of combining these features up-to-convergence and the models’ abilities on unseen modalities, tasks, and species. In the following sections, we will focus on 1. looking at more diverse and truthful datasets in size, quality, and source domains; 2. using more scores and ground truth validations; 3. defining tasks that better reflect the possibilities and real-life use of these models.

### A diverse dataset of 350 million cells pushes generalization to unseen organisms

One of the most critical features of foundation models (FMs) is the breadth of their training dataset. From vision to language, AI advancement has been driven by training models on ever-larger datasets^35–39^. Nowadays, most scFMs are trained on 20 to 50 million cells, except the recently released Geneformer-v2 and STATE-SE models, which have been trained on roughly 300 million cells^40,41^.

### scPRINT-2 pre-training corpus

In conjunction with our model’s architecture, the scPRINT-2 corpus and its 16 organisms enable generalization to organisms unseen during training. This broader cell type diversity, however, comes with additional challenges: annotation quality has decreased due to missing annotations in scBasecount. Additionally, the skew toward low sequencing depth and highly similar cells has increased with the inclusion of spatial transcriptomics datasets and less curated databases such as Tahoe-100M and Arc’s scBasecount (see <u>Methods</u>).

Fortunately, a key feature of our dataloader, scDataLoader^3^, is its ability to perform weighted random sampling, thereby mitigating the heavy dataset imbalances that currently exist across diverse cell types, sequencing methodologies, and different organisms assessed. We thus present methods to successfully train scPRINT-2 on this large dataset. The first, called cluster-weighted sampling, allows datasets with unclear annotations to benefit from weighted random sampling by defining clusters of high expression similarity (see <u>Methods</u>). This lets us define cell states without requiring any label information and perform sampling that is aware of the different cell states, regardless of the size of each cluster. We address the second issue of uneven cell quality by also skewing sampling toward cells with more non-zero genes (nnz). Both methods were enabled on such a vast database thanks to essential updates to scDataLoader. This re-weighting is performed jointly with weights on cell type, disease, organism, and sequencer labels, thereby addressing the size/diversity issues that plague these larger cell databases^42^.

Interestingly, the number of training steps required to achieve convergence increased only 2-fold, indicating that, as in scPRINT-1, the model did not sample as many cells as actually exist in the pre-training dataset before reaching convergence. However, with data augmentation and nearest-neighbor sampling, the model still encountered roughly 2 billion distinct input cell profiles during pre-training, corresponding to 2000 cell profiles per step.

After implementing this feature and training scPRINT-2, its cell-type classification performance on the validation dataset was 76%. For its other predicted labels, its performance was 59% (disease), 96% (ethnicity), 96% (assay), 94% (age), 100% (cell culture), 100% (organism), 93% (sex), and 70% (tissue of origin).

On the live benchmark Open Problem from November 2025, it achieved an average zero-shot performance of 75%, putting it above scPRINT-1 (47%) and other zero-shot FMs (40-60%), even above Liger, a supervised technique^43^ (see Figure 2A; see <u>Methods</u>). But scPRINT-2 was the only scFM with UCE that could run on all datasets^44^. Against the two human datasets on which Scimilarity-KNN could be run, it performed slightly better than scPRINT-2. This is most likely due to the smaller capacity of scPRINT-2 (20M parameters) compared to scimilarity (100M parameters), as we also observed in our additive benchmark (see Table 1). Another likely reason is that the model likely saw those datasets more often during pre-training, since it is trained only on CxG’s human datasets.

**Figure 2:**
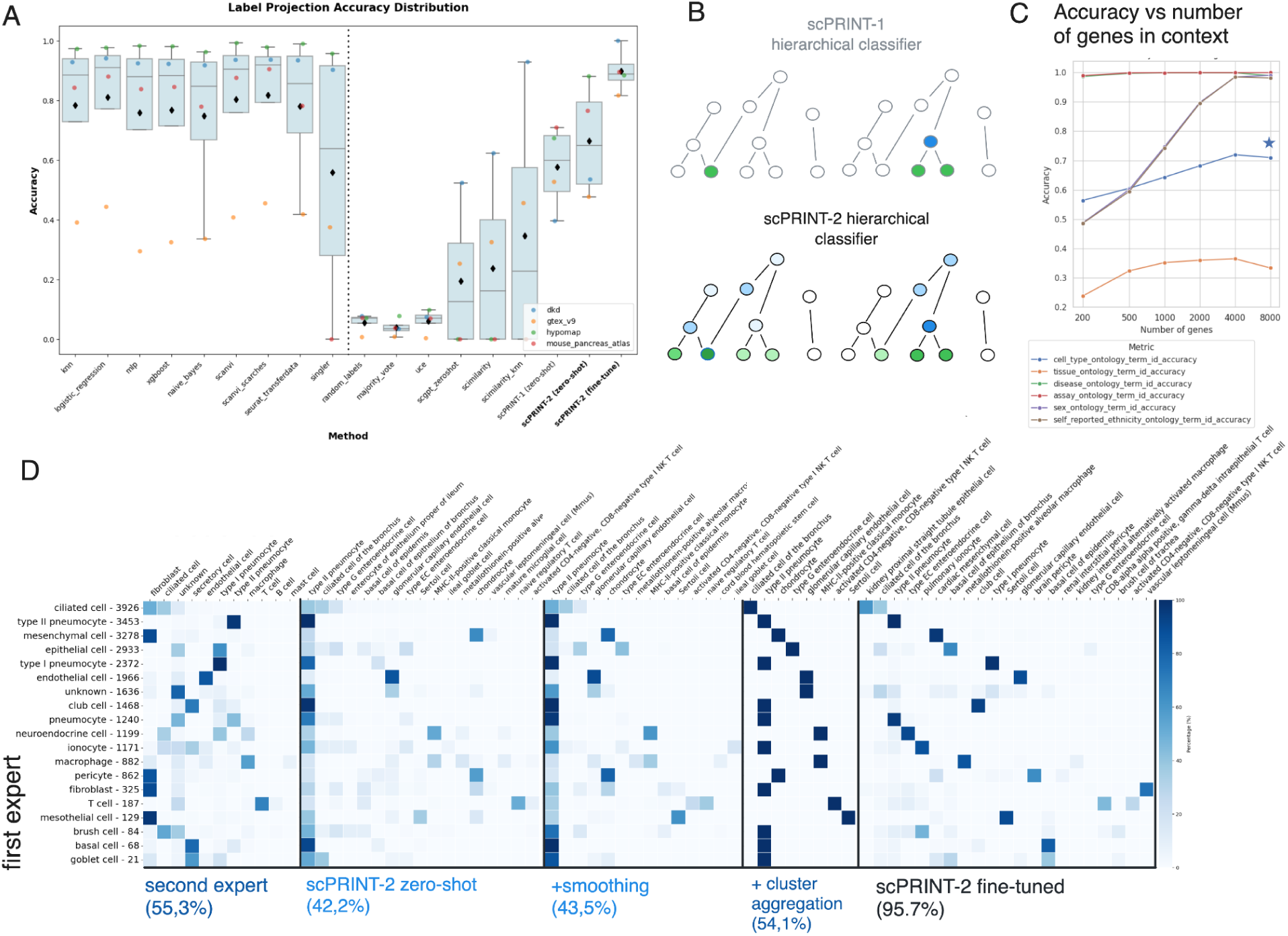
presentation of the updated classifier, and results on classification tasks. (a) Open Problems benchmark results and comparison of scPRINT-1 and zero-shot and fine-tuned scPRINT-2. (b) Illustration of our updated hierarchical classifier loss. (c) Unseen organisms cell type classification for cat and tiger datasets, across two experts and scPRINT-2 zero-shot, after label smoothing, after cluster aggregation, and after fine-tuning. (d) Change in classification accuracy as the number of genes in context increases for high-quality single-cell datasets. The star represents the model’s score when label smoothing is used.

We then performed fine-tuning using our XPressor-based Parameter-Efficient Fine-tuning (XPEFT), in which we fine-tune only the XPressor layers of scPRINT-2 (see <u>Methods</u>). In this context, we show that scPRINT-2 fine-tuned outperforms every existing supervised and unsupervised method on the Open-problem (see Part 4; see <u>Methods</u>)^45^. We observed similar trends in the macro-F1 scores (see supp. Figure S2). Of note, neither scGPT nor Geneformer are currently tested in their fine-tuned version on the platform.

These performances are enabled in part by our update to scPRINT-1’s hierarchical classification loss (see Figure 2B). The scPRINT-1 classifier generates predictions for all possible labels in a hierarchical ontology, while producing logits only for the leaf labels. To predict the other labels, it only has to aggregate their leaf logits. In scPRINT-2, we improve on this loss by using the entire ontological graph, meaning that, e.g., given a ground truth of *olfactory neuron,* we will penalize a prediction of *inhibitory neuron* less overall than a non-neuron label, like *fibroblast*. In conjunction with our weighted sampler, this allows the model to learn rich gradients from a low volume of data.

### scPRINT-2 generalizes to unseen classification tasks

We have, however, noticed that classification performance does not generalize sufficiently to correctly recover the exact phylogenetic relationships within organisms or, similarly, within ethnicities (see supp. Figures S3, S4, S5). This could be biased heavily by tissue representation in rare ethnicities and organisms. However, some relationships were found, such as *Singaporean Indian/Singaporean Chinese, Korean/Japanese/Chinese, American/Latin American,* or *Macaque/Marmoset/Chimpanzee, Drosophila/C. elegans, Human/Mouse, Pig/Cow*, suggesting that with greater diversity and representation, scFMs might learn this relationship classification of gene expression on their own.

We show that this does not prevent scPRINT-2 from generalizing to unseen organisms. Using a randomly selected tomato plant dataset and its corresponding ESM3 gene embeddings, unseen at training time, scPRINT-2 generates an organism label prediction for the two plant organisms it knows about 67% of the time. This is despite the very low prevalence of these organisms in the pre-training dataset (see Figure 1B). For a horse dataset, scPRINT-2 predicted mammalian organisms 72% of the time.

Unfortunately, these datasets lacked cell-type annotations. Using well-annotated datasets from Zhong et al.^46^ of cat and tiger lung tissues, organisms not seen at training time, we generate cell type predictions using scPRINT-2 and achieved a prediction accuracy of 42.2% across the 500 potential cell type leaf labels scPRINT-2 knows about. While this score may seem low compared to supervised approaches, it is worth noting that labels from a secondary source were available in the datasets. Comparing them to the initial ground truth, we found only a 55.3% agreement between the two. Furthermore, we noticed that for some cells, annotations were quite different, such as: *fibroblast* being labelled as *ciliated cell*, *macrophage* as *neuroendocrine cell, and ionocyte* as *secretory cell*.

Given the low correspondence between the two expert annotations, we wanted to determine which was correct between scPRINT-2 zero-shot or the expert ground-truth labels. We conducted a differential expression analysis between cells labeled as *type 2 pneumocyte* by scPRINT-2 (zero-shot) but as *macrophage* by the ground truth (see supp. Figure S6). We saw that the most highly differentially expressed genes were *MAGI1*, *NPNT*, *TEAD1*, and *LMO7*, which are involved in cell-cell junctions, epithelial cells, alveolar cells, and lung tissues. Moreover, the first differentially expressed gene was *SFTPC*, a known “type 2 pneumocyte” marker. This means that, even in this challenging unseen-organism dataset, scPRINT-2 seems to legitimately correct expert annotations. This showcases strong generalization to unseen organisms.

To further improve scPRINT-2’s accuracy, we use a method first presented in Hu et al. to aggregate predictions based on **nearest neighbor smoothing** of the model’s class logits (see <u>Methods</u>)^47,48^. This approach increased accuracy in most of our use cases but yielded a small 1.3% improvement here. We also provide tools to perform **top-K predictions** and **confidence-based selection**. This means that scPRINT-2 can list multiple putative labels for each cell. When multiple labels have high logits, it can output their shared parental label for that cell instead. When labels disagree, or the logits are low, scPRINT-2 can output an “unknown” label instead. Using both approaches together, we get an additional 3% improvement in accuracy, with 10% of the cells now listed as “unknown”.

Additionally, the low accuracy is also related to scPRINT-2 predictions being cell-specific, whereas most ground truth labels are cluster-specific. We propose a **cluster-based logits averaging,** which can be viewed as an extreme case of smoothing (see <u>Methods</u>). With this tool, scPRINT-2 performance increased by 12% (see Figure 2C). Beyond improved accuracy, these inference-time contributions significantly enhance the usefulness of scFM-based cell annotation for biologists.

Finally, we also demonstrate that with our XPEFT method (presented further in Results section 4), scPRINT-2 can improve its predictions to 95% accuracy in the test subset, while preserving some fine-grained cell-type distinctions not present in the training data (see Figure 2C).

We then assessed scPRINT-2’s performance as we increased the number of genes in context. We used a Smart-seq-v4 dataset from Jorstad et al., averaging around 6000 nnz genes per cell (see supp. Figure S7)^49^. As shown in Figure 2D, we observed an overall increase in prediction accuracy across all labels as we increased the context from 200 to 8000 genes, even though scPRINT-2 was pre-trained on only 3200 genes, demonstrating generalization to larger input contexts. Interestingly, classes such as sex and ethnicity reached much better predictive accuracy as we increased the number of genes. When using only the most expressed genes in context, we observed that cell types, which are often defined by highly expressed canonical genes, remained relatively high, even with only 200 genes in context (see supp. Figure S8).

Training scFMs on large dataset sizes does not necessarily improve the model’s performance. It is the breadth of cell types, conditions, organisms, and cell quality that produces real generalization abilities. We showcased it here, with scPRINT2 able to label unseen organisms, improving its predictions across various context lengths and rare modalities. We also showed scPRINT-2 reaching state-of-the-art classification accuracy with our fine-tuning.

We will now see how some of our contributions in training loss and data augmentation can similarly improve performance in denoising and imputation in unseen modalities.

### A multi-cell denoising auto-encoder task unlocks new modalities and performances

Not all single-cell datasets are at the sequencing depth and quality of Smart-seq-v4. On average, single-cell data has very low depth, preventing scFMs from learning features that may only be seen in higher-quality cellular profiles.

### Meta-cells and graph neural network encoder

In addition to biasing sampling toward cells with more non-zero genes (nnz), scPRINT-2’s dataloader now uses neighborhood information, whether defined in expression space or via spatial transcriptomics (see figure 3A; see <u>Methods</u>). This allows users to create models that take into account nearest neighbor cells during pre-training. This can be done, for example, by creating **meta-cells**. Meta-cells average the expression over the cell and its neighbors to artificially create a higher-depth cell with less dropout. We demonstrate that this approach achieves improved results across multiple model metrics, but not in denoising (see Table 1). While 17% of cells in the dataset have more than 2600 non-zero values, 11% had at least 3200. With nnz-weighted sampling, we reach 33%. By adding metacells, half of our input expression profiles now have more than 3200 nnz elements—allowing us to extend scPRINT-2’s context to 3200 genes.

However, one can go beyond meta-cells and, instead of averaging, use a graph neural network (**GNN**) (see Figure 3A; see <u>Methods</u>)^50,51^. In this case, the set of neighbors’ expressions is encoded in the input token of the transformer. We show that this improves the model’s denoising ability. However, we also noticed a decrease in cell embedding and classification (see Table 1). Further experiments showed that this was mitigated with longer training time. As in the variable context case, we variably select 0 to 6 neighbors per minibatch, so the model learns to use a variable number of cell neighbors (see supp. Figure S9, see <u>Methods</u> for details on the choice of neighbors).

**Figure 3:**
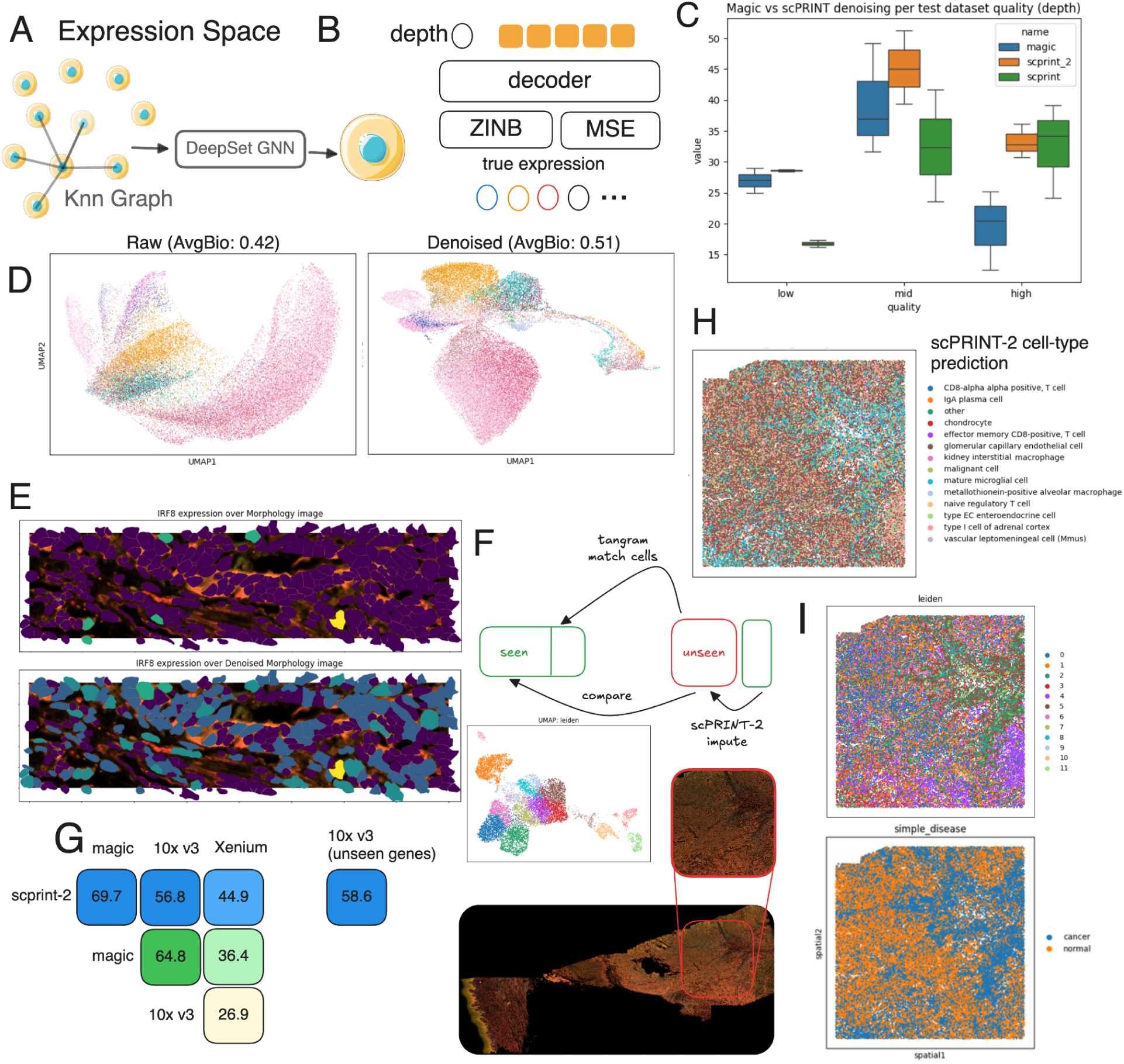
Presentation of the expression encoder and decoders and performance on denoising and imputation tasks. (a) Overview of scPRINT-2’s multi-cell expression encoder and (b) scPRINT-2’s expression decoder loss. Circles represent scalar values, orange blocks represent vectors. (c) Benchmark of scPRINT-2 on expression denoising over nine datasets of varying quality, compared to MAGIC and scPRINT-1. (d) Umap of the Xenium’s patches of cells’ expression pre/post denoising with scPRINT-2. (e) Expression denoising of IRF8 with scPRINT-2 over a sub-patch of the Xenium melanoma dataset with cell contour overlaid. (f) Overview of the patch selection in the Xenium dataset, and of the mapping and pseudo-imputation with Tangram using a matched melanoma 10x v3 single-cell RNA-seq dataset. (g) Correlation-based denoising & imputation scores of scPRINT-2 and denoising of MAGIC on the matched dataset. (f) scPRINT-2 cell type prediction over the Xenium melanoma patch. (i) Expression-based clusters and scPRINT-2 disease prediction of cells from the Xenium melanoma patch analyzed. Source data are provided as a Source Data file.

Pushing our analysis further, we realized that a mix of both scores, which we call **ZiNB+MSE** (see Figure 3B; see <u>Methods</u>), yields a better denoising score while retaining the ability to model zero inflation and uncertainty (see Table 1). Together, these updates have already made scPRINT-2 better than scPRINT-1 and even better than MAGIC on our denoising benchmarks (see Table 1, see Figure 3C)^52^. While these results are already state-of-the-art, we wanted to explore the effects of denoising and how to assess our model in unseen contexts.

Looking at denoising scores across technologies, we notice that scPRINT-1 tends to perform much better on datasets with higher nnz genes, i.e., higher-quality datasets (see Figure 3C, see <u>Methods</u>). However, within each dataset, scPRINT-1 struggles more with low-depth cells than MAGIC & scPRINT-2, which is more consistent overall. We explain this paradox by the fact that, beyond nnz genes, the high-quality dataset often exhibits lower biases in the distribution of nnz genes per cell (see supp. Figure S10). This also explains why MAGIC and scPRINT-2 perform better than scPRINT-1 in these biased datasets. Indeed, they can look at the neighbor’s expression and model the expression biases this way. This usage explains the significant improvements in the low- and mid-quality datasets, making scPRINT-2 state-of-the-art across all tested contexts and modalities using its estimate of zero-inflation.

### scPRINT-2 generalizes to unseen denoising tasks

Additionally, we decided to look at performance on a Xenium dataset, a modality completely absent from scPRINT-2’s training (see <u>Methods</u>)^53^. We elected to use a large, recent skin melanoma dataset with a 5000-gene panel, reaching the upper limit of what is doable with current technology.

A first proof of scPRINT-2’s denoising is the scIB biological truthfulness of the Xenium dataset, which improves over the raw expression embedding when using its embeddings (see Figure 3D, 3E; see supp. Figure S11; see supp. Table S2). To further assess how well scPRINT-2 can denoise this unseen data modality, we leverage the optimal transport-based method Tangram^54^. We used Tangram to map each Xenium cell to another cell in a non-spatial 10X v3 dataset of similar skin melanoma^55^ (see Figure 3F). Here, the mapping quality is low due to many differences between the two technologies, e.g., number of cells, number of genes per cell, or biases in cell and gene types (see supp. figure S12). Still, using the 10X v3 dataset as ground truth, we can see that MAGIC and scPRINT-2 recreate an expression profile that correlates more than 30% better with the 10X dataset than does Xenium (see Figure 3G). There, MAGIC creates expression profiles closer to the 10X ones, while scPRINT-2 remains closer to the initial Xenium profiles, and both scPRINT-2 and MAGIC tend to agree more with each other than with anything else (see Figure 3G). Overall, this suggests that using a tool like scPRINT-2 might be a better alternative for denoising and imputing expression from Xenium than using a secondary non-spatial 10X dataset and aligning it with Tangram.

At the same time, MAGIC can only perform denoising and cannot impute expression for unseen genes. We thus use scPRINT-2 to impute a random subset of 5000 genes present only in the 10X v3 dataset. Interestingly, we noticed that feeding all 5000 (expressed in Xenium) + 5000 (unexpressed in Xenium) genes in context did not lead to good imputation. However, using scPRINT-2’s generative architecture, we directly decoded the 5000 10X-only genes from the scPRINT-2’s cell tokens generated on the 5000 Xenium genes (see supp. Figure S13). We show that this imputation scores as high as the denoised Xenium genes (see Figure 3G).

Finally, we also wanted to examine scPRINT-2’s cell-label predictions on this unseen modality. While we did not have access to ground-truth labels in this dataset, we could already spot-check the validity of the predictions. Indeed, many cell types were labeled as *basal* or *epidermis*, with numerous immune cell labels in the cancer-induced lesion in the tissue (see Figure 3H). This entire lesion region was labeled as *cancer* by scPRINT-2. This was striking as it contained mostly non-cancerous activated immune cells (see Figure 3I; see supp. Figure S14). It likely reflects the biases of the pre-training dataset, where disease labels are often applied at the dataset level rather than the cell level, making scPRINT-2’s disease predictions sometimes imprecise. Thankfully, many cells had the cell-type label ‘*malignant cell*’. These cells were distributed throughout the tissue and showed a strong signal for the five key literature melanoma genes (*BCL2, IGF1, EGFR, FGFR2, SOX10*) (see supp. Figures S15 and S16).

Overall, we have seen how scPRINT-2 can be used on challenging modalities to augment a given dataset with cell label predictions, expression denoising, and gene imputation. Showing yet again another axis of generalization. We will now focus on how structural changes to scPRINT-2’s transformer architecture improve the quality of its embeddings.

### An efficient, hierarchical attention architecture makes scPRINT-2 generative

#### Efficient attention architectures and compression methods

Implementing transformer models on new modalities is a potent way to rethink some of their mechanisms. A common issue with transformer models is their memory and compute requirements, which grow quadratically with their context length (e.g., the number of genes in their input). This is even more pronounced in bidirectional transformers like most scFMs. With the introduction of scPRINT-1, we presented a model that could train in 3 days on a regular A40 GPU and on 50M cells, an order of magnitude faster than most similar scFMs. A first contribution to the scPRINT-2 architecture is the addition of state-of-the-art approaches to reduce the memory footprint and increase training speed. We modified the attention mechanism in multiple ways, using grouped-query attention (GQA) to reduce memory usage. We benchmarked additional attention mechanisms alongside Flash-Attention-3 to assess their performance and their speed.

Over-representation plot of the top positively differentially expressed genes in both human-like mouse and real human vs. mouse; the red line indicates random chance. Source data are provided as a Source Data file.

A first one is flash-**hyper attention**, which computes specific attention only on sets of keys and queries known to be similar via locality-sensitive hashing and clustering^56^. A second one is flash-**softpick attention**, a rectified softmax that decreases hyperactivation of specific tokens, often called attention sinks^57^. We also present our own sub-quadratic attention mechanism: **criss-cross attention** (see <u>Methods</u>), inspired by advanced concepts such as the Recurrent Interface Network (RIN) and the Induced Set Attention Block (ISAB)^58,59^. It compresses attention by sketching it in context, using a doubly cross-attention mechanism with a set of latent tokens that get updated across layers (see supp Figure S17). We show that only criss-cross attention dramatically improved the model’s speed while retaining all its abilities (see Table 1). However, it is not yet compatible to retrieve gene networks from; for this reason, our scPRINT-2 architecture, for now, uses flash-attention-3 and XPressor.

On another direction, while single-cell analysis has leveraged VAEs for years to generate meaningful compressed representations of cells, transformers inherently lack this ability^60–62^. We use the **XPressor** architecture presented in Kalfon et al.^33^, which compresses output gene embeddings into a set of cell embeddings and decompresses them back into their original gene embeddings (see figure 1C, figure 4A, and <u>Methods</u>). This innovative architecture draws on ideas that have existed in the transformer literature for several years^59,63–66^. We show in our ablation study that using XPressor results in a slightly better cell representation overall, but does not meet the statistical threshold. This difference might be explained by the limit in the number of epochs and the model’s smaller size compared to Kalfon et al. (see Table 1, see supp. Table S1). We include an extension to this approach, in which one appends VAEs to each output embedding of XPressor to regularise the different cell embeddings generated by the model (see Figure 4B). This addition allows us to choose a specific dimension for each cell embedding that is lower than that of XPressor. A second constraint is defined by applying the Kullback-Leibler divergence (KL) loss (see figure 1C, see <u>Methods</u>). This creates an information bottleneck for the different cell embeddings, pushing the model to select only the minimum amount of relevant information to represent the label. While our ablation study does not show improvement in cell embeddings with this approach, this is likely because each method was trained for only 20 epochs. Indeed, the VAE-infused model is taking longer to learn to classify cells. However, the batch correction score improved significantly, indicating that the different cell tokens mainly contained information about the class they encoded (see supp. Table S2). Now that we have highly compressed cell-level embeddings (i.e., tokens), we can apply a **dissimilarity loss** between each for a given cell. This actively pushes them to be as different as possible (see supp figure S18; see <u>Methods</u>). We demonstrate that this tends to slightly improve the model’s output embedding in our ablation study (see Table 1).

**Figure 4:**
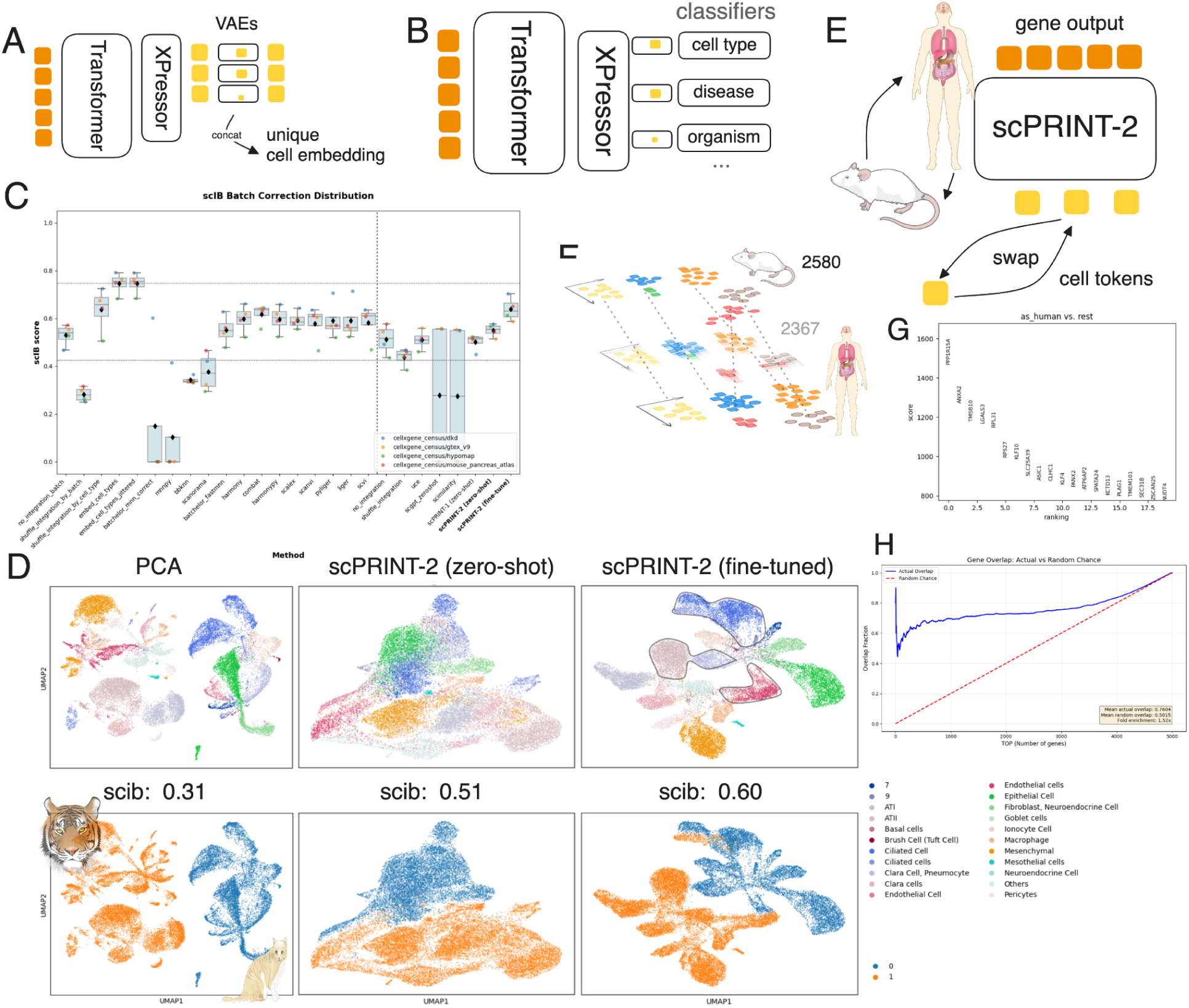
Presentation of the XPressor architecture and performance on cell embedding tasks: (a) Presentation of the XPressor with VAE-based compression. (b) Schematic representation of going from expression to classification with scPRINT-2, XPressor, and VAE-based compression. (c) Open-Problem scores for scPRINT-2 across all methods. (d) Umaps of, respectively, PCA embeddings, scPRINT-2 zero-shot cell-type embeddings, and scPRINT-2 fine-tuned cell-type embedding colors by known cell types and batches, with scib total scores. overlaid in the upper-right figure are cell types that have different labels but are actually the same (see supp Figure (e) schematic representation of the counterfactual generation using scPRINT-2’s embedding and replacing them for the organism class from mouse to human. (f) Illustration of the decrease in distance between initially unrelated datasets from applying this counterfactual approach. (g) differentially expressed genes post vs pre mouse “humanization” with scPRINT-2. (h)

These architectural changes make scPRINT-2 much more efficient at compression and zero-shot batch-correction. Indeed, on the open problem’s benchmark, we observe an overall improvement over scPRINT-1, again becoming the state-of-the-art zero-shot method on the platform (see Figure 4C). This zero-shot performance increase is solely due to the improvement in the batch-correction score from using our VAE method (see supp figure S19). We then fine-tune the XPressor architecture alone – our XPEFT approach – to further learn to remove batch effects and predict expert-annotated cell-type labels. We add a Maximum Mean Discrepancy (MMD) loss (see <u>Methods</u>) that penalizes the distance between batch elements^67,68^. Doing so, we observe a jump in scIB scores, especially in biological truthfulness, as measured by the scIB metrics (see Figure 4C; supp Figure S20), making scPRINT-2 the best-performing method in the benchmark.

### scPRINT-2 generalizes to unseen cell embedding tasks

We then wanted to push our analysis further and test the zero-shot organism-level integration of scPRINT-2 on organisms unseen during training. Again, using our cat and tiger dataset presented in the <u>second result section</u>, we saw that already, scPRINT-2’s general cell embedding performs better than doing no correction and keeps lot of biological truthfulness, as shown by the scIB score of 0.44 vs 0.37 for PCA (see Figure 4D, see supp. Figures S21, S22, see supp. Table S4). Then, as often, taking the cell-type-specific embedding further increases the biological truthfulness to 0.49, mainly by generating a more faithful biological representation, as reflected in the scIB scores (see Figure 4D, supp. Figures S22, S23, supp. Table S4). Again, using XPEFT, we achieve a tremendous 0.60 scIB score, placing us among the top 3 best-performing models in this category, behind SATURN and scGEN (see Figure 3D, supp. Figure S22, S24, supp. Table S3). We note that even in this domain, many cell types didn’t overlap across organisms. It is a common behavior in this benchmark, and similar cell types now almost overlap in the UMAP, hinting at shared neighbors (see Figure 3D, see <u>Methods</u>)^69^.

Finally, we wanted to examine the model’s ability not only to integrate cellular profiles but also to generate entirely new ones at inference time in a zero-shot manner by combining cell tokens (see Figure 4E). We first approach it using a matched mouse-human multi-organ atlas from Zhong et al.^46^. We then generated cell embeddings for all cells and computed an average “human”-ness cell embedding using the *organism* embeddings of all human cells. We regenerate an expression profile using 1. the human gene embedding and 2. the mouse cell embeddings, replacing the organism cell embedding with the human one (see Figure 4E and <u>Methods</u>). We thus generate a set of human-like cell expression profiles from mouse expression profiles. Using the 5000 most variable orthologous genes, we indeed observed a decrease in the Wasserstein-2 (W2) distance on this counterfactual conversion to human (see Figure 4F, see <u>Methods</u>)^70,71^.

Applying a similar approach, but this time to generate females from males in the human dataset, we also notice a similar reduction in expression W2-distance from 1076 to 938.

Looking at how cell expression patterns change after this transition, we found that most of the top differentially expressed genes are the same as those identified in the differential expression analysis of the real human dataset (see Figure 4G, see supp. Figure S25). Computing an over-representation test, we observe a robust 58% enrichment compared to random, with more than half of the top differentially expressed genes correctly predicted by scPRINT-2 in both over-and under-expressed genes (see Figure 4H, see supp. Figures S26, S27). Looking at *Reactome_2022* pathway enrichments, we see multiple pathways related to immune system function, membrane-ECM (Extra-Cellular Matrix) interactions, and tissue elasticity, as well as many other molecular-level pathways (see supp. Figure S28). These align with previous analyses highlighting ECM and immune function differences between human and mouse tissues^72,73^.

Overall, we have shown that an entirely novel architecture and a set of learning constraints enable scPRINT-2 to generate high-quality embeddings in a zero-shot manner. Thanks to its multi-organism training, this can be extended to unseen species, while achieving even stronger results with fine-tuning. We have also demonstrated how one can use the scPRINT-2’s cell embeddings to generate counterfactual cellular profiles. This makes it a strong contender for performing atlas-scale analysis across tissues, diseases, and organisms, by learning to disentangle each cell component. We will now see how other parts of the models can be used to extract additional information.

### High-quality contextual gene representations from scPRINT-2

#### scPRINT-2 has rich gene embeddings

scFMs don’t just provide cell-level embedding, they have also been used to generate contextual gene-level embeddings given a cell’s expression profile or to predict gene-gene connections.

The model’s gene embeddings can be used for fine-tuning, such as to predict ATAC-seq activities or gene essentiality^1,2^. We investigate the gene embeddings produced by scPRINT-2 and then delve into how its gene networks can be better extracted and assessed.

A good output gene embedding is also defined by the quality of its input. With scPRINT-2, we introduced a fine-tuning adapter layer on top of ESM3’s protein embeddings, jointly trained with the model (see <u>Methods</u>). This approach is one of the few that improve gene network inference without decreasing any other metrics in our additive benchmark (see Table 1). It allows us to update gene representations during pre-training while maintaining the ability to work with unseen representations, e.g., from unseen species (see Figure 5A).

**Figure 5:**
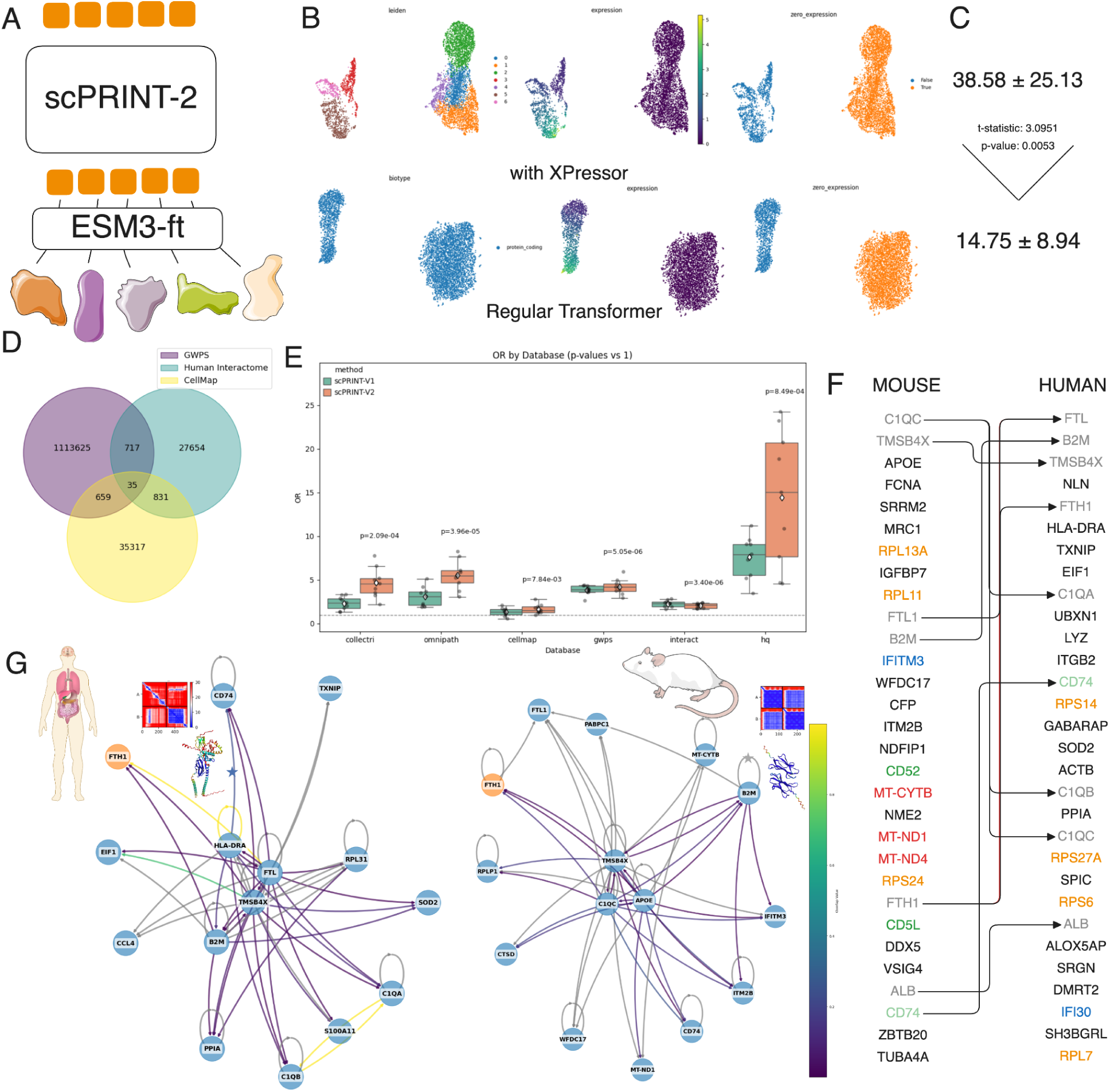
Presentation of the ESM3 fine-tuning and gene network study. (a) Illustration of fine-tuning of ESM3 while training scPRINT-2 using an adaptor layer. (b) Comparison of gene output-embeddings for a random cell in a model with the XPressor architecture and a model trained without. (c) On the side, the average number of pathways shown to be enriched in the gene output embedding clustering of each method using three main pathway databases. The number below is on the non-fully trained regular transformer; otherwise, no pathways are enriched (see <u>Methods</u>). (d) Comparison of ground truth networks’ overlap between cellmap, the human interactome, and genome-wide perturb-seq. (e) Benchmark over six ground truth gene networks of scPRINT-1’s gene networks with its extraction method and scPRINT-2’s gene networks with its extraction method, over nine different human cell types from the same dataset. (f) Comparison of the top-30 hub nodes on both gene networks. Arrows link similar genes, and colors represent similar gene groups. (g) Subset of a gene network generated by scPRINT-2 seeded at FTL1, on human macrophage cells, and on mouse macrophage cells, edge color represents the RoseTTAFold2-PPI scores for these connections, grey means no score was computed. The AlphaFold-Multimer structure and amino-acid distance map are provided for the star-marked connections. Source data are provided as a Source Data file.

It remains unclear, however, what the right approach is for selecting output gene embeddings, with some heuristics proposing using the last or second-to-last layer. Using our regular transformer model trained with masking, we demonstrate that its output gene embeddings contain only their own expression values (see Figure 5B). However, when trained with the Xpressor architecture, clusters of genes appear (see Figure 5B). This is sensible because Xpressor forces gene embeddings to be rich in meaning, as the compression block must query them. We have, however, noticed that for regular models that are not fully trained (only up to 20 epochs), the output gene embedding still contains some input ESM3 features (see supp. Figure S29). The number of enriched pathways in its output gene embedding cluster is still significantly less than for scPRINT-2’s XPressor architecture (see Figure 5C; see <u>Methods</u>)

### extracting gene networks from scPRINT-2

Thanks to the transformer architecture, one can go beyond gene output embeddings to examine gene-gene interactions via the model’s attention layers. Following the tests reported in Kalfon et al., we observed, on average, no dramatic performance gains across the methods we tested (see Table 1). An issue we noticed is that the problem is not well-defined. Indeed, the ground truths widely disagreed with one another (see Figure 5D; see <u>Methods</u>). Between the genome-wide perturb-seq (gwps) ground truth and omnipath, only 800 gene-gene connections were in common over the hundreds of thousands that each contained. This suggests that diversity of ground truth will be key to showcasing the breadth of potential gene-gene connections in the cell.

We thus gathered a new set of ground truth gene networks (GN)s from recent works. Our first approach was to use protein-binding datafrom AP-MS experiments within the O2US cell line, called the *cellmap*^74^ (see <u>Methods</u>). Additionally, thanks to protein structure models, we are now able to compute putative interactions across millions of protein pairs; a first version of this analysis has been defined in the human *interactome* (see <u>Methods</u>). But here again, the disagreement was significant, with only ∼1-4% of the connections in each ground truth being found in another, and no connections were reliably found across all five ground truths (see supp. Figure S30).

Acknowledging these disagreements, we benchmarked them against nine human cell types from the same dataset using scPRINT-1 and scPRINT-2. We use a gene network extraction method that is more computationally demanding but biases the network towards co-expressed genes (see <u>Methods</u>). We see that scPRINT-2’s performance was often greater or similar across all benchmark networks, as indicated by the odds-ratio measures (see Figure 5E; see <u>Methods</u>). We did not see a similar trend, however, on AUPRC (see supp. Figure S31). This suggests that our method is more accurate for its top-K connections. Indeed, the strongest human interactome connections were overrepresented in scPRINT-2, more so than in scPRINT-1.

### cross-organism gene network analysis

To continue on our cross-organism analysis, we also aimed to further characterize some of the genes observed in our previous human/mouse datasets by interrogating the cell-specific GN identified by scPRINT-2 in *Macrophage* cells from both mammals. Looking at their hub nodes, we see that many are common and represent key conserved cell immune pathways, such as *feroptosis*, *vitamin B12*, and *Pathogen Phagocytosis Pathways* (*WikiPathway_2023_Human*), with genes like *C1Qs, RPs, ALB, and APOE*^75,76^. These mainly relate to the macrophage’s internal machinery, which is designed to eat and destroy pathogens. Other genes were clear markers of macrophages (*CD74;LYZ) and/or immune cells (HLA-DRA, B2M) or their pathways, such as interferons alpha/gamma and MHC Class II (MSigDB_Hallmark_202076–78)*. Interestingly, these networks share only 30% similarity when considering the top 20 connections for each gene. But what seemed like differences in connections and top 50 hub genes tended to disappear after thorough analysis, such as with the Ribosomal proteins, which are related in the kinds of pathways they are part of, or in their relationships in the PPI_Hub_Proteins database^77^ (see Figure 5F).

We then extracted a subset of the macrophage networks, seeded at the *FTH1* gene, for both organisms, focusing on the top 15 connected nodes and their top 60 edges (see Figure 5G; see <u>Methods</u>). We observed a set of hub genes in both subnetworks, with some genes being shared between human and mouse. Interestingly, these hub genes had more interactions in the human interactome ground truth than non-hub genes. We also noticed that the “hub-ness” of the subnetworks can be very variable and seems to depend on the “seed” gene (see supp. Figure S32).

By overlaying the human interactome ground-truth values on our subnetworks, we found that only a small subset of connections was marked as valid (i.e., score above 0.6) in the ground truth (see Figure 5G). In the mouse *Macrophage* subnetwork, almost no connections were recovered, but this may be explained by the fact that the ground truth is the “human” interactome, computed using human proteins rather than mouse proteins. We thus wondered whether we could use scPRINT-2 to cross-validate the interactions present in this ground truth. Indeed, we know that the human interactome values are not directly computed from AlphaFold-multimer’s interaction probability (ipTM); they come from a simpler model called “RoseTTAFold2-PPI”. Testing a couple of connections predicted to be low ipTM by RoseTTAFold2-PPI but found by scPRINT-2, we readily identified two: HLA-DRA/CD74 and B2M/B2M, which, when passed to AlphaFold-Multimer, indeed formed an interaction with an ipTM of more than 0.6. This showcases the potential of scPRINT-2 in this domain and future directions for GN inference.

We have seen here how scPRINT-2’s output gene embeddings and attention matrices can be used to extract meaningful biological insights and drive hypothesis generation in a cell-to-cell, state-specific manner. These outputs can also be used for fine-tuning purposes and in explainable AI-driven analysis. We also pushed our GN analysis further, defining additional benchmarks and a more powerful GN extraction mechanism. We demonstrated cross-species analysis and presented the tantalizing possibility of merging foundation models at different scales, including ESM3 fine-tuning, AlphaFold Multimer, RoseTTAFold2-PPI, and scPRINT-2.

While these are just examples, they demonstrate what aggregating multiple bodies of evidence across scales can achieve for genetic interaction predictions. A first step towards using scFMs, protein Language Models, and structural models in coordination, to shed light on the cellular machinery.

## Discussion

In this work, we present a gymnasium of tasks to benchmark scFMs in multiple contexts. Together with an efficient and reproducible pipeline, we test the benefits of 42 different parts of scFMs structures, encoding, and training. In this additive benchmark, 12 of these are our own contributions to scFMs, including GNN-based expression encoding, cross-foundation model fine-tuning, sub-quadratic attention mechanisms, and rich losses. This massive benchmark is the first of its kind for scFMs and assesses four different tasks. It allowed us to identify bottlenecks and limitations, issues that we solved in subsequent analysis. Indeed, future benchmarks will benefit from using more diverse datasets, tasks, and ground truths.

We have also presented the largest pre-training database to date, encompassing more organisms, conditions, and data modalities. We have seen that, while more work is needed to obtain higher-quality, well-annotated datasets, our dataloader and preprocessing pipeline have made the most of this vast database.

Using the best feature combinations from our additive benchmarking, we build and train a next-generation cell Foundation Model, scPRINT-2. We demonstrate that, although currently 5 times smaller, scPRINT-2 outperforms scPRINT-1 across all benchmarks tested. On denoising, scPRINT-2 becomes state-of-the-art, and with our fine-tuning approach, it also outperforms every other method on the batch-correction and classification tasks of the open-problem benchmarks.

We then challenge scPRINT-2 on tasks of high relevance for cellular biology, highlighting some pitfalls in current benchmarks. We show that scPRINT-2 acquires generalizable abilities across unseen modalities and organisms, while remaining consistent in its predictions. We demonstrate it across many tasks, including cross-organism integration, unseen gene imputation, and counterfactual reasoning.

Finally, we present tools for easily extracting labels, cell-specific gene embeddings, imputing gene expression, performing gene network inference, and working with organisms unseen during pre-training. We believe our results demonstrate many domains where scFMs might confidently replace approaches that rely on heuristics, atlases, and a variety of tools and packages. However, much work remains.

Current ground-truth cell annotations are cluster-based and obfuscate the complexity of cellular states by inherent clustering biases. Batch correction metrics are similarly biased, and top scores can be easily gamed; gene network ground-truths are not cell type specific and likely filled with false negatives. Data diversity and quality are the principal pre-training bottlenecks, and efforts will be needed to improve foundation models. Many other key modalities, such as measuring time and perturbation effects, remain scarce. They will become increasingly helpful for enriching the future comprehensive benchmarks of next-generation cell foundation models.

Our analysis and contributions highlight powerful features of scFMs and provide guidance for designing benchmarks that better highlight their strengths and weaknesses. scPRINT-2 presents a direction for future improvements, with more specialized architectures and using a combination of biological FMs working jointly across modalities and scales. This next-generation scFM is a step forward in the design of AI for cell biology.

## Methods

We present an additive benchmark with over a dozen contributions to the pre-training tasks, losses, and architecture of single-cell foundation models. Along with it, **scPRINT-2** (pronounced “sprint”), a next-generation model trained on the best-performing contributions. We analyze its out-of-distribution generalization and present methods for querying and fine-tuning it to solve various tasks. We will go through the specific techniques that made it possible.

### Additive benchmark

We now describe in matched order with respect to Table 1, the methods behind the multiple contributions tested in our additive benchmark (see <u>Results</u> section 1). We bolded the ones that are further defined in the methods. In this benchmark, we are using and testing the:

1. **“base model”,** every subsequent element is applied to the base model
2. “medium model”, larger base model, see the <u>base model section</u>
3. “negative control”, untrained base model Architecture
4. “no dropout”, where we remove the dropout initially applied in the base model
5. “large classifier”, where the classifier sizes are increased from [input - output] in the base model to [input - 256 - output]
6. “MVC”, where we replace the base model’s decoder with the cell embedding’s MVC approach of scGPT^1^
7. “no decoders/generation”, where we removed the base model’s decoder, getting a masking+classification only pre-training Data
8. replacing our pre-training dataset with “Tahoe”’s 100M dataset
9. Chan Zuckerberg Institute (“CZI”)’s cellxgene database (version 2024)
10. replacing our pretraining dataset with “CZI + Tahoe” with Tahoe’s 100M database
11. replacing our pretraining dataset with “all databases”, both CZI, Tahor, and Arc’s scBasecount^26,27^
12. replacing our pretraining dataset with “only 200 random” human datasets
13. replacing our sampling with a “sampling without replacement”
14. replacing our sampling with “cluster-based sampling only”
15. **adding “meta-cell” during pre-training** Attention
16. replacing FA3 with “softpick” attention, using the approach of Zuhri et al.^57^
17. replacing FA3 with “hyper”-attention, using the approach of Han et al.^56^
18. replacing self-attention with **“criss-cross” attention layers**
19. adding “XPressor” layers Losses
20. **adding “contrastive learning”**
21. **adding “elastic cell similarity”**
22. **“no embedding independence loss”, removing the embedding independence loss**
23. replacing the ZiNB loss with Mean Squared Error (“MSE”)-loss
24. **replacing the ZiNB loss with “ZINB+MSE” loss**
25. adding a “VAE compressor” loss to the Base model Tasks
26. adding “variable context length” and a larger context
27. **replacing masking with a Transcription Factor “(TF)-masking” task**
28. replacing masking with “denoising”, using the approach in scPRINT, with a random level of denoising (see below)
29. “no classification”, removing the classification pre-training task
30. adding an “adversarial classifier” Input
31. replacing log1p normalization with “sum normalization” where each expression profile is normalized to sum to 10,000
32. “no random level of denoising” where we remove the random level of denoising, see the <u>denoising section</u>
33. where we replace the expression encoder with a Graph Neural Network (“GNN”) encoder
34. where we replace the continuous expression encoder with a “binning” version, following the approach of scGPT^1^
35. where we are “using only expressed genes”, as in scGPT and geneformer
36. “without using gene location”, removing the gene location information in the input tokens.
37. “learn gene embedding” where we replace the ESM3 gene embedding with learnt embeddings, as in scGPT and Geneformer.
38. replacing the ESM3 gene encoder with a “fine-tuned ESM3” gene encoder

The full training traces of the entire additive benchmark are available on weights and biases: https://wandb.ai/ml4ig/scprint_ablation/reports/scPRINT-2-additive-benchmark--Vmlldz oxNTIyOTYwNA?accessToken=0mzwwu64py309mds6zzbgcxllrgcdnd10laivhs3ykh9pq mbs0wxutcu60py2bld

#### Base model and training

The additive benchmark is performed on a small model with 18.2M parameters, an embedding dimension of 256, and 8 layers and 4 heads. The model trains for 20 epochs of 20,000 batches of 64 cells per batch. Validation is performed on 10,000 minibatches. We otherwise use the same optimizer and hyperparameters as for scPRINT-2 (see **pre-training** in <u>Methods</u>)

Gene expression is encoded using ESM3 embedding, with gene location and MLP-based expression encoding added, as described by Kalfon et al. The output is decoded using an MLP that takes the output embeddings and depth information, then outputs a scalar expression value.

The base model is trained on CZI’s cellxgene census dataset, version 2024 (compared to 2022 in Kalfon et al.). The pre-training task uses a 30% gene expression mask with an MSE loss (as is common for BERT-like encoder transformers)^1,2,7^. The Base model also uses a multi-cell-token generative loss as described in Kalfon et al.^3^. It also performs matched multi-class hierarchical classification, as defined below (see **Hierarchical classifier** in the <u>Methods</u>). Finally, it also uses a dissimilarity loss between each of our cell embeddings for a given cell (see <u>embedding</u> <u>independence in Methods</u>). Each of these decisions is assessed within our additive study.

We pre-train the base model 6 times across multiple seeds to generate error bounds. We train using Flash-Attention-3 on 1 H100 GPU, each training of 20 epochs taking roughly 2 days. Some runs were done on A100s and V100s; we thus had to rescale the time duration for some of these runs.

Some additive study runs use denoising as a training strategy or larger context lengths when it seemed likely that this would best highlight the abilities and shortcomings of the benchmarked element.

The **medium model** size uses an embedding dimension of 512, with 16 layers and 8 heads. The **negative control** is a model that was not trained at all.

#### Weighted sampling

The goal of weighted random sampling is to de-bias regular random sampling of cells in contexts where many cells have similar profiles and expression patterns, while others are rare cell types.

We use weighted random sampling on our training data based on all the different class values we have to predict. We use a factor of *S*_1_, meaning the rarest elements will, on average, be sampled only *S*_1_ times less than the most common ones. The sampling factor used for each group is then 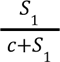, instead of 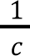, where *c* is the number of cells in each group.

#### Cluster-weighted sampling

The goal of cluster-weighted sampling is to improve weighted sampling in the condition where cell-type annotations are poor or non-existent.

For cluster-weighted sampling, we simply use the labels obtained by applying Leiden clustering to the K-NN graph of cells for each dataset during preprocessing. We used a resolution of 1 and 15 neighbors. We merge clusters if their centroid correlation exceeds a threshold (here 94%). This cluster label is then treated similarly to other labels, such as *cell_type*, *sequencer*, etc.

In this context, within datasets that lack information about tissue of origin or sequencer, or that belong to the same categories, cells from cluster 1 will be sampled with equal weight from those datasets. The sampling is not dataset-specific. This decision arises because most datasets contain some information about their tissue of origin or disease, and cluster sizes of data from the same tissue/disease often represent similar cells. This can be applied to any dataset for training models.

#### Depth-weighted sampling

The goal of depth-weighted sampling is to sample cells with higher quality, in terms of the number of genes expressed, more often.

For depth-weighted sampling, we scale each cell’s sampling probability by its non-zero (nnz) gene count. Similarly, we scale this value, but this time we apply a sigmoid function beforehand to reduce the impact of extreme values.

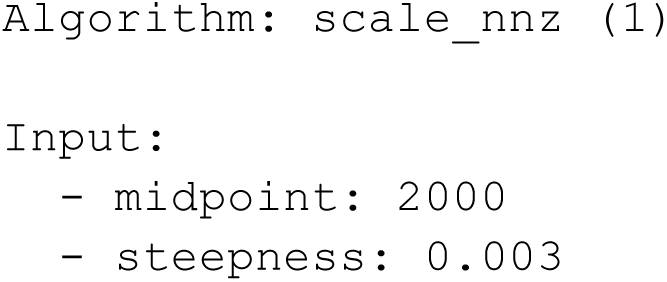
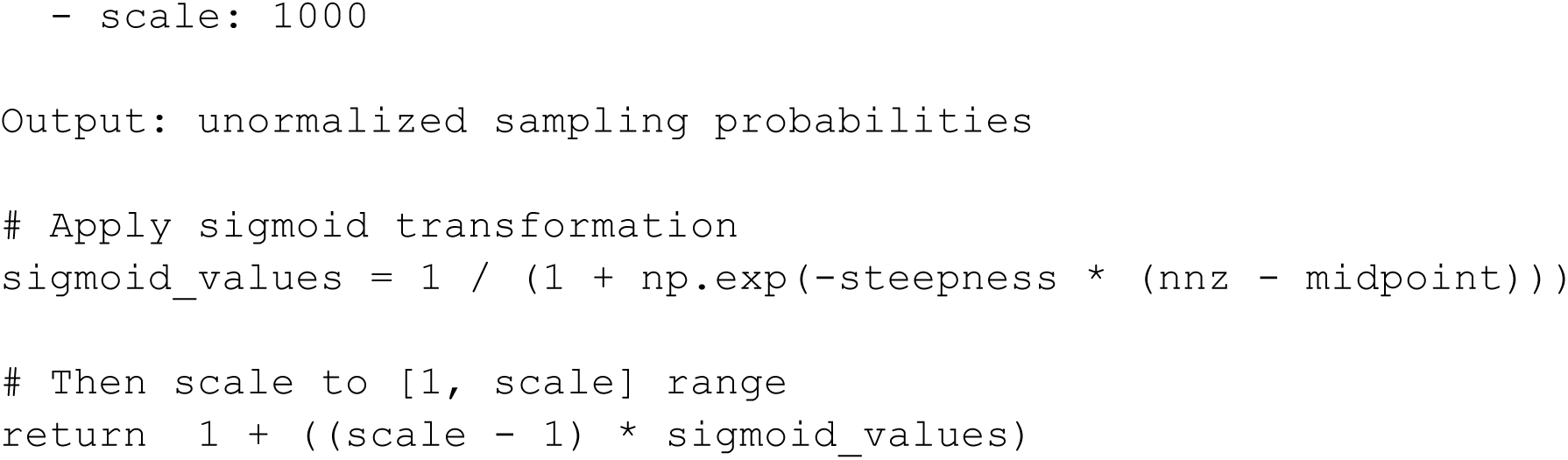

The values shown for Input were the ones we used across our research and were selected manually.

This can be applied to any single-cell dataset for training models.

#### Multi-cell sampling

For all our datasets, our preprocessing pipeline computes a K-NN graph from the PCA of the scaled, log-transformed expression data. For each sampled cell, scDataloader also retrieves its k-NN cell ID and loads them, along with their distance information. Here, we set K to 6 and the PCA components to 200.

We set the number of PCA components to 200 to retain as much information as possible, while accounting for rare cells whose expression might have only a small impact on the first PCA components.

We set K to 6 to balance the computational resources required to sample 6 times more cells per minibatch with the need for enough neighboring cells. Indeed, these computational resources are more prevalent for smaller models that perform fast iterations across many cells than for larger models. 6 neighbors per sampled cell was our limit for a small foundation model like scPRINT-2. We also note that there is likely a rapid diminishing returns beyond 6 to 15 cells for most datasets as we start sampling more often cells that are less similar to the center one. During scPRINT-2 training, we select 0 to 6 neighbors per minibatch, so the model learns to use a variable number of cell neighbors.

#### GNN Expression encoder

The goal of the GNN expression encoder is to increase the information the foundation model can obtain from 1 cell to a set of neighboring cells, thereby dramatically reducing input noise.

The GNN takes multiple expression values as input, optionally along with corresponding cell-cell distances, and returns a vector encoding this information. Both continuous and GNN encodings can be configured to receive either logp1-transformed expression data, sum-normalized expression, or both. The GNN follows the DeepSet^78^ implementation:

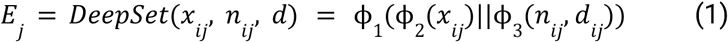

Where:

– *x_j_* is the center cell’s expression for the gene *j*
– *n_ij_* ∈ ℜ^k^ is the K nearest neighbor cell’s expression for gene j and cell i
– *d_ij_* ∈ ℜ^k^ is the distance of each neighboring cell to the center one
– ϕ_i_ are MLPs.
– || is the concat operation

We selected K to be a random number between 0 and 6 during training and 6 at inference.

#### ESM3 fine-tuning gene-encoder

The goal of ESM3 fine-tuning is to get the best of both worlds between learning token features from the data and using learnt protein representations from a pLM as a prior.

We encode/tokenize gene IDs using ESM3^79^. The mapping process happens in the following way:

– A gene name is mapped to its canonical protein name using Ensembl114.
– We recover the protein sequence of the protein using Ensembl
– We use the protein sequence to generate an embedding using ESM3 by averaging all its amino-acid output embeddings.

For the fine-tuning part, we reuse the fine-tuning approaches presented in Kalfon et al., which place an additional adapter layer after mean-pooling and before feeding the protein representation to the model. Interestingly, using gene expression as a further signal to the adaptor layer often led to training instability.

#### Biased attention

The goal of biased attention is to orient our attention matrix towards genetic interaction priors to improve learning and the model’s biological fidelity.

We leveraged the Rcistarget computation and ranking of the human genome for 10kb down- and upstream of each target gene^80^, available at: https://resources.aertslab.org/cistarget/databases/homo_sapiens/hg38/refseq_r80/mc_v10_clust/gene_based/hg38_10kbp_up_10kbp_down_full_tx_v10_clust.genes_vs_motifs.rankings.feather. Using this information, we generate a weight matrix *M* that links each motif-defined TF to its target genes.

Given this matrix, we bias the attention matrix for all heads and layers using the attn_mask parameter of the `torch.nn.functional.scaled_dot_product_attention` function.

It appears in the attention computation like so: 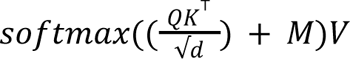 where *M* is the attn_mask matrix and is real-valued.

#### Criss-Cross attention

The goal of criss-cross attention is to create an efficient attention mechanism by learning, in context, a factorisation of each attention matrix.

In criss-cross attention, we replace the self-attention mechanism with a double cross attention between the *N* input elements and *M* latent tokens (see supp. Figure S18). This is thus replacing *N*^2^ computation with a 2*NM* one, hence going below the quadratic bottleneck of attention. This bears resemblance to the ISAB architecture, XPressor, and perceiverIO^58,63–65,81,82^. M, in our case, is set to 10: our 9 predicted classes plus an additional token.

Effectively, the *M* latent tokens are learnt at the first layer of the models. At the same time, they could also be generated from a sketching or principal components analysis (PCA) of the input tokens. They also get updated during the attention computation, so that at the second layer.

We replace the traditional attention computation *X_l+1_ = Attention(X_l_, X_l_, X_l_*) + X_l_, where Attention takes as input the Query, Key, Value elements, with:

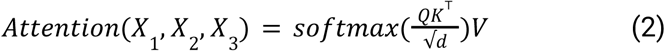

With

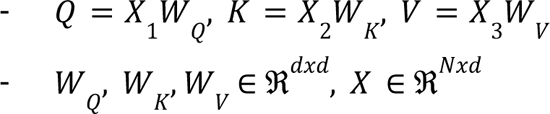

In self-attention *X*_1_ = *X*_2_ = *X*_3_

In Criss-Cross attention, the algorithm becomes:

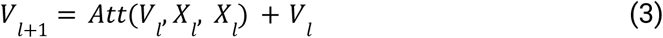

for the latent update and

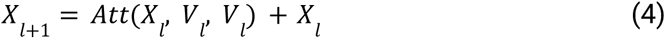

for the main update with *X_l_ ∈ ℜ^N×d^* the main embeddings and *V_l_ ∈ ℜ^M×e^* the latent embeddings

#### XPressor model

The goal of the XPressor architecture, as presented in Kalfon et al., is to replace and generalize the class-pooling of other transformer models and the bottleneck learning of scPRINT. This makes the model more powerful at encoding cell-level features while also separating cell-level tokens from gene-level tokens. Finally, it enables a new mode of Parameter-Efficient

Fine-Tuning. This bears similarities to the ideas presented in criss-cross attention above.

The **Xpressor** block uses as input a set of learned latent tokens *T*. It then performs cross-attention between the last layer of the gene embeddings and the latent tokens. The goal is for the **Xpressor** layers to be of smaller dimensions and context size than the main transformer layers, such that we end up with *C*_*j*_ a set of *n* tokens of dimension *d_t_* generated from the encoded gene expression and ID matrices *E*_*j*_, and *G*. Where *G* and *E*_*j*_ are sets of *m* tokens of size *d_t_*_*c*_ representing the IDs of the genes and their corresponding expression in cell $j$, respectively, where *d_t_*_*c*_ < *d_t_*_t_ and *d* << *m*:

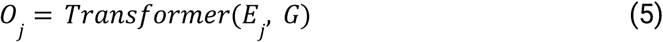

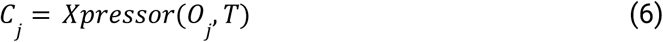

For a cell *j*, with the **Xpressor** being initialized with a learned set of input cell tokens, and *C*_*j*_ being the cell tokens associated with the input *E*_*j*_.

The **Transformer** and **Xpressor** are both transformers with *l*_1_ and *l*_2_ layers, respectively. Indeed, we have designed both layers to contain a cross-attention architecture (see Figure 4A, supp. Figure S1) such that we can also do:

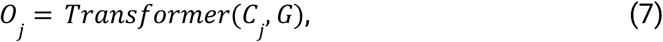

With *O*_*j*_ the output of the **Transformer** when using the **Xpressor** representation as input.

We add an optional MLP after cross-attention to transform the embeddings before the self-attention round. In our example, the decompression is performed using gene ID tokens as input only. These tokens remain the same for all cells of a given organism and thus do not depend on *j*. In the context of protein language models, for example, this would be replaced by positional tokens.

As shown in supplementary figure S1, the **Transformer** blocks are applied twice. The first application serves as an “encoder”, using only self-attention, while the **Xpressor** and the second application of the **Transformer** blocks act as “decoders”. We follow these definitions from the original “Attention is All You Need” paper^83^. It should be noted that, in our case, cross-attention is performed before self-attention.

Related ideas have also been explored in the NVEmbed paper, where the authors propose a cross-attention-based method to update tokens using “latent” tokens and some additional prompting tricks^66^.

XPressor can be applied during pre-training or fine-tuning to replace mean-max-class pooling in Foundation models.

#### VAE-based compressor model

The goal of the VAE-based compressor is to reduce information sharing between output embeddings by penalizing the amount of information each embedding stores (see Figure 4A, supp. Figure S1)^84^.

Each VAE-based compressor is explicitly applied to a cell embedding, compressing it into a relevant latent dimension. It has a 2-layer MLP encoder and a 2-layer MLP decoder. In cases where only a small set of possible elements exists, such as in sex embeddings or cell culture, one can use the Finite Scalar Quantization (FSQ)-VAE^85^.

FSQ-VAE discretizes each latent dimension **independently**. Specifically, the encoder outputs *d_t_* values, each constrained to lie within a bounded range (e.g., [-1, 1]). Each dimension is then quantized into one of *M* discrete levels within that range (in our case 2). This dimension-wise quantization can be implemented as either a hard nearest-bin assignment or a differentiable approximation thereof. Because FSQ enforces scalar-level discretization, it provides a simpler and more fine-grained alternative to VQ’s vector-level codebook approach, while still offering strong regularization of the latent space.

In our case, all VAEs with fewer than 8 latent dimensions used the FSQ-VAE approach. It can be applied on top of any output embedding at pre-training or fine-tuning.

#### ZiNB+MSE loss

The goal of the ZINB+MSE loss is to make the model’s expression-level prediction as precise as possible (thanks to the MSE) while preserving the ZiNB’s expressivity and uncertainty estimation. scPRINT-2 uses a novel expression decoder for foundation models that outputs the parameters of a zero-inflated negative binomial (ZiNB) distribution for each gene *i* in cell *j*. The *ZiNB* distribution is defined as

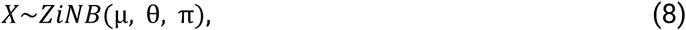

Where the parameters µ, θ, π are obtained from a multi-layer perceptron (MLP) applied to the expression embeddings outputted by the transformer model at its last layer (e), which are the:

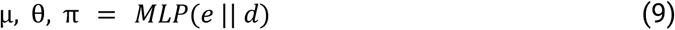

The MLP is a two-layer neural network with dimensions [*d+1, d*, 3], where || denotes the concatenation operation.

Based on the work of Jiang et al.^86^, zero inflation is the best distribution for a broad range of transcriptomic measurements, as some measurements exhibit sufficiently high dropout rates and require a zero-inflation term to model them. In our case, and similarly to scVI^87^, we define our *ZiNB* as

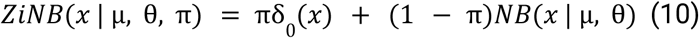

Where δ (*x_j_*) is a point mass at zero, and *NB*(*x_j_*_0_ | µ, θ) is the negative binomial distribution with mean µ and dispersion θ.

Compared to scVI, where the overdispersion parameter θ is learned for each gene, we make scPRINT-2 output it together with µ, π (see Supp. Figure S13)

Effectively, the model learns that dispersion may vary across genes, sequencers, cell types, and sequencing depths.

In addition, the loss adds an MSE term computed from the µ and θ output of the MLP, comparing for a gene *i*, *e* = µ_*i*_ × (1_*i*_ − σ(π_*i*_)) to the logp1-transform of the expression using mean-squared-error.

Where *e*_i_ is the predicted expression of gene *i* and σ is the sigmoid function:

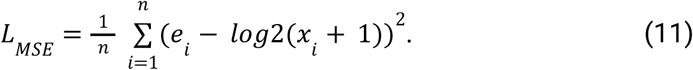

The zinb+mse loss is the addition of both losses with a scale parameter, here:

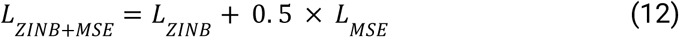

This loss comes as a replacement for the classical MSE or ZiNB in scRNA-seq models.

#### Embedding contrastive loss

The goal of this contrastive loss is to remove some batch effect by pushing cell embeddings obtained from the expression profile after different perturbations to be more similar to each other than they are from cell embeddings of other cell profiles, using the InfoNCE^88^ loss:

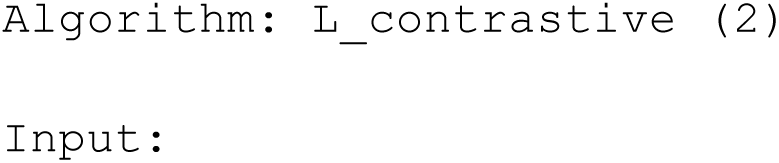
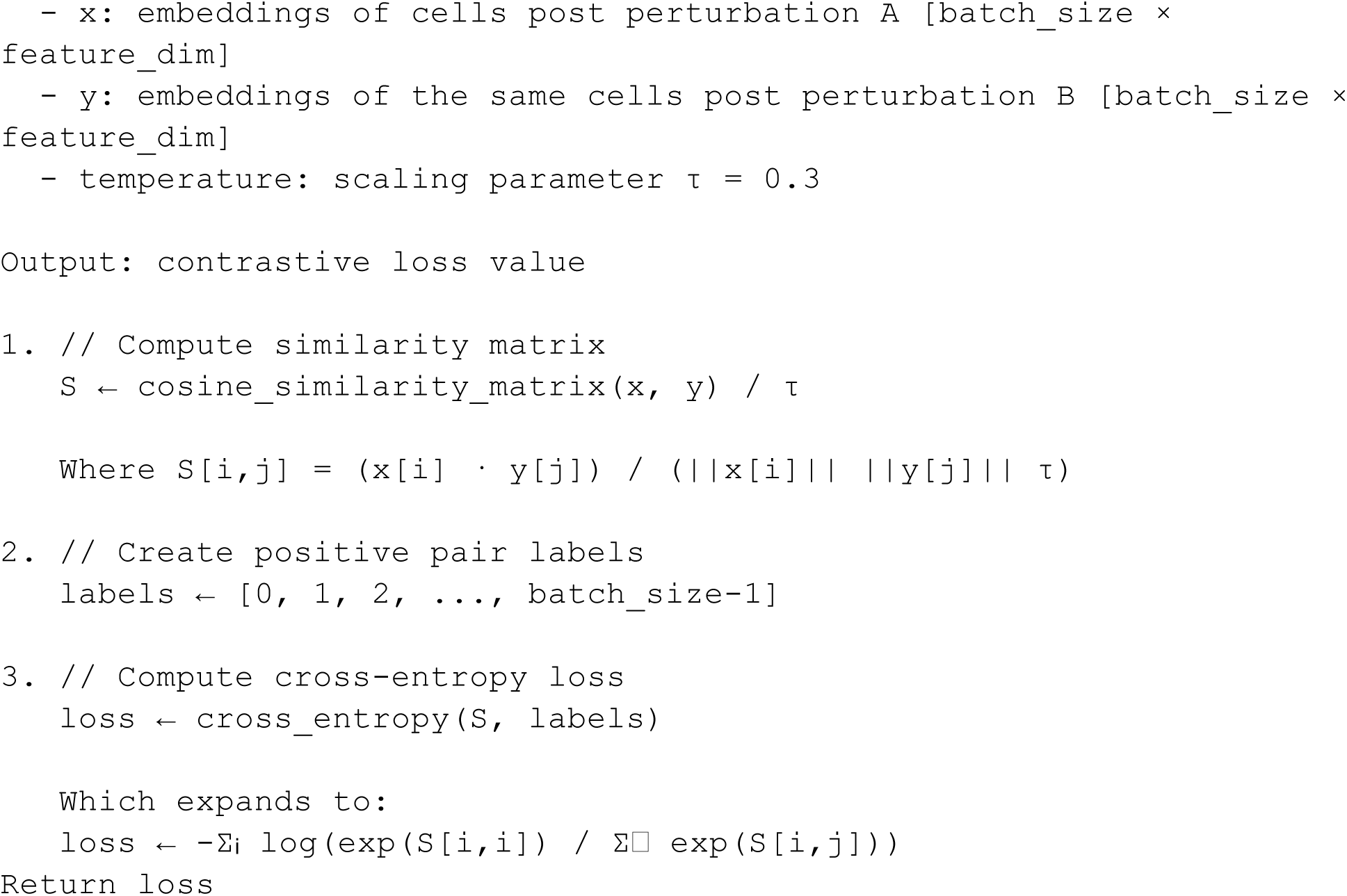

This loss can be added to any scFMs at pre-training or fine-tuning (see supp. Figure S17).

#### Elastic cell similarity loss

The goal of this loss is to reduce batch effects by pushing cells that are similar to become more similar and cells that are dissimilar to become more dissimilar^1^.

We implement the **cell similarity loss** of scGPT, where, given cell embeddings, where *m* is the number of cells and *d_t_* is the embedding dimension:

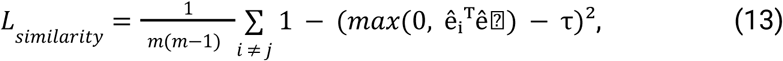

Where:

ê*i* = *e* /||*e* ||₂ is the L2-normalized embedding of the cell *i*

τ is the similarity threshold (default 0.3)

*m*(*m* − 1) is the number of off-diagonal pairs,

This loss can be added to any scFMs.

#### Embedding independence loss

The goal of the embedding independence loss is to push the different class-level embeddings of a cell to encode distinct information by making them orthogonal (see supp. Figure S17).

Implementing a set of disentangled embeddings is not straightforward. In our case, we push the embeddings to be as different from one another as possible, with an **independence loss** defined as

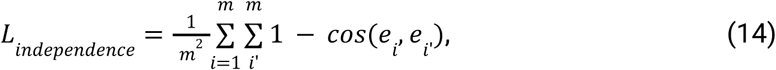

where *e*_i_ and *e*_i_ are the cell embeddings, *m* is the minibatch size, and *cos* denotes the cosine similarity. This pushes each embedding to represent different information from the others.

This loss can be added to any scFMs at pre-training or fine-tuning.

#### Hierarchical classifier loss

The goal of the hierarchical classifier is to enable efficient label predictions for a set of related labels defined by a known graph.

The scPRINT-1 classifier generates predictions for all possible labels in a hierarchical ontology, while producing logits only for the most fine-grained elements. To predict the other elements, it only has to aggregate their children’s logits. We improve this loss in scPRINT-2 by using the entire ontological graph: e.g., if a cell is an *olfactory neuron*, then it is also a neuron. If the classifier predicts *glutaminergic neuron*, it is wrong at this level but correct for *neuron*, meaning we penalize it less overall than a non-neuron label, like *fibroblast* (see Figure 2B).

In conjunction with our weighted sampler, this allows the model to learn rich gradients from a low volume of data. We also implement two additional classes for predictions in our hierarchical classifier compared to scPRINT-1: age and tissue of origin.

During pre-training, we perform label prediction for different classes, e.g., cell type, disease, assay, age, tissue, ethnicity, sex, and organism. We created a specific relabeling of the age label that could be very fine-grained, e.g., 2 weeks, 3 weeks, 35 years old, 36 years old, into biologically relevant groups such as *embryo, fetal*, *6-month-old*, *1-year-old*, *adolescent*, young adult, and so on. We mapped both human and mouse data this way to a common age profile.

These were the only two species with such labels available. The labels follow a hierarchy defined by ontologies: the Cell Ontology for cell type, MONDO for disease, EFO for assay, HANCESTRO for ethnicity, HSAPDV for age, UBERON for tissue, NCBITaxon for organism, and EFO for sex^89–92^. We do not compute the loss for cells with the unknown label.

The algorithm thus becomes:

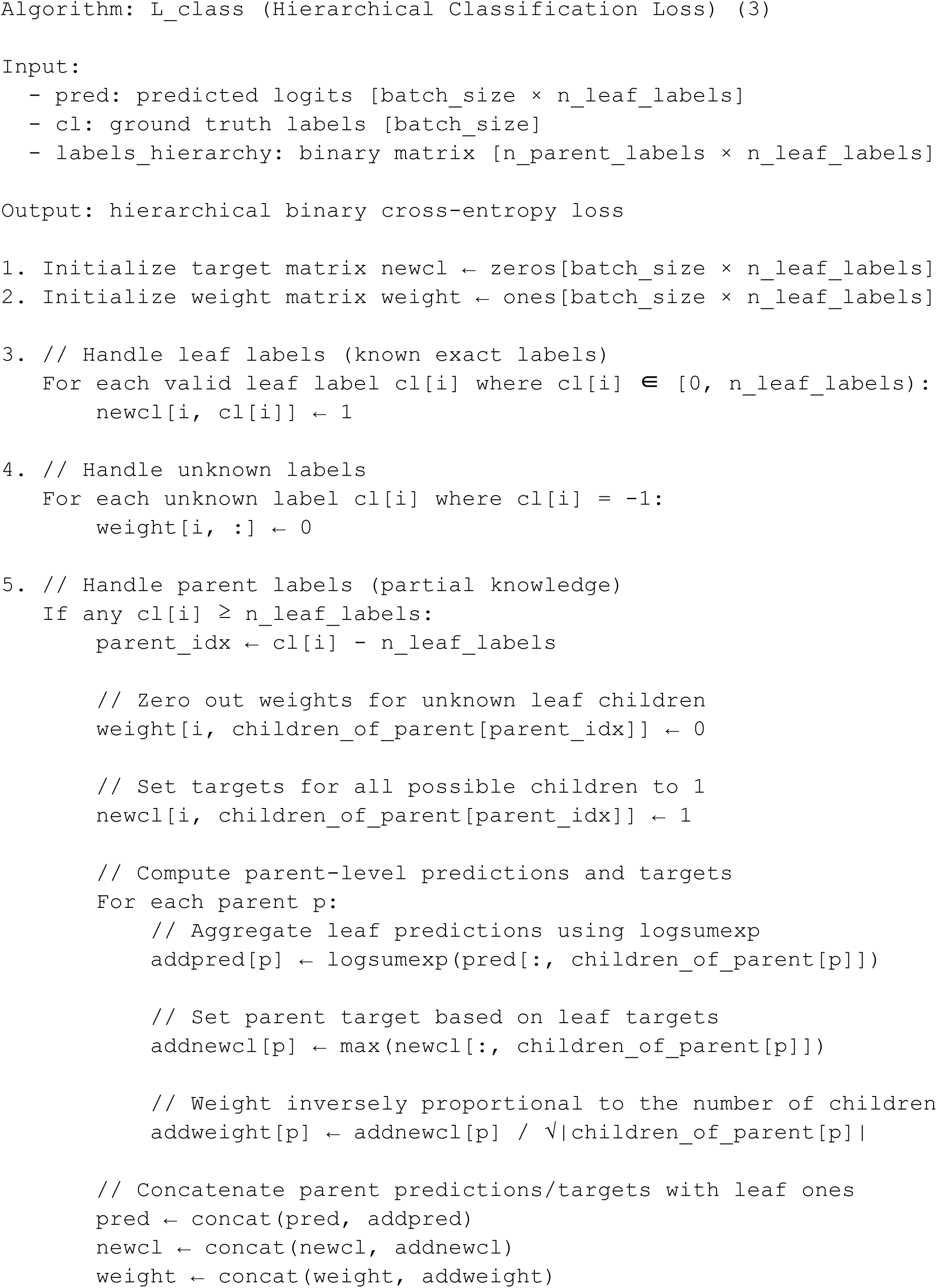
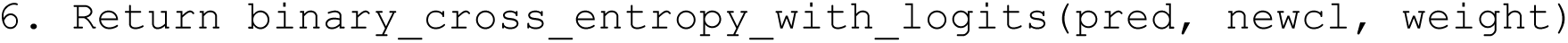

The hierarchical loss is available as a standalone function on GitHub Gist: https://gist.github.com/jkobject/5b36bc4807edb440b86644952a49781e.

This loss replaces a classical pytorch classifier loss, such as

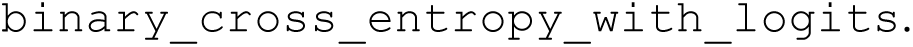

#### Variable context length

The goal of the variable context length method is to decrease the model’s bias toward a specific number of elements in context.

Indeed, we noticed that at inference time, the model’s performance could be lower in variable-context situations (e.g., on gene-panel datasets or when using only expressed genes). We thus introduced a **variable-context** training scheme in which the model’s context sometimes drops by a random amount (see Table 1; see <u>Methods</u>). This makes the model less biased toward a specific input context during inference and decreases training time (see supp. Table S2). Again, here we see strong consistent improvement in the model’s performance across our additive benchmark. This can be applied to any transformer models where the number of elements in context can be chosen arbitrarily.

#### Adversarial classifier loss

The goal of the Adversarial classifier is to remove batch effect^93,94^.

The adversarial classifier is applied only to the *cell_type* cell embedding and is tasked to classify the organism of origin for each cell. It uses the same MLP as regular classifiers (2 layers, 256 as inner dimension). We use the reverse_gradient operation on top of a simple softmax-based binary cross-entropy classifier loss as follows:

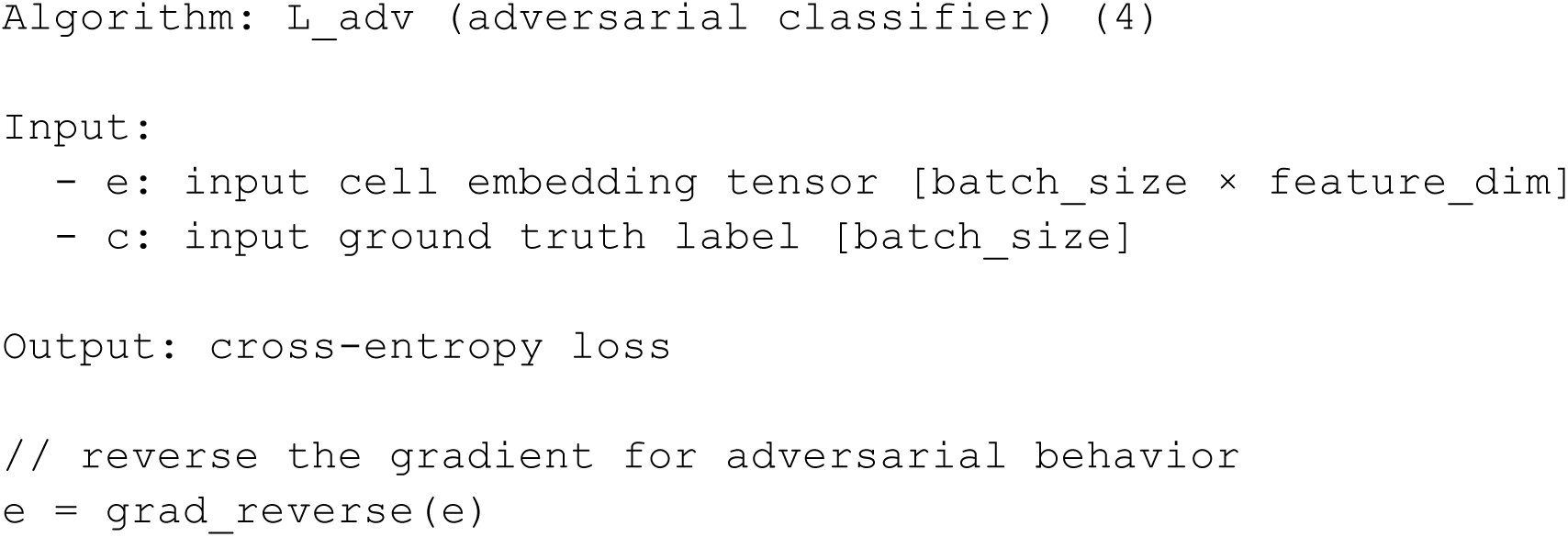
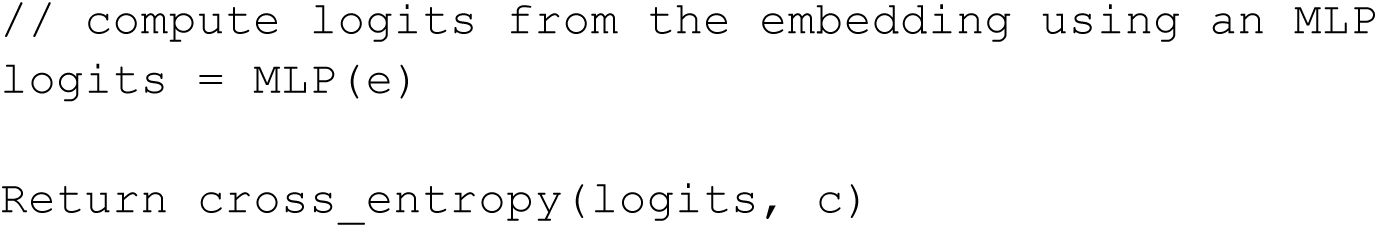

We use it to predict both organisms and sequencers. Sequencers are mapped to a set of coarser labels, as we cannot use the hierarchical classifier in an adversarial context. Indeed, as it is sigmoid-based, it could easily set all label logits to -inf.

This loss can be added during pre-training or finetuning of a foundation model, provided batch labels are available.

#### TF-masking task

The goal of the Transcription Factor (TF) masking task is to push the model to pay more attention to TFs than to other genes.

For the Transcription Factor masking task, we reuse the classic 30% masking task used in the base model (see <u>Base Model</u>). We then list the ENSEMBL IDs of all 13,000 TFs across our 16 organisms and sample our mask, giving increased weight to the TFs. Here, the weight is set up to be 10 for TFs and 1 for the rest.

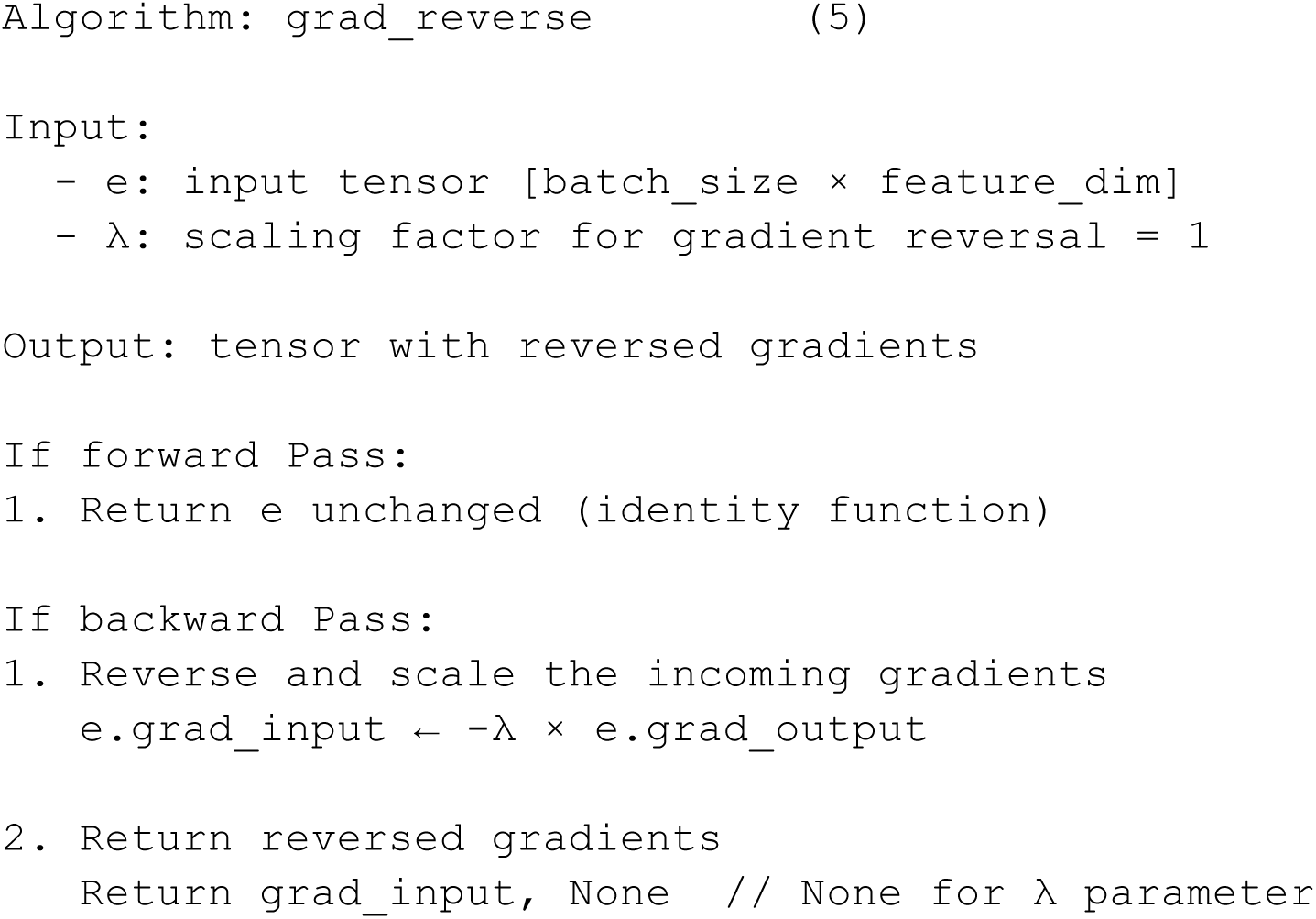

The tool can be applied to any other set of genes as a replacement for classical masking in scFMs.

### Additive Benchmark’s datasets

The gene network analysis is performed on a test kidney single-cell dataset, using 1000 cells from the same cell type, and is compared with the omnipath ground truth (also known as the omnipath benchmark) across all cell types. It is also performed for 1000 K562 cells, comparing it to a network assembled from all genes “i” whose expression changes significantly when gene “j” is perturbed, using a genome-wide perturb-seq dataset called GWPS benchmark^95^.

Knowing that perturb-seq still often implies cell-type- and patient-specific off-target effects and cannot detect many direct effects^96–99^.

The cell type prediction uses accuracy, and batch correction uses scIB v2, as in Kalfon et al.^3^. Both the lung and pancreas datasets have also been used in Kalfon et al. They are test datasets, removed from the pre-training corpus, and both come from the initial scIB paper^100^.

### scPRINT-2

The model architecture is composed of:

– An **encoder/tokenizer** that takes multiple inputs, such as raw expression data, gene names, and gene locations, and embeds them in a high-dimensional space used by the transformer.
– A **trunk** with a bidirectional multi-head transformer, an XPressor bidirectional multi-head transformer, and a set of VAEs applied to each XPressor output embeddings.
– A **class decoder** that transforms the output cell embeddings of the XPressor into cell-specific label prediction logits over a range of classes.
– An **expression decoder** to transform the output embeddings into expression values

Of the above-cited additive benchmark elements, scPRINT-2 contains: **XPressor, all databases, denoising**, **cluster-based sampling, elastic cell similarity, ZINB+MSE, VAE compressor, variable context with larger context, TF masking, GNN expression encoder, and fine-tuned ESM3** (See supp. Figures S1, S9, S17, S18) We now go into some more details about the model:

#### Encoder / Tokenizer

In scPRINT-2, each gene in a cell is converted to an embedding: It corresponds to the sum of 3 different elements:

1. An embedding representing the gene itself using ESM3 with a fine-tuning adaptor layer (see <u>Methods</u>)
2. An embedding of the gene location in the genome. This helps the model understand that genes with similar locations tend to be regulated by similar regulatory regions^101^, a relationship well-known in cellular biology. We encode the genes’ locations using positional encoding. Every gene within 10,000 bp of the next is considered to be in the same location; otherwise, we increment the location by 1. We do this for all genes in the Ensembl database per organism. We then embed these locations using the Positional Encoding (PE) algorithm of Vaswani et al.^83^. We notice that adding this embedding was important to prevent divergence during training.
3. An embedding of the gene expression in the cell and its neighbor using our **GNN** (see <u>Methods</u>)

Finally, during pre-training, a subset of 3200 genes is used to encode a cell expression profile. If fewer than 3200 genes are expressed in both the cell and its neighbors, we pad them with randomly sampled unexpressed genes (meaning with an expression value of 0). This approach allows the model to see different patches of the same cell profile during training.

The full set of embeddings of cell i sent to the transformer is the matrix *X* where

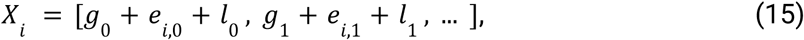

Where *g*_j_ is the gene j encoding, *e*_ij_ is the encoding of the expression of gene j in cell i, *l*_j_ is the gene j location encoding.

Additionally, the Xpressor layers will receive a set of learnt prototype tokens representing the different class-level cell embeddings.

#### Trunk

The model “trunk” is a bidirectional encoder similar to BERT^102^ with *n* layers, *h* attention heads, and a dimension of *d*. It uses the flashattention2^103^ methodology implemented in Triton to compute its attention matrix. It uses the pre-normalization technique^104^, with a sped-up layer norm implemented in Triton’s tutorial^105^. It uses stochastic depth with increasing dropout probability^106^ (see <u>Base for details about small and medium-sized models</u>).

It has a 2-layer MLP with a 4x width increase in its hidden layer and a GELU activation function.

Each Layer or block is composed, in order, of a layer-norm, self-attention, layer-norm, MLP, and layer-norm, cross-attention, layer-norm, MLP, which are only used during the decoding step. It has an additional m Xpressor blocks/layers applied to its 10 latent cell tokens.

The output cell embeddings of the Xpressor layers are then compressed with VAEs with respective latent for the [cell_type, tissue, age, sex, disease, sequencer, ethnicity, organism, cell culture, additional] classes of: 64, 32, 8, 2, 16, 8, 8, 8, 2, None (no VAE)

#### Class Decoders

The class decoders are MLPs applied to compressed representations of their respective VAEs, with a shape of [µ, 256, *d*] with *d* the number of labels in the class c and µ the dimension of this class for the VAE.

#### Expression Decoder

We had noticed that scPRINT-1 initially produced embeddings that could be biased by the cell-depth token. We thus push scPRINT-2 to be depth-invariant by introducing the sequencing depth information only in the Expression Decoder, ensuring that the output gene-cell tokens contain little absolute sequencing depth information (see Figure 3B, see supp Figure S1). This debiases cell embedding to depth data and also improves denoising (see Table 1).

The expression decoder thus gets applied to the output gene embeddings and also receives the log2p1-transformed sequencing depth (also called total cell expression count) *c* andis of the form:

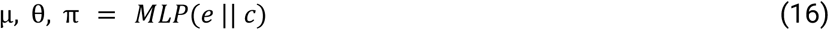

The MLP is a two-layer neural network with dimensions [*d+1, d*, 3], where || denotes the concatenation operation.

The parameters µ, θ, π are the parameters of the ZiNB and are used in the ZINB+MSE loss.

### Pre-training

The three main tasks in the multi-task pre-training of scPRINT-2 are denoising, classification, and bottleneck learning. While the denoising loss enhances the model’s ability to find meaningful gene-gene connections, the other two try to make the model and its cell embedding representation more robust and cell-type-specific. The tasks are presented below.

#### Optimization method

Optimization is performed with fused ADAMW and a weight decay of 0.01. We observed a complete inability to learn when using the base ADAM algorithm, which has a similar weight decay schedule. This can be explained by a known inequivalence issue in ADAM^107^.

We do not use the stochastic weight averaging^108^ method during training.

During pre-training, the hyperparameters are set to a dropout of 0.1, a learning rate (LR) of 1e-4, and the precision is set to 16-mixed with residuals in fp32. We clip gradients to 10 and train over many sub-epochs of 20,000 training and 20,000 validation batches, with a warmup of 2,000 steps. Across epochs, we use a linear LR decrease of 0.6 with a patience of 2, and we stop training after 4 consecutive increases in validation loss. We initialize weights to a normal distribution around 1, biases to 0, and biases for the final layer of the Classifiers to −0.12.

Our batch size is 128, and we use a pre-norm strategy for the transformer with a linearly increasing stochastic depth dropout rate of 0.02 per layer. We use a noise parameter of 70%. We split the cells in the datasets into 98% for training and 2% for validation, and reserve at least 2% of the split datasets for testing. Our reconstruction loss is ZiNB+MSE (see the <u>ZINB+MSE</u> <u>section in Methods</u>).

While many pre-training variants can be selected from contrastive learning, classification, adversarial classification, compression (with XPressor and VAE), masking, biased masking, and imputation, the choice may depend on specific biological assumptions.

scPRINT-2 is trained with denoising an input cell profile, given its nearest neighbor’s expression. Given the same information, it also performs label prediction during pre-training for: cell type, disease, sequencer, age, tissue, ethnicity, sex, cell culture, and organism. The classification task is performed jointly with the denoising task, meaning that labels are predicted from corrupted expression data and from nearest-neighbor expression information. The hierarchical classifier is applied to the VAEs’ latent embeddings.

During decoding, it regenerates the expression profile for all input genes, including those dropped during variable context selection. This effectively does gene imputation.

The decoder receives only the gene location and ESM3 embedding and performs cross-attention on cell embeddings. The cell embeddings are the output of the VAEs and Xpressor layers, so the input is:

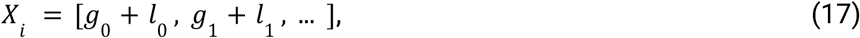

And cell-embeddings are:

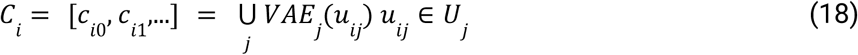

With *U*_*j*_ the matrix output of Xpressor.

Finally, Embedding independence and Elastic Cell similarity losses are applied to the cell embeddings *C*_*i*_ for all cells *i* in the minibatch.

#### Database and sampling

The scPRINT-2 pre-training corpus is composed of all listed databases with weighted random sampling over all predicted labels, together with cluster-weighted sampling to compensate for missing cell-type labels in the Arc’s scBasecount database.

Practically, during training, we apply a curriculum learning strategy whereby the *S*_1_ factor slowly increases from 1 to 1000, letting the model initially learn across the diversity of cells and slowly retrieve the true cell state and modality distribution. We also apply depth-weighted cell sampling to each cell group (see the <u>cluster-weighted sampling section in Methods</u>).

#### Denoising pre-training task

We downsample an expression profile using a zero-inflated Poisson model of the data, following the approach in Kalfon et al. With this formulation, on average, half of the counts to be dropped are removed by randomly selecting some reads per gene, sampled from a Poisson distribution with a lambda parameter proportional to the gene’s count. The remaining half of the counts to be dropped are dropped by randomly setting some genes to 0, i.e., complete dropout of those genes. It is to be noted that, with this definition of downsampling, the exact average number of counts dropped in both parts depends slightly on the dropout *rrate.* During our pre-training, *r* is set to 0.7, meaning, on average, 35% of the transcript counts are dropped per cell.

Let *x_j_* be the gene expression vector of cell i with dimensions *n*_genes_; we create a downsampled *version* by doing

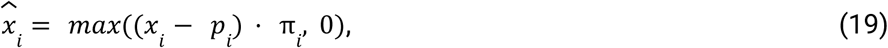

with:

– *m* ∼ *Uniform*(0, *r*) the noise level
– *p*_i_ ∼ *Poisson*(*x_j_* × *r* × 0. 55) a vector of size *d* where the Poisson is sampled for each element *x_j_* of x
– π = *I*(*u* ≥ *r* × 0. 55) a vector of size *n*_genes_, the binary mask vector indicating non-dropout genes.
– *u* ∼ *Uniform*(0, 1), a vector of size *n*_genes_, of random values drawn from a uniform distribution.
– · denotes the element-wise multiplication.
– *r* being the dropout amount. We scale it by a tuning hyperparameter of 0.55 instead of 0.5 for numerical reasons.

We uniformly sample a value between 0 and 0.8 for our *r*, per GPU, during training of scPRINT-2 and other additive models based on denoising, except if noted otherwise.

For the GNN-encoder, we add a second “denoising” step in which we set the noise to 1 and set all expressions to 0 for the center cell. This required the model to predict its expression from the expressions of its neighbors in expression space on the same dataset.

#### Bottleneck learning pre-training task

During training, we predict gene expression at both the decoder output and the scPRINT-2→Expressor→scPRINT-2 pipeline outputs, following the XPressor approach in Kalfon et al. During training, 20% of the time, scPRINT-2 drops between 0 and 2800 genes from its input context per GPU. This pushes the model to learn across a variety of context lengths, it also makes the contrastive loss more robust. Finally, at the output of the decoding step in the bottleneck learning part, the model always predicts across the full 3200 genes, effectively performing imputation during pre-training. When cross-GPU training is performed, cell-embedding-level losses are computed across all GPUs.

#### Classification pre-training task

The Classification task follows the new hierarchical classifier presented in Methods and adds two novel classes: patient age and tissue of origin.

#### Loss aggregation

The losses are aggregated as follows:

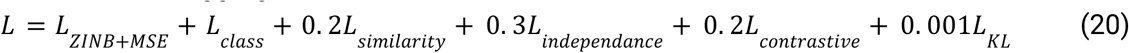

The *L*_ZINE+MSE_ is effectively added 4 times, for the reconstruction post perturbations with denoise of 0.8, 1.0, TF-masking, and post bottleneck learning.

### Fine-tuning Task

Our fine-tuning (see <u>Results</u> section 2 and 4) reuses the classification, bottleneck learning, and VAE (KL) loss of our pre-training for 4 epochs with a learning rate of 0.0001. For batch correction and organism integration, we add the MMD loss between samples from batches 1 and 2 within each minibatch^67,68^. At the same time, an effective MMD loss requires minibatches that are large enough to include a good mix of both label types and cannot accommodate many labels.

With the MMD loss defined as:

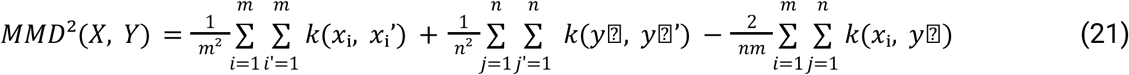

For a finite set of elements from distribution source X and Y, where we use the energy distance kernel:

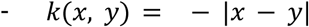

When more than 2 domains exist, we compute MMD between each domain and the remaining domains.

All analyses are defined in the notebook:

notebooks/scPRINT-2-repro-notebooks/fine_tuning_cross_species_emb_mmd.ipynb

### Classification task

For our classifications tasks (see <u>Results</u> section 2), we use the F1-accuracy as our primary metric. When computing it across hierarchical classes, we consider parental relationships to ensure that even if a more precise cell type is predicted than the ground truth, it remains valid.

For example, given a ground truth label of *neuron,* a predicted label of *excitatory neuron* will be considered correct.

If “unknowns” exist in the ground truth or the prediction, they are discarded from the metric.

The cross-organism generalization classification dataset was extracted from the supplementary datasets of the paper titled “Benchmarking cross-organism single-cell RNA-seq data integration methods: towards a cell type tree of life”^46^ available at https://figshare.com/s/6187811b6c3fae02a4d3?file=50608386.

The context-increase classification analysis was performed on the “human multiple cortical areas”^109^ Smart-seq v4 dataset available at https://datasets.cellxgene.cziscience.com/a1d40c84-c81c-406f-bef4-e25edeb651e5.h5ad. For each nnz gene level, we used only cells with at least that many genes expressed. We did not apply the same logic to the second version and used a smaller dataset, so the impact of zero-expressed genes in context could be more clearly seen.

All analyses are defined in the notebooks:

notebooks/scPRINT-2-repro-notebooks/cross_species_embedding.ipynb

notebooks/scPRINT-2-repro-notebooks/smart_seq_class.ipynb

notebooks/scPRINT-2-repro-notebooks/unknown_species_classification.ipynb

notebooks/scPRINT-2-repro-notebooks/large_dataset_anallysis.ipynb figures/nice_umap.py

notebooks/scPRINT-2-repro-notebooks/batch_corr_op ft.ipynb

notebooks/scPRINT-2-repro-notebooks/batch_corr_op v1.ipynb

notebooks/scPRINT-2-repro-notebooks/batch_corr_op.ipynb

notebooks/scPRINT-2-repro-notebooks/plot.ipynb

#### Logits refinement (Laplacian smoothing)

We apply logits smoothing at inference by computing the k-nearest neighbors of each cell and their distances, listed in the squared sparse matrix D, and solving for:

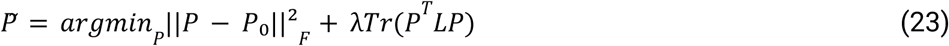

Where *P*_o_ are the initial logits,

L is the graph Laplacian, and λ controls the strength of regularization and has a default value of 0.1^47^. In our case, we set K to be 6, and *L*_ZINE+MSE_ = (*D* + *D*^⊤^ − *C*) where *C* is the diagonal degree matrix of *D* + *D*^⊤^.

The solution has a closed form: *P* = (*I* + λ*L*_ZINE+MSE_)⁻¹*P*_o_

#### Cluster-aggregation

We compute the per-cluster logits aggregation by first clustering the test dataset and then taking the maximum logits across all cells in each cluster as the label for that cluster. Solving for:

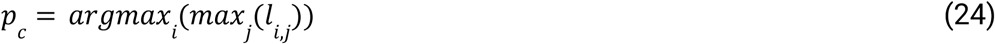

For *l* ∈ *L*_ZINE+MSE_ the j logits across all cells i in the cluster C and *p* the prediction for cluster C

### Denoising task

The denoising benchmark (see Results section 3) was performed on eight datasets of varying quality, assessed by the number of non-zero genes (nnz), sequencing depth, and the distribution of gene counts.

We compute denoising as the dataset-wise percentage improvement in correlation over the 5000 most variable genes, considering only genes that are non-zero in the ground-truth.

Here is the dataset list: retina:

https://datasets.cellxgene.cziscience.com/53bd4177-79c6-40c8-b84d-ff300dcf1b5b.h5ad, kidney:

https://datasets.cellxgene.cziscience.com/01bc7039-961f-4c24-b407-d535a2a7ba2c.h5ad

pancreas: https://figshare.com/ndownloader/files/24539828

intestine:

https://datasets.cellxgene.cziscience.com/d9a99b4a-3755-47c4-8eb5-09821ffbde17.h5ad

glio_smart_highdepth:

https://datasets.cellxgene.cziscience.com/6ec440b4-542a-4022-ac01-56f812e25593.h5ad

lung_smart:

https://datasets.cellxgene.cziscience.com/6ebba0e0-a159-406f-8095-451115673a2c.h5ad human from scbasecount ID: SRX24486462 and SRX22526970

All analyses are defined in the notebook:

notebooks/scPRINT-2-repro-notebooks/denoising_V2.ipynb

### Xenium analysis

We apply the Xenium analysis on the FFPE Human Skin Primary Dermal Melanoma with 5K Human Pan Tissue and Pathways Panel found on the 10X genomics platform under:

https://www.10xgenomics.com/datasets/xenium-prime-ffpe-human-skin

Information on the dataset and its preprocessing can be found on the same webpage.

We extract a dense patch that covers 30% of the cells in the dataset, on which we perform all our analyses (see Results section 3).

All analyses are defined in the notebooks:

notebooks/scPRINT-2-repro-notebooks/xenium_analysis.ipynb

### Embedding task

We perform the organism-level integration task on the same two datasets listed above from the “Benchmarking cross-organism single-cell RNA-seq data integration methods: towards a cell type tree of life” paper, using the scIB metrics and the same ground truth labels (see Results section 4).

All analyses are defined in the notebooks:

notebooks/scPRINT-2-repro-notebooks/cross_species_embedding.ipynb

notebooks/scPRINT-2-repro-notebooks/generative_modelling.ipynb

notebooks/scPRINT-2-repro-notebooks/batch_corr_op ft.ipynb

notebooks/scPRINT-2-repro-notebooks/batch_corr_op v1.ipynb

notebooks/scPRINT-2-repro-notebooks/batch_corr_op.ipynb

notebooks/scPRINT-2-repro-notebooks/plot.ipynb

### Generative task

We perform the generative tasks on two human/mouse datasets extracted from the supplementary datasets of the paper titled “Benchmarking cross-organism single-cell RNA-seq data integration methods: towards a cell type tree of life” (see Results section 4).

We generate cell-embeddings for all mouse cells, giving us a matrix *M* of size [10, *n*_genes_, *d_emb_*].

We then retrieve an average human organism embedding by using 2000 randomly selected human cells and averaging their organism cell-embedding, resulting in a vector v of size [*d_emb_*].

We then regenerate an expression profile using the mouse cell-embeddings and the human average organism embedding by replacing it in the matrix like so: *M*[:, *i*,:] = *v*.

We then apply the decoder part of scPRINT, which performs cross-attention over the matrix M and takes the human gene embeddings as input tokens.

All analyses are defined in the notebook:

#### scRNA-seq datasets distances

To compute our distance metric across two scRNA-seq datasets, we first identify the 5000 most variable genes that are also orthologous between the datasets. We use human and mouse data because orthology was readily accessible and well-defined.

We then compute the W2-distance directly on the raw mouse counts, the humanized counts predicted by scPRINT-2, and the human counts. We do not expect a zero or near-zero W2 distance between the humanized mouse data and the human data, as the number of cells, the types of cell, and their composition differ between the two datasets. We perform a similar analysis of the male-to-female conversion.

#### Over-representation measure

For the overrepresentation analysis and plot, we work on the ordered differential expression gene lists for both human-to-mouse and humanized-mouse-to-mouse, and similarly for male-to-female conversion. We compare the overlap in genes between the two lists at all possible cutoff values from 1 to 5000 to obtain our curve and, therefore, define scores.

### Assessment of gene output embeddings

We assess scPRINT-2’s gene output embeddings by computing output gene embedding oe a random *vascular lymphangioblast* cell from the glioblastoma Smart-seq-v2 dataset using its 5000 most expressed genes in that cell type. We then cluster it using the Leiden algorithm and, for each clustered group of genes, compute the number of pathways enriched using the “KEGG_2021_Human”, “GO_Molecular_Function_2025”, “WikiPathways_2024_Human”, and “GO_Cellular_Component_2025” gene set databases. Doing this for both XPressor and non-XPressor architectures, we then compute a t-test between the two sets of numbers.

All analyses are defined in the notebooks:

notebooks/scPRINT-2-repro-notebooks/output_embeddings.ipynb

### Extracting meta-cell gene networks from attention matrices in scPRINT

Transformers compute multiple attention matrices per layer, called attention heads. This is done by splitting the generated *K, Q*, and *V* embedding into *m* sub-embeddings, thus defining *m* attention heads. Each attention head computes the attention matrix via the equation:

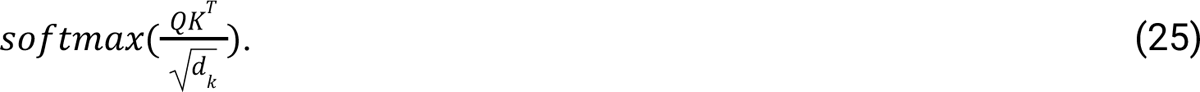

However, we want to aggregate those across multiple cells with similar cell states to increase the signal from a single cell. We are doing so by averaging the Keys and Queries embeddings over the set of cells *U* passed to the model:

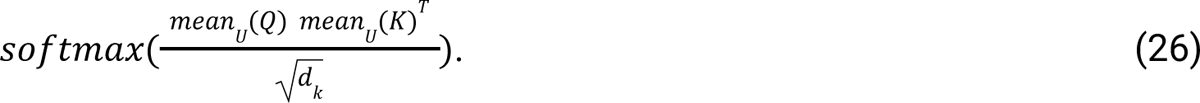

By doing this, the attention matrix behaves as if each query vector for cell i were “looking” across the key vectors of all the cells in U. The resulting object is a row-wise normalized *n*n* matrix, where *n* is the size of the input context (i.e., the number of genes passed to the model).

#### in scPRINT-2

In scPRINT-2, we found, after in-depth review, that while the solution from equation (25) allows for faster computation of much larger gene networks from attention matrices, it also decreases accuracy. We thus instead directly took:

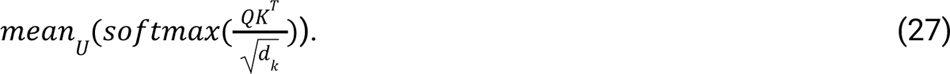

However, to prevent adding QK from genes that are not expressed in the given cell, we generate Qs and Ks from forward passes using only the expressed genes in each cell (see “using only expressed genes” in <u>additive benchmark</u>). This has the benefit of biasing the gene network towards genes that are co-expressed in the set of cells we are computing it on.

This means that for a list of n genes, each cell will have a subset of m Qs,Ks. We thus take the average of the set, computing the mean per gene by counting how many times each gene was expressed across the set of cells.

#### plotting gene sub-networks

To plot a subset of our gene networks, we choose a seed gene and get all its top-K connected nodes. We then overlay the top-N edges in this sub-network, ordered by connection strength. Here K=15 and N=50

### Gene network task

We generated gene networks from notebooks: https://figshare.com/s/6187811b6c3fae02a4d3?file=50608386

We used a matched cross-tissue human and mouse dataset from Zhong et al.^46^

We computed the network across all 10 cell types that were common to both human and mouse in the dataset, using the 4000 most variable genes within each cell type, with a maximum of 1024 cells (see Results section 5).

All analyses are defined in the notebooks: notebooks/scPRINT-2-repro-notebooks/gene_networks.ipynb

notebooks/scPRINT-2-repro-notebooks/gene_networks_var_2.ipynb

#### The Cellmap Ground truth

We used the Cellmap dataset available at https://ndexbio.org under uuid f693137a-d2d7-11ef-8e41-005056ae3c32.

It has a total of 36842 connections across 7543 genes, mainly computed from protein-binding data of AP-MS experiments in the O2US cell line^74^.

#### The Collectri and Omnipath Ground truth

We used the Collectri ground truth from the Decoupler: https://github.com/scverse/decoupler package and the Omnipath ground truth from the Omnipath package^110,111^: https://github.com/saezlab/omnipath, both accessible with given versions within the BenGRN package: https://github.com/jkobject/benGRN.

#### The human interactome Ground truth

We use the RF2-PPI predicted network available at https://conglab.swmed.edu/humanPPI/. We set a cutoff of 0.4 for the benchmark and 0.7 for the high-quality (hq) network^112^.

### Gene network metrics

We use the packages benGRN and GRnnData released with this manuscript to work With Gene networks and perform our benchmarks (see Results section 5).

Our two main metrics are OR and AUPRC. They all take advantage of the fact that the predictions are generated as scores over edges between nodes:

– We have computed the diagnostic odds ratio (OR) as (TP x TN) / (FP x FN) at the cutoff score that yields *K* positive predictions, where *K* is the number of positive elements in the ground truth. In this context, 1 represents a random prediction, and inf represents a perfect prediction; values below one indicate that inverting the predictor would yield better results.
– Area Under the Precision-Recall Curve (AUPRC) is the area (computed with the composite trapezoidal rule) under the curve defined by the precision (*PR = TP / (TP + FP*)) and recall (*RE = TP / (TP + FN*)), where *TP* is the number of true positives. FP is the number of false positives. *FN* is the number of false negatives. This curve is obtained by varying the cutoff from 0 predicted positives to all predicted positives. Here, we compute a version of the AUPRC where the floor of the area is not given by the Precision=0 line but by the prevalence line of the positive class. Moreover, we do not interpolate the curve between the last recall value and the perfect recall: 1. We do this to properly compare AUPRC values across benchmarks and models. Random precision values are given in the supplementary data.

### Open Problem benchmarks

We ran all the open-problem benchmark datasets for scPRINT-2 and scPRINT-1 on a local machine, following the instructions at https://openproblems.bio/documentation. We used the same datasets and labels available at: s3://openproblems-data/resources/ (see <u>Results</u> sections 2 and 4). We used the non-transformed count matrices as input. We used the same metrics for classification, the same scIB package version, and the same train-test splits as in the latest run of Open Problems. All other scores displayed are directly copied from that latest run.

On Open-problems, scIB’s batch correction score is equal to (avgBatch + 1.5*avgBio)/2.5, which are themselves averages over many scores. Details of each value are available in our package’s notebooks^100^.

– scIB avgBio is a combination of label-based and label-free metrics, using, for example, the Adjusted Rand Index (ARI)^113^ and the Normalized Mutual Information (NMI)^114^ on clusters computed from the K-Nearest Neighbor graph. Other scores are used, some based on the conservation of trajectories and cell-cycle variance, others on the conservation of rare-cell populations, the overlap of highly variable genes (see scIB^100^), and more.
– scIB avgBatch is a similar combination of label-based and label-free metrics, using, for example, the average connectivity across clusters of different batches: ASW^115^, the graph integration local inverse Simpson’s Index: graph iLISI^116^, the k-nearest-neighbor Batch Effect Test (kBET)^115^, and more.

Finally, we also use two metrics in our classification task:

– Macro-F1: also called macro-average, is the average of the F1 score across each class in a multi-class task, where the F1 score is: 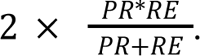.
– Accuracy: is computed as 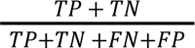

We did not run on two datasets of Open Problems: immune_cell_atlas & tabula_sapiens, as their sizes were too large for us to run scib on any of our available machines.

Moreover, while we believe it is the same for other foundation models assessed in this benchmark, most of these datasets are part of the pre-training corpus of scPRINT. Therefore, the “zero-shot” performance claims, especially classification, should be viewed in this context.

Finally, Open Problem is a living benchmark. Methods, Results, datasets, and metrics will likely change as the scores are continuously updated. We hereby present our results as they were in the 12th of November 2025.

## Data availability

The model weights are publicly available on HuggingFace under: https://huggingface.co/jkobject. pre-training logs to assess the model’s training are publicly available in weights and biases under: https://wandb.ai/ml4ig/scprint_ablation/reports/scPRINT-2-additive-benchmark--VmlldzoxNTIyOTYwNA?accessToken=0mzwwu64py309mds6zzbgcxllrgcdnd10laivhs3ykh9pqmbs0wxutcu60py2bld

The embeddings and classification results over the 350 million cells are available under the public google bucket: <u>gs://scprint2/</u>. The interactive viewer for a subset of these cells is available at https://cantinilab.github.io/scPRINT-2/.

The pre-training dataset is publicly available on CellxGene: https://cellxgene.cziscience.com/, under its census data release version: LTS 2024-07-01, Tahoe and ARC’s scBasecount are available on https://github.com/ArcInstitute/arc-virtual-cell-atlas, commit version 68da110. All other datasets used in this work can be downloaded from their respective public databases using the helper scripts in the scPRINT, BenGRN, GRnnData, and scDataLoader packages.

Source data is provided with this paper to re-generate the figures. Code to download the input dataset, generate the source data, and the figures are available as a notebook in https://github.com/cantinilab/scPRINT-2. Source data are provided with this paper.

## Code availability

The code and notebooks used to develop the model, perform the analyses, and generate results in this study are publicly available and have been deposited in cantinilab/scPRINT-2 at https://github.com/cantinilab/scPRINT-2 under GPLv3 license. The specific version of the code associated with this publication is archived in the same repository under the tag 1.0.0 and is accessible via https://github.com/cantinilab/scPRINT-2/tree/1.0.0/ and DOI:10.5281/zenodo.

Additional packages for this analysis are defined in the pyproject file and project submodules. Together with packages developed by us:

- GrnnData: https://github.com/cantinilab/GRnnData DOI:10.5281/zenodo.10573141
- BenGRN: https://github.com/jkobject/benGRN DOI:10.5281/zenodo.10573209
- scDataLoader: https://github.com/jkobject/scDataLoader DOI:10.5281/zenodo.10573143

## Acknowledgment

The project leading to this manuscript has received funding from the Inception program (Investissement d’Avenir grant ANR-16-CONV-0005) L.C., and the European Union (ERC StG, MULTIview-CELL, 101115618) L.C.. We acknowledge the help of the HPC Core Facility of the Institut Pasteur and Déborah Philipps for the administrative support. L.C..

The work of G. Peyré was supported by the French government under the management of Agence Nationale de la Recherche as part of the ‘Investissements d’avenir’ program, reference ANR19-P3IA-0001 (PRAIRIE 3IA Institute) G.P..

This work was granted access to the HPC resources of IDRIS under the allocation 20 made by GENCI-20XX-AD011014789R2 and GENCI-20XX-AD011016159R1 made by GENCI.

Figure 1B, 1C, 3A, 4D, 4E, 4F, 5G and supplementary Figure S9, S13 used icons by Servier https://smart.servier.com/ is licensed under CC-BY 3.0 Unported https://creativecommons.org/licenses/by/3.0/ NIAID Visual & Medical Arts. RNA. NIAID BIOART Source. bioart.niaid.nih.gov/bioart/452. DBCLS https://togotv.dbcls.jp/en/pics.html is licensed under CC-BY 4.0 International https://creativecommons.org/licenses/by/4.0/. Marcel Tisch https://twitter.com/MarcelTisch is licensed under CC-0 1.0 Universal https://creativecommons.org/publicdomain/zero/1.0/. *Library v1.1. Available via Zenodo* (https://zenodo.org/records/17229908).

## Author Contribution

J.K., L.C., and G.P. designed the study. J.K. developed the tool and performed all the analysis. J.K., and L.C wrote the manuscript. G.P. revised the manuscript.

