## Supplementary tables and figures for "scPRINT-2: Towards the next-generation of cell foundation models and benchmarks"

#### Table S1: detailed version of the additive benchmark

| | names | GPU time per epoch | denoising | embed & batch corr. | | cell type prediction | | gene regulatory network prediction | | | | | |
|---|---|---|---|---|---|---|---|---|---|---|---|---|---|
| | | | score | lung | pancreas | lung | pancreas | OR gwps | OR omni. | AUPRC gwps | AUPRC omni. | | run id |
| Base | ross seeds (masking; ZINB loss; ? + continous expr. emb.; classif. + generative task) | 130 | 2,5 +- 1,5 | 50.1 +-2.1 | 42.3 +-2 | 50 +- 8 | 45 +- 5 | 3,8. +-0.7 | 1,4 +- 0.3 | 0,044 +-0.002 | 0,00165 +-0.00015 | efwkxesx, te0uwaz1, jsls4j6n, hobjefdj, r18nhnuz | blooming-dew-714, driven-valley-750, summer-deluge-783, silent-poltergeist-843, unraveling-pumpkin-841 |
| | **medium model** | 180 | 3 | 50.3 | 42.8 | 65.1 | 60.2 | 4.7 | 1.15 | 0.048 | 0.00188 | p0znio7y | solar-durian-637 |
| | negative control | 0 | -24 | 40.2 | 34.2 | 0 | 0 | 1 | 0.8 | 0.021 | 0.00123 | | |
| architecture | no dropout | 130 | -1 | 49.9 | 45.7 | 47 | 55.1 | 3.9 | 1.5 | 0.044 | 0.00157 | dipgk9u5 | expert-feather-748 |
| | large classifier | 130 | 0 | 52 | 43 | 53 | 49 | 4.4 | 1.8 | 0.046 | 0.00148 | 9q261ics | chocolate-snowball-718 |
| | MVC | 130 | 1 | 51.7 | 44.4 | 54.8 | 45.8 | 3.9 | 1.4 | 0.043 | 0.00143 | yfvvk4cb | celestial-sun-749 |
| | no decoders / generation | 90 | 2 | 52.2 | 44.9 | 53.2 | 47.7 | 4 | 3.5 | 0.044 | 0.00166 | z3abxa21 | northern-voice-777 |
| | **XPressor** | 160 | -4 | 50.4 | 45.6 | 47.3 | 46.8 | 4.2 | 2 | 0.046 | 0.00163 | dsemm200 | crimson-wildflower-791 |
| data | only Tahoe | 130 | 0 | 40 | 33 | 0 | 0 | 4.8 | 0.5 | 0.0041 | 0.00104 | mxu0p3fs | young-bush-669 |
| | CZI + Tahoe (denoising) | 130 | 16.1 | 53.9 | 43.1 | 52.6 | 51.1 | 4.5 | 1.9 | 0.046 | 0.00143 | nmc2l1gf | lurking-cat-846 |
| | CZI | 130 | 1 | 52.3 | 43.1 | 47.8 | 49.1 | 3.8 | 2 | 0.043 | 0.00162 | 4u5c4plu | radiant-oath-793 |
| | **all databases (denoise)** | 140 | -4 | 48.6 | 39 | 44.9 | 40.7 | 3.6 | 1.3 | 0.043 | 0.00157 | ujzjjsi3 | macabre-apparition-844 |
| | 200 human datasets only | 130 | 0 | 49.3 | 43.4 | 40.3 | 50 | 3.5 | 1.8 | 0.044 | 0.00166 | c60vguww | playful-frost-804 |
| | sampling without replacement | 130 | 0 | 50 | 45 | 36 | 34 | 4.5 | 2.4 | 0.045 | 0.0016 | lg84geoq | autumn-aardvark-702 |
| | **cluster-based sampling only** | 130 | 2.7 | 43.3 | 42 | 49.4 | 40.2 | 3.5 | 1.3 | 0.042 | 0.00157 | s8wwlmxr | rosy-firefly-805 |
| | **meta-cell** | 130 | 21 | 52.8 | 47.7 | 53.6 | 51.3 | 3.4 | 1.7 | 0.04 | 0.00155 | gp90j8vn | unique-dawn-806 |
| attention | softpick (larger context) | 130 | 3.1 | 50 | 41 | 53.8 | 44.6 | 3.6 | 1.8 | 0.042 | 0.00156 | s6alkcvp | eldritch-fang-834 |
| | criss-cross (larger context) | 90 | 5.6 | 51.2 | 42.5 | 42.4 | 43.7 | x | x | x | x | u5udxv4v | uncanny-raven-835 |
| | hyper (denoise, larger context) | 160 | 2 | 50.1 | 43.4 | 42.1 | 40.6 | 3.7 | 0.6 | 0.04 | 0.00115 | l44og0s3 | silver-grass-803 |
| loss | contrastive learning (masking + denoising) | 130 | 21.4 | 49 | 41.5 | 39.5 | 40.4 | 4 | 1.3 | 0.043 | 0.00149 | wcg8g3hr | northern-frog-797 |
| | **elastic cell similarity** | 130 | 2.5 | 52.7 | 43.1 | 44.8 | 34.9 | 4.3 | 1.6 | 0.046 | 0.00167 | qn2tyayf | hopeful-monkey-796 |
| | **no embedding ind loss** | 130 | 2.2 | 51.6 | 43 | 50 | 50 | 4 | 2.3 | 0.043 | 0.00156 | 4v84b9nm | devoted-wave-795 |
| | **ZINB+MSE (denoising)** | 130 | 25.5 | 51.3 | 48 | 49.2 | 42.9 | 3.4 | 1.3 | 0.04 | 0.00163 | bv1r4d3h | firm-silence-747 |
| | MSE | 130 | -4 | 54 | 46 | 62 | 43 | 3.3 | 1.2 | 0.042 | 0.00166 | mnk73zbd | generous-dawn-666 |
| | **VAE compressor** | 160 | 3 | 51 | 42 | 38 | 27 | 4.2 | 1.7 | 0.044 | 0.0016 | jwkcrxb9 | wild-terrain-694 |
| pretraining task | **var. context (larger context)** | 170 | 29.1 | 53 | 46 | 52.9 | 52.2 | 3.1 | 1.2 | 0.038 | 0.00146 | 44p3fv3v | apricot-snowflake-756 |
| | **TF masking** | 130 | 2.6 | 49.8 | 42.8 | 49.8 | 42.8 | 3.7 | 2.3 | 0.043 | 0.00169 | 8vmjnnsb | winter-meadow-772 |
| | **denoising** | 130 | 21 | 52.6 | 45.1 | 50.9 | 54.5 | 3.6 | 1.3 | 0.043 | 0.0016 | bk37305v | efficient-firebrand-753 |
| | no classification | 130 | 3 | 50 | 40 | 0 | 0 | 3.9 | 1.2 | 0.043 | 0.00129 | oxlxtzim | copper-frost-625 |
| | adv. classifier (+larger classif) | 130 | 1 | 52 | 42 | 48 | 43 | 4.1 | 1.6 | 0.044 | 0.0014 | 2tzkv7m8 | sunny-morning-629 |
| input | sum normalization (denoise) | 130 | 12.8 | 45.6 | 46.5 | 21.4 | 22.9 | 2.4 | 1 | 0.029 | 0.00136 | ldh1fw8d | snowy-galaxy-744 |
| | no random level of denoising | 130 | 19 | 54.1 | 45.3 | 50.7 | 45.2 | 3.6 | 2 | 0.041 | 0.00179 | 0ayw97iw | divine-monkey-798 |
| | binning | 130 | 0 | 51.8 | 45.5 | 58.4 | 52 | 4.2 | 1.3 | 0.047 | 0.00162 | op7at8xm | twilight-breeze-874 |
| | **GNN expression encoder** | 150 | 44 | 48 | 42 | 38 | 35 | 4 | 1.4 | 0.042 | 0.00128 | bv6d9wpl | balmy-totem-727 |
| | using only expressed genes | 130 | 1.2 | 52.2 | 42.9 | 53.2 | 40.1 | 3.8 | 1.3 | 0.043 | 0.00157 | xz238fyr | sage-snow-873 |
| | without gene location | 130 | 3.4 | 36.2 | 35 | 4 | 5.9 | 4.8 | 1.5 | 0.048 | 0.0017 | r2n83z4k | youthful-snowflake-792 |
| | learn gene emb (denoising) | 130 | 20.9 | 51.7 | 45.1 | 49.7 | 46 | 3.2 | 1.7 | 0.041 | 0.00154 | npayct6q | jumping-night-755 |
| | **fine-tuned ESM3** | 130 | 21.4 | 51.5 | 42.8 | 55.6 | 44.2 | 3.7 | 1.4 | 0.042 | 0.00181 | fkcgp56s | not-snowflake-755 |
| Main | small model (V2) | 1820 | 44 | 53 | 49 | 46 | 47 | 3.5 | 1.6 | 0.041 | 0.0015 | | honest-vortex-815 |
| | medium model (V2) | 5600 | x | x | x | x | x | x | x | x | x | | x |
| | medium model (V1) | 520 | 20.9 | 52.6 | 45.6 | 61.8 | 57.6 | 3.4 | 2.2 | 0.041 | 0.0017 | | bewitched-poltergeist-857 |
| | small model (V1) | 160 | 31.7 | 52.4 | 50 | 44.7 | 44.7 | 3.6 | 1.5 | 0.042 | 0.00138 | | dry-smoke-852 |

Detailed version of the additive benchmark, listing every value.

### Table S2: Full detailed table of the additive benchmark scores

| Name | Notes | grn_omni_old_kidney/or | grn_gwps/auprc | # grn_gwps/epr | grn_omni_lung_smart/or | emb_bone_marrow_5batch/ct_class | emb_lung/ct_class | emb_pancreas/ct_class | emb_bone_marrow_5batch/scib | emb_lung/scib | emb_pancreas/scib | emb_kidney/ct_class | emb_kidney/scib | emb_pancreas/scib_batch | emb_pancreas/scib_bio | denoise_retina/reco2full_vs_noisy2full | denoise_lung_smart/reco2full_vs_noisy2full | denoise_kidney/reco2full_vs_noisy2full | denoise_glio_smart_highdepth/reco2full_vs_noisy2full | denoise_intestine/reco2full_vs_noisy2full |
|---|---|---|---|---|---|---|---|---|---|---|---|---|---|---|---|---|---|---|---|---|
| Name: twilight-breeze-874 | binned | | | | | | | | | | | | | | | | | | | |
| Name: macabre-apparition-844 | scPRINT-V2 (all+tahoe+scbase) filtered | 1.27 | 0.04 | 3.58 | 2.60 | 0.29 | 0.45 | 0.41 | 0.44 | 0.49 | 0.39 | 0.38 | 0.38 | 0.24 | 0.49 | -0.05 | 0.33 | -0.04 | 0.23 | -0.16 |
| Name: eldritch-fang-834 | softpick-flash | 1.59 | 0.03 | 3.36 | 1.36 | 0.29 | 0.55 | 0.44 | 0.46 | 0.51 | 0.41 | 0.44 | 0.43 | 0.24 | 0.52 | -0.01 | 0.00 | 0.03 | 0.02 | 0.00 |
| Name: silver-grass-803 | hyper attention | 0.66 | 0.04 | 3.73 | 2.53 | 0.31 | 0.42 | 0.41 | 0.47 | 0.50 | 0.43 | 0.42 | 0.46 | 0.29 | 0.53 | -0.01 | 0.01 | 0.02 | -0.08 | -0.02 |
| Name: sage-snow-873 | mask zeros | 1.27 | 0.04 | 3.83 | 1.48 | 0.31 | 0.53 | 0.40 | 0.48 | 0.52 | 0.43 | 0.49 | 0.48 | 0.24 | 0.55 | 0.00 | 0.01 | 0.01 | 0.00 | 0.00 |
| Name: lurking-cat-846 | (all+tahoe) filtered | 1.93 | 0.05 | 4.55 | 2.60 | 0.29 | 0.53 | 0.51 | 0.49 | 0.54 | 0.43 | 0.44 | 0.50 | 0.28 | 0.53 | 0.16 | 0.37 | 0.16 | 0.32 | 0.32 |
| Name: uncanny-raven-835 | criss-cross | 0.73 | 0.03 | 2.00 | 0.61 | 0.26 | 0.42 | 0.44 | 0.47 | 0.51 | 0.43 | 0.48 | 0.48 | 0.26 | 0.54 | 0.08 | 0.32 | 0.06 | 0.22 | -0.09 |
| Name: unraveling-pumpkin-841 | seed 42 regular | 1.31 | 0.04 | 3.76 | 2.27 | 0.31 | 0.57 | 0.40 | 0.47 | 0.51 | 0.42 | 0.43 | 0.43 | 0.23 | 0.54 | -0.01 | 0.00 | 0.01 | 0.00 | -0.01 |
| Name: silent-poltergeist-843 | seed 122 regular | 1.25 | 0.04 | 3.97 | 2.40 | 0.30 | 0.49 | 0.43 | 0.47 | 0.48 | 0.40 | 0.44 | 0.44 | 0.24 | 0.52 | -0.01 | 0.01 | 0.02 | -0.01 | 0.00 |
| Name: summer-deluge-783 | regular | 1.34 | 0.04 | 4.11 | 1.50 | 0.31 | 0.50 | 0.49 | 0.49 | 0.52 | 0.44 | 0.47 | 0.46 | 0.27 | 0.56 | 0.01 | 0.00 | 0.03 | 0.00 | 0.00 |
| Name: driven-valley-750 | regular 2 | 1.49 | 0.04 | 4.18 | 2.14 | 0.25 | 0.47 | 0.45 | 0.49 | 0.50 | 0.42 | 0.37 | 0.45 | 0.24 | 0.53 | 0.00 | 0.01 | 0.03 | 0.01 | 0.00 |
| Name: rural-pine-670 | regular | | 0.05 | 4.67 | | 0.44 | 0.50 | 0.16 | 0.47 | 0.52 | 0.45 | 0.55 | 0.48 | 0.26 | 0.57 | -0.01 | 0.02 | 0.01 | 0.02 | -0.01 |
| Name: devoted-dragon-662 | normal | | 0.05 | 4.09 | | 0.37 | 0.63 | 0.22 | 0.48 | 0.51 | 0.43 | 0.51 | 0.46 | 0.25 | 0.55 | 0.01 | 0.30 | 0.03 | 0.28 | 0.02 |
| Name: northern-frog-797 | cce noise mask | 1.32 | 0.04 | 4.00 | 1.19 | 0.25 | 0.40 | 0.40 | 0.46 | 0.49 | 0.41 | 0.38 | 0.44 | 0.27 | 0.51 | 0.20 | 0.34 | 0.21 | 0.33 | 0.32 |
| Name: rosy-firefly-805 | clust_cell_type | 1.29 | 0.04 | 3.49 | 1.27 | 0.17 | 0.43 | 0.40 | 0.45 | 0.49 | 0.42 | 0.39 | 0.47 | 0.24 | 0.54 | 0.01 | 0.01 | 0.03 | -0.01 | 0.00 |
| Name: youthful-snowflake-792 | no gene pos | 1.54 | 0.05 | 4.80 | 1.83 | 0.01 | 0.04 | 0.06 | 0.36 | 0.36 | 0.35 | 0.04 | 0.31 | 0.17 | 0.47 | -0.01 | 0.08 | 0.03 | 0.17 | 0.00 |
| Name: crimson-wildflower-791 | Xpressor | 2.05 | 0.05 | 4.17 | 1.78 | 0.30 | 0.47 | 0.47 | 0.49 | 0.50 | 0.46 | 0.32 | 0.43 | 0.27 | 0.58 | -0.04 | 0.01 | -0.04 | 0.02 | -0.01 |
| Name: radiant-oath-793 | V2 full | 3.21 | 0.04 | 3.93 | 2.03 | 0.22 | 0.48 | 0.49 | 0.48 | 0.52 | 0.43 | 0.38 | 0.45 | 0.27 | 0.54 | -0.01 | 0.01 | 0.01 | 0.00 | -0.02 |
| Name: devoted-wave-795 | no emb dissim | 2.33 | 0.04 | 4.05 | 1.90 | 0.23 | 0.50 | 0.50 | 0.48 | 0.52 | 0.43 | 0.43 | 0.46 | 0.26 | 0.54 | -0.01 | 0.00 | 0.02 | 0.01 | -0.01 |
| Name: hopeful-monkey-796 | ecs (I like it) | 1.72 | 0.05 | 4.32 | 1.44 | 0.27 | 0.51 | 0.35 | 0.48 | 0.53 | 0.43 | 0.31 | 0.45 | 0.29 | 0.53 | 0.01 | 0.02 | 0.02 | 0.02 | 0.00 |
| Name: divine-monkey-798 | noise without randsamp | 1.97 | 0.04 | 3.64 | 1.54 | 0.34 | 0.51 | 0.45 | 0.50 | 0.54 | 0.45 | 0.42 | 0.47 | 0.28 | 0.57 | 0.17 | 0.39 | 0.19 | 0.34 | 0.29 |
| Name: unique-dawn-806 | metacell | 1.89 | 0.04 | 3.39 | 2.69 | 0.30 | 0.56 | 0.53 | 0.49 | 0.52 | 0.48 | 0.45 | 0.49 | 0.30 | 0.60 | 0.18 | 0.40 | 0.21 | 0.34 | 0.33 |
| Name: playful-frost-804 | some good quality human | 1.80 | 0.04 | 3.47 | 1.19 | 0.23 | 0.41 | 0.50 | 0.46 | 0.49 | 0.43 | 0.49 | 0.46 | 0.28 | 0.54 | -0.01 | 0.01 | 0.00 | 0.00 | -0.01 |
| Name: winter-meadow-772 | TF masking | 2.25 | 0.04 | 3.75 | 1.80 | 0.34 | 0.50 | 0.47 | 0.47 | 0.50 | 0.43 | 0.37 | 0.45 | 0.27 | 0.54 | 0.01 | 0.00 | 0.03 | 0.00 | 0.00 |
| Name: efficient-firebrand-753 | denoise | 1.30 | 0.04 | 3.58 | 1.39 | 0.25 | 0.51 | 0.54 | 0.49 | 0.53 | 0.45 | 0.52 | 0.48 | 0.30 | 0.56 | 0.19 | 0.36 | 0.21 | 0.31 | 0.32 |
| Name: worthy-snowflake-754 | fine tune gene emb | 1.82 | 0.04 | 3.88 | 2.88 | 0.34 | 0.56 | 0.49 | 0.47 | 0.53 | 0.43 | 0.47 | 0.47 | 0.27 | 0.54 | 0.00 | 0.03 | 0.02 | 0.03 | 0.03 |
| Name: northern-voice-777 | no generate | 3.49 | 0.04 | 4.05 | 2.93 | 0.27 | 0.53 | 0.48 | 0.47 | 0.52 | 0.45 | 0.45 | 0.47 | 0.28 | 0.56 | -0.01 | 0.27 | 0.02 | 0.27 | 0.01 |
| Name: expert-feather-748 | no dropout | 1.53 | 0.04 | 3.92 | 3.52 | 0.28 | 0.47 | 0.55 | 0.49 | 0.50 | 0.46 | 0.55 | 0.45 | 0.28 | 0.58 | -0.01 | 0.01 | -0.01 | 0.01 | 0.00 |
| Name: apricot-snowflake-756 | var context len | 1.20 | 0.04 | 3.13 | 1.84 | 0.36 | 0.53 | 0.52 | 0.49 | 0.53 | 0.46 | 0.44 | 0.50 | 0.29 | 0.57 | 0.16 | 0.38 | 0.21 | 0.34 | 0.29 |
| Name: not-snowflake-755 | finetune gene emb + denoise | 1.37 | 0.04 | 3.75 | 1.83 | 0.30 | 0.56 | 0.44 | 0.47 | 0.52 | 0.43 | 0.37 | 0.46 | 0.26 | 0.54 | 0.20 | 0.37 | 0.21 | 0.32 | 0.30 |
| Name: snowy-galaxy-744 | sum norm | 1.05 | 0.03 | 2.39 | 3.40 | 0.13 | 0.21 | 0.23 | 0.41 | 0.46 | 0.46 | 0.16 | 0.48 | 0.40 | 0.51 | 0.11 | 0.28 | 0.13 | 0.25 | 0.28 |
| Name: celestial-sun-749 | mvc | 1.40 | 0.04 | 3.88 | 3.25 | 0.24 | 0.55 | 0.46 | 0.48 | 0.52 | 0.44 | 0.42 | 0.47 | 0.29 | 0.55 | 0.00 | -0.02 | 0.01 | 0.00 | 0.00 |
| Name: firm-silence-747 | zinb_mse | 1.34 | 0.04 | 3.41 | 1.60 | 0.29 | 0.49 | 0.43 | 0.53 | 0.51 | 0.48 | 0.47 | 0.54 | 0.34 | 0.57 | 0.22 | 0.28 | 0.26 | 0.21 | 0.34 |
| Name: wise-frog-736 | large clf and weight scaler increase | 1.76 | 0.05 | 4.52 | 1.63 | 0.23 | 0.52 | 0.42 | 0.47 | 0.52 | 0.43 | 0.37 | 0.46 | 0.26 | 0.53 | 0.01 | 0.02 | 0.02 | 0.02 | 0.00 |
| Name: mild-valley-737 | debugged TF masking | 1.86 | 0.04 | 3.92 | 3.05 | 0.31 | 0.50 | 0.45 | 0.48 | 0.52 | 0.43 | 0.42 | 0.46 | 0.27 | 0.55 | 0.00 | -0.02 | 0.02 | 0.00 | -0.01 |
| Name: balmy-totem-727 | GNN | 1.38 | 0.04 | 4.07 | 1.08 | 0.28 | 0.38 | 0.35 | 0.46 | 0.48 | 0.42 | 0.48 | 0.47 | 0.25 | 0.54 | 0.33 | 0.46 | 0.32 | 0.36 | 0.44 |
| Name: colorful-bush-721 | zinb+noise + mse | 1.93 | 0.04 | 3.42 | 2.00 | 0.34 | 0.53 | 0.46 | 0.51 | 0.55 | 0.45 | 0.39 | 0.54 | 0.31 | 0.55 | 0.22 | 0.30 | 0.23 | 0.18 | 0.31 |
| Name: hopeful-totem-719 | metacell mode denoise | 1.08 | 0.04 | 3.38 | 1.38 | 0.31 | 0.54 | 0.53 | 0.49 | 0.54 | 0.46 | 0.47 | 0.50 | 0.30 | 0.57 | 0.17 | 0.38 | 0.20 | 0.33 | 0.32 |
| Name: polished-morning-726 | adv classifier | 1.49 | 0.05 | 4.16 | 2.19 | 0.24 | 0.43 | 0.43 | 0.47 | 0.51 | 0.46 | 0.33 | 0.43 | 0.28 | 0.58 | 0.01 | 0.02 | 0.04 | 0.02 | 0.00 |
| Name: chocolate-snowball-718 | large clsf | 1.45 | 0.05 | 4.39 | 1.50 | 0.26 | 0.53 | 0.49 | 0.47 | 0.53 | 0.44 | 0.38 | 0.46 | 0.26 | 0.56 | -0.01 | 0.01 | 0.02 | 0.01 | -0.01 |
| Name: blooming-dew-714 | normal | 1.57 | 0.04 | 3.73 | 2.14 | 0.30 | 0.50 | 0.45 | 0.48 | 0.50 | 0.43 | 0.43 | 0.44 | 0.26 | 0.54 | 0.00 | 0.02 | 0.02 | 0.01 | 0.00 |
| Name: sage-bird-657 | binning | 1.33 | 0.05 | 4.24 | 3.68 | 0.23 | 0.43 | 0.41 | 0.48 | 0.52 | 0.46 | 0.29 | 0.48 | 0.31 | 0.57 | -0.02 | -0.01 | -0.02 | 0.00 | -0.02 |
| Name: leafy-sea-695 | larger classif | 1.13 | 0.04 | 3.65 | 2.43 | 0.21 | 0.44 | 0.38 | 0.47 | 0.51 | 0.43 | 0.41 | 0.46 | 0.25 | 0.55 | 0.00 | 0.06 | 0.02 | 0.02 | 0.00 |
| Name: autumn-aardvark-702 | no rws | 2.45 | 0.04 | 3.99 | 1.09 | 0.21 | 0.36 | 0.34 | 0.48 | 0.50 | 0.45 | 0.36 | 0.48 | 0.25 | 0.57 | -0.01 | 0.01 | 0.02 | 0.01 | -0.01 |
| Name: wild-terrain-694 | VAE | 1.69 | 0.04 | 4.02 | 1.50 | 0.19 | 0.38 | 0.27 | 0.53 | 0.51 | 0.42 | 0.25 | 0.46 | 0.31 | 0.50 | 0.01 | 0.14 | 0.03 | 0.16 | 0.00 |
| Name: generous-dawn-666 | MSE | 1.21 | 0.04 | 3.46 | 1.62 | 0.48 | 0.63 | 0.23 | 0.54 | 0.55 | 0.47 | 0.50 | 0.51 | 0.25 | 0.61 | -0.04 | -0.02 | -0.08 | -0.01 | -0.04 |
| Name: copper-frost-625 | no clss | 1.17 | 0.04 | 3.50 | 1.82 | 0.00 | 0.00 | 0.00 | 0.48 | 0.50 | 0.40 | 0.00 | 0.47 | 0.22 | 0.52 | 0.01 | 0.23 | 0.03 | 0.24 | 0.02 |
| Name: faithful-dragon-663 | V2 full results | | 0.04 | 3.75 | | 0.31 | 0.67 | 0.18 | 0.47 | 0.51 | 0.44 | 0.54 | 0.49 | 0.26 | 0.55 | 0.00 | 0.25 | 0.03 | 0.24 | 0.00 |
| Name: solar-durian-637 | regular medium model | | 0.05 | 4.70 | | 0.52 | 0.65 | 0.60 | 0.48 | 0.50 | 0.43 | 0.62 | 0.48 | 0.28 | 0.53 | 0.02 | 0.02 | 0.03 | | 0.03 |
| Name: sunny-morning-629 | adv classifier | | 0.04 | 4.12 | | 0.41 | 0.47 | 0.11 | 0.46 | 0.49 | 0.42 | 0.34 | 0.37 | 0.26 | 0.53 | 0.00 | 0.01 | 0.00 | -0.01 | -0.02 |

details of the additive study as copied from wandb. processed scores can be viewed in supp. table 1

### Table S3: detailed scIB biological conservation scores on the xenium dataset

| Bio Metric | Isolated labels | KMeans NMI | KMeans ARI | Silhouette label | cLISI Bio conservation | **Aggregate score** |
|---|---|---|---|---|---|---|
| X_pca | 0.446 | **0.255** | **0.036** | 0.382 | 0.955 | **0.415** |
| scprint_emb | 0.468 | **0.376** | **0.125** | 0.383 | 0.979 | **0.466** |

scPRINT vs PCA on expression. ScPRINT performs better, likely by denoising the expression.

### Table S4: detailed scIB scores on the unseen species integration task

| | **Bio conservation** | | | | | **Batch correction** | | | | | **Aggregate score** | | | **Recomputed scores using the same elements** | | |
|---|---|---|---|---|---|---|---|---|---|---|---|---|---|---|---|---|
| | Isolated labels | KMeans NMI | KMeans ARI | Silhouette label | cLISI | BRAS | iLISI | KBET | Graph connectivity | PCR comparison | Batch correction | Bio conservation | **Total** | Batch correction | Bio conservation | **Total** |
| **scprint_2 ft (cell_type emb)** | 0.64 | 0.81 | 0.75 | 0.69 | 1.00 | 0.69 | 0.00 | 0.01 | 0.91 | 0.00 | 0.32 | 0.78 | **0.60** | | | |
| **scprint_2 ft** | 0.55 | 0.72 | 0.63 | 0.56 | 1.00 | 0.58 | 0.00 | 0.00 | 0.65 | 0.00 | 0.25 | 0.69 | **0.51** | | | |
| **scprint_zeroshot (cell_type emb)** | 0.57 | 0.41 | 0.31 | 0.53 | 0.98 | 0.65 | 0.00 | 0.77 | 0.00 | 0.28 | 0.56 | 0.45 | **0.49** | | | |
| **scprint_zeroshot** | 0.49 | 0.00 | 0.00 | 0.49 | 0.68 | 1.00 | 0.86 | 0.00 | 0.20 | 1.00 | 0.61 | 0.33 | **0.44** | | | |
| **random** | 0.54 | 0.23 | 0.14 | 0.50 | 0.97 | 0.80 | 0.00 | 0.00 | 0.72 | 0.00 | 0.30 | 0.48 | **0.41** | | | |
| **no integration (pca)** | 0.57 | 0.21 | 0.10 | 0.36 | 0.99 | 0.69 | 0.00 | 0.00 | 0.60 | 0.00 | 0.26 | 0.44 | **0.37** | | | |
| saturn | 0.81 | 0.97 | 0.49 | 0.13 | 1.00 | 0.79 | 0.13 | 0.05 | 0.91 | 0.92 | 0.62 | 0.92 | **0.79** | 0.56 | 0.68 | **0.63** |
| scGen | 0.60 | 0.77 | 0.49 | 0.23 | 0.99 | 0.88 | 0.23 | 0.16 | 0.91 | 0.92 | 0.85 | 1.00 | **0.68** | 0.62 | 0.61 | **0.62** |
| Seurat v4 CCA | 0.58 | 0.57 | 0.48 | 0.23 | 0.97 | 0.84 | 0.23 | 0.13 | 0.90 | 0.89 | 0.73 | 0.92 | **0.50** | 0.60 | 0.57 | **0.58** |
| SAMap | | 0.62 | | 0.01 | 0.98 | 0.91 | 0.01 | 0.22 | 0.74 | | 0.60 | 1.00 | **0.47** | 0.47 | 0.53 | **0.51** |
| scVI | 0.51 | 0.55 | 0.50 | 0.23 | 0.95 | 0.83 | 0.23 | 0.09 | 0.91 | 0.98 | 0.80 | 0.93 | **0.47** | 0.61 | 0.55 | **0.57** |
| BBKNN | | 0.56 | | 0.11 | 0.99 | 0.82 | 0.11 | 0.05 | 0.82 | | 0.31 | 0.66 | **0.41** | 0.45 | 0.55 | **0.51** |
| Scanorama | 0.56 | 0.54 | 0.49 | 0.27 | 0.96 | 0.76 | 0.27 | 0.09 | 0.84 | 0.93 | 0.59 | 0.79 | **0.37** | 0.58 | 0.56 | **0.57** |
| fastMNN | 0.52 | 0.54 | 0.48 | 0.09 | 0.96 | 0.70 | 0.09 | 0.03 | 0.89 | 0.86 | 0.37 | 0.72 | **0.36** | 0.51 | 0.52 | **0.52** |
| Harmony | 0.51 | 0.54 | 0.44 | 0.06 | 0.96 | 0.70 | 0.06 | 0.03 | 0.86 | 0.70 | 0.15 | 0.70 | **0.16** | 0.47 | 0.50 | **0.49** |

Details of the full scIB results comparing no integration, random embeddings sampled from the multivariate Gaussian, and different versions of scPRINT zero-shot or fine-tuned, using the merged embeddings or the cell type ones

### Supplementary figures

#### FIG S1: illustration of the full scPRINT-2's architecture, input, and output

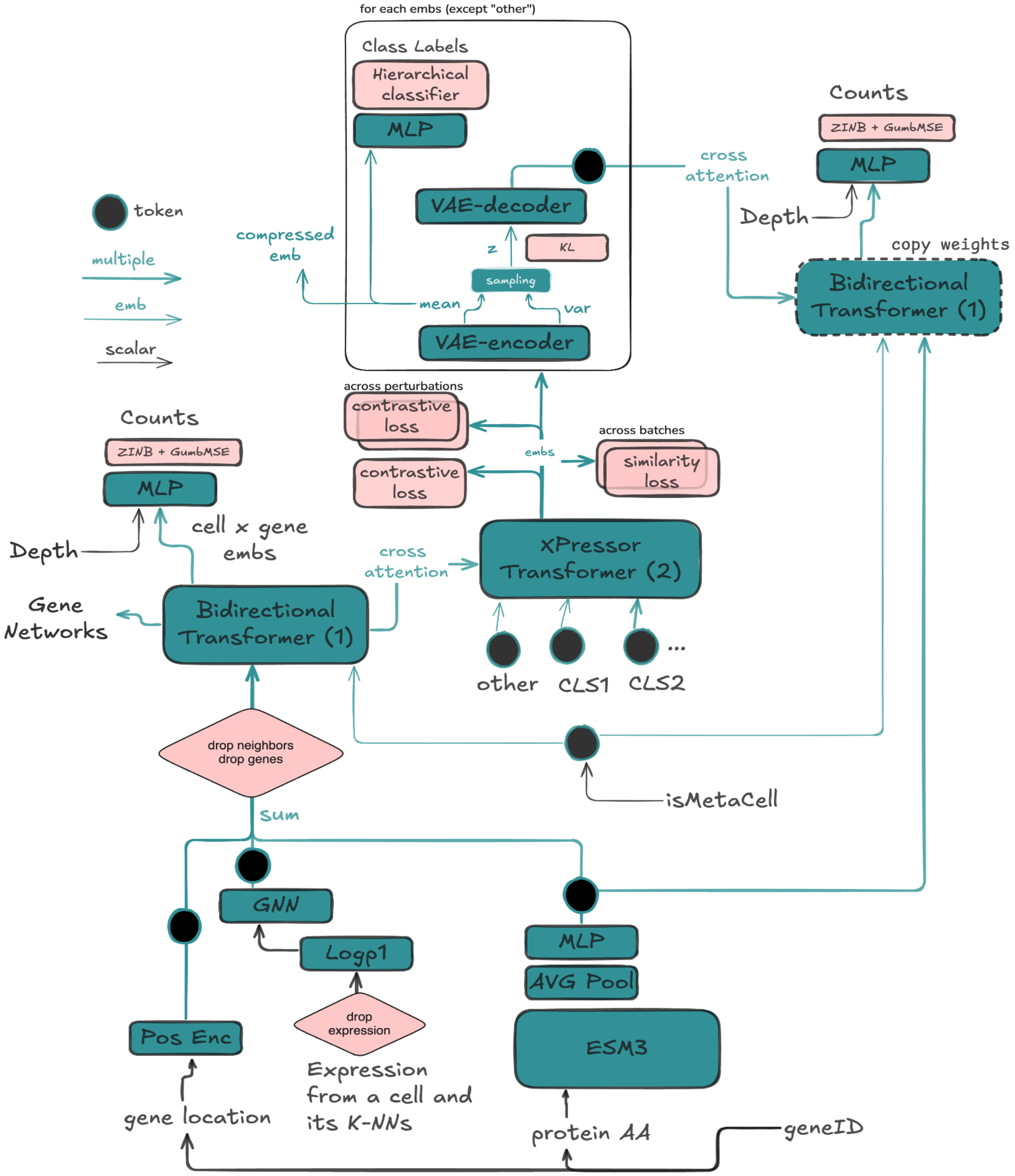


In-depth illustration of the full scPRINT-2's architecture, input, and output with all its main different components and the data flow.

### FIG S2: Whisker plot of the F1-macro scores on the label-projection task of the Open Problem benchmark

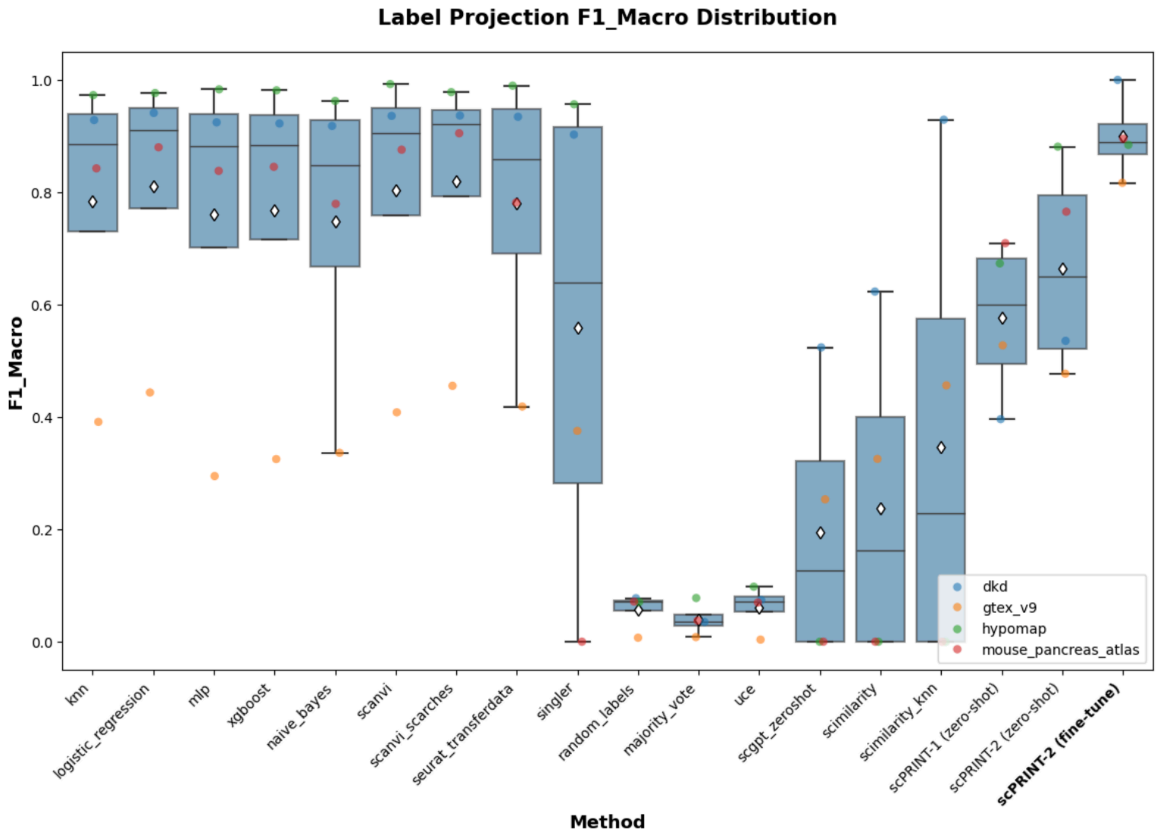


Comparison of scPRINT-1 and scPRINT-2, zero-shot and finetuned, with all other tested methods in Open Problems.

### FIG S3: heatmap of ethnicity prediction relationship across samples

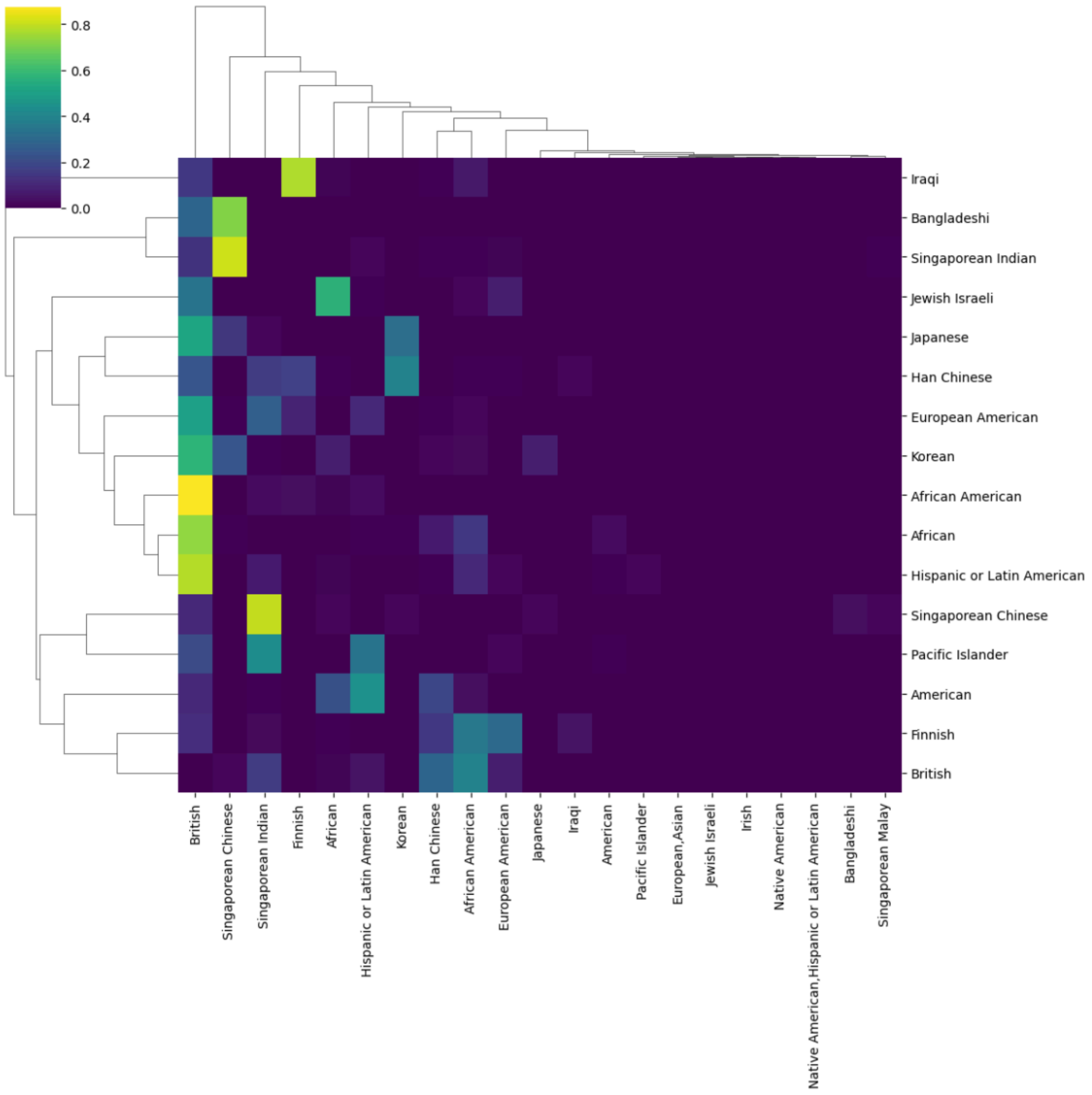


It is generated using labels predicted as top-1 (x-axis) vs second-best prediction (y-axis) across 10,000 random cells for each predicted label from the scPRINT-2 corpus.

### FIG S4: heatmap of organism prediction relationship across samples

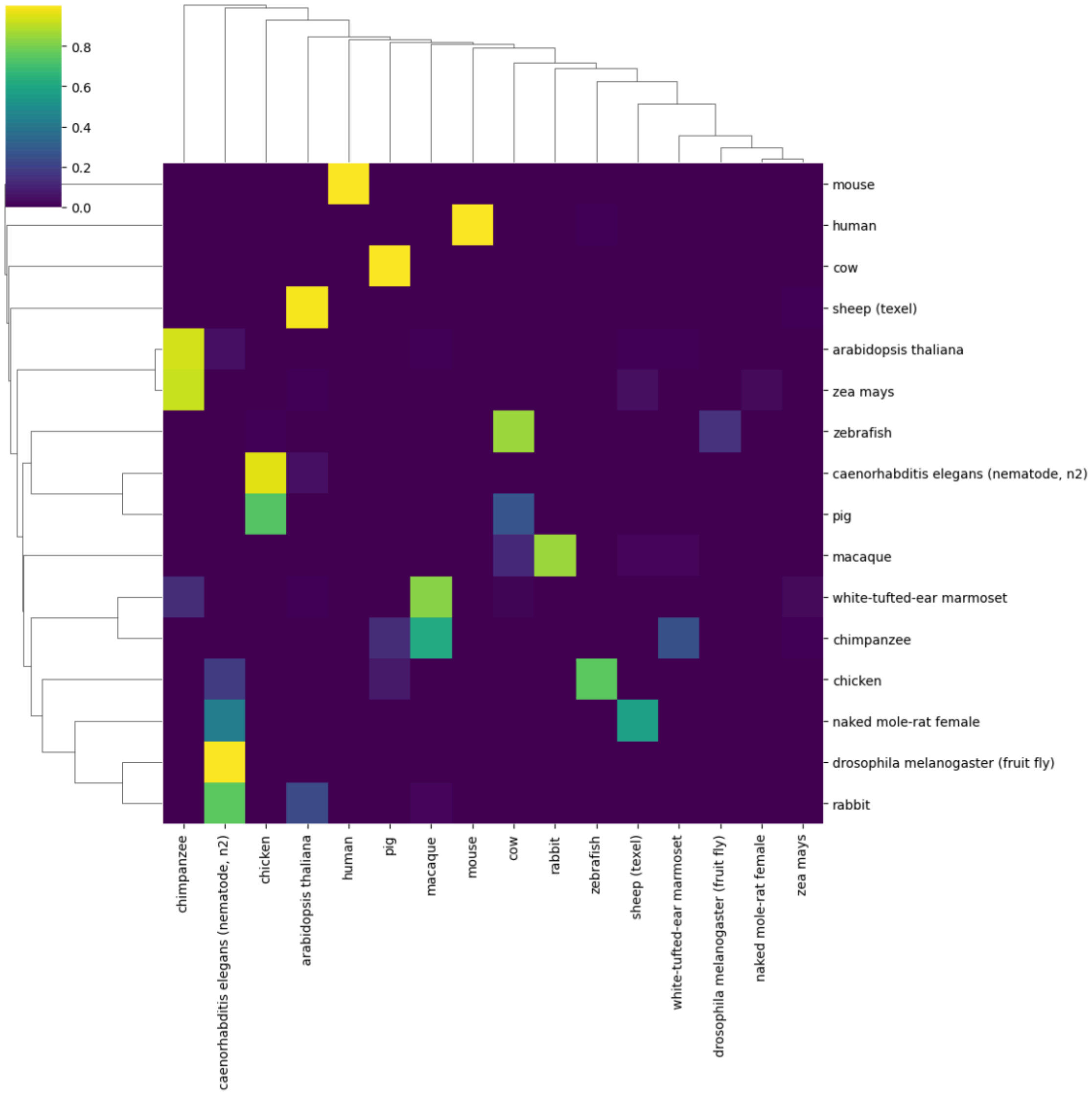


It is generated using labels predicted as top-1 (x-axis) vs second-best prediction (y-axis) across 10,000 random cells for each predicted label from the scPRINT-2 corpus.

#### FIG S5: heatmap of organism prediction relationship using organism embedding similarity across samples

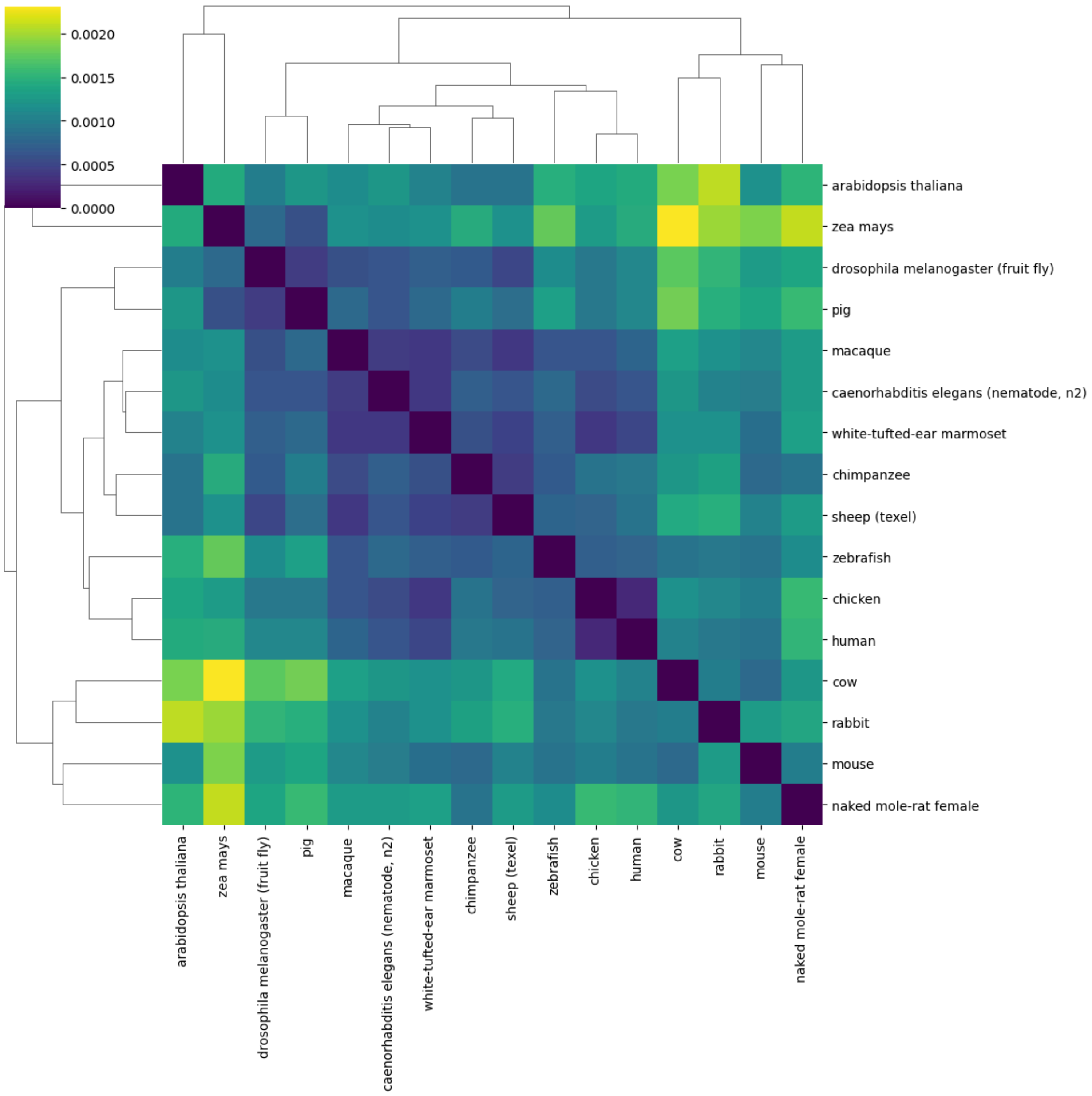


It is generated by averaging the embeddings for each predicted organism across 10,000 random cells for each predicted label in the scPRINT-2 corpus, and using the L2 distance.

### FIG S6: Differential expression plots of the disagreeing cells between scPRINT-2 and ground truth

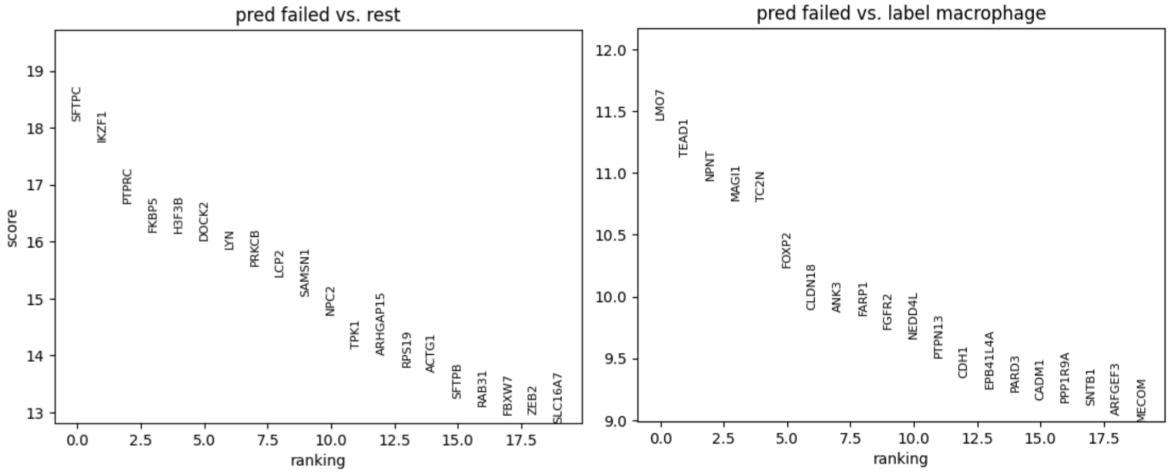


The differential expression is made on the cat/tiger cross-species dataset. “pred failed” is the macrophages labeled as type 2 pneumocytes by scPRINT-2.

### FIG S7: Umap of the smart-seq dataset used in the varying context classification task

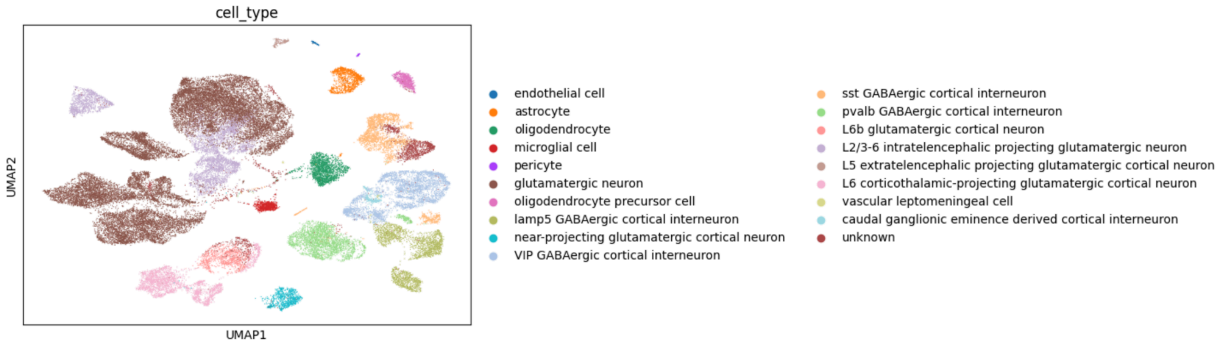


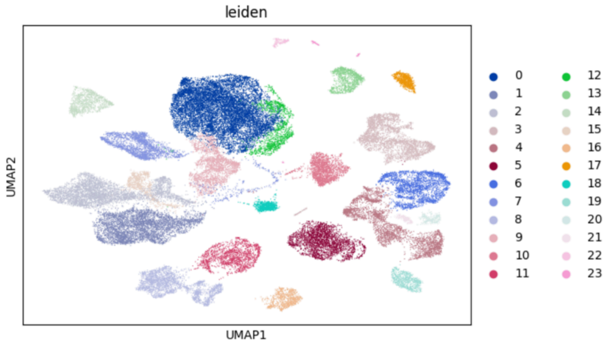


Umap of the cortical areas smart-seq v4 dataset used in the varying context classification task in results section 2, showing Leiden clusters and ground truth cell types.

#### FIG S8: line plot of the classification across varying context length, using the most expressed genes

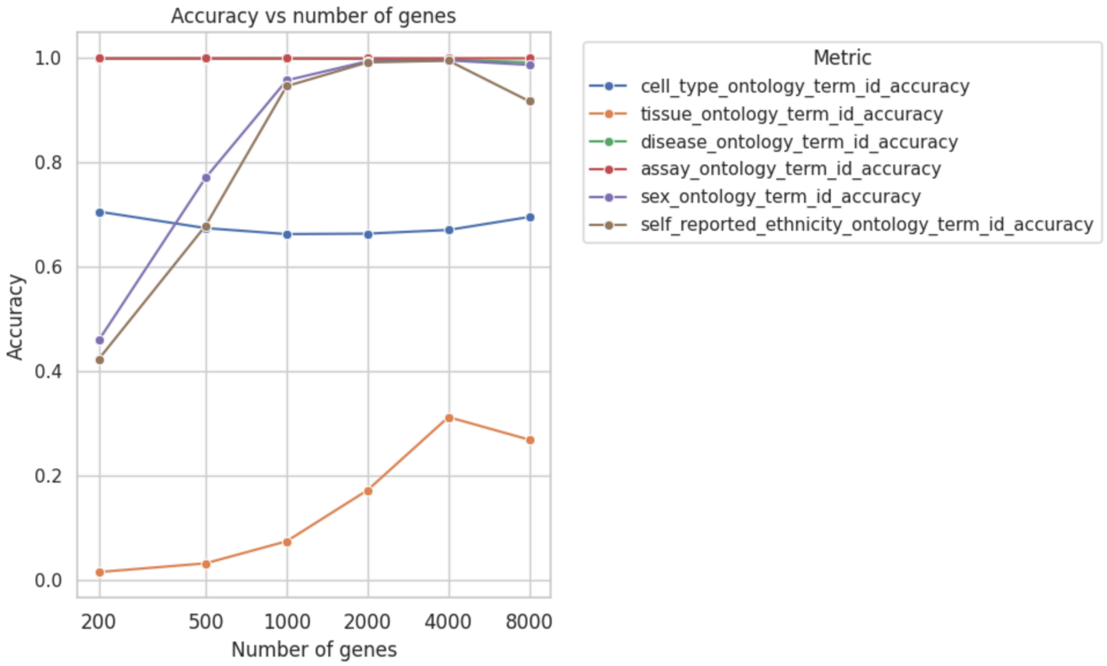


On the same dataset, but this time using the most expressed genes. Meaning each new gene in context is 200 most expressed, then 500, 1000, etc. We can see that while cell types are often defined by their most expressed genes, and thus this doesn't change classification accuracy much, other, more complex labels continue increasing in accuracy as context length increases.

#### FIG S9: Illustration of the multiple perturbations applied to expression data in scPRINT-2

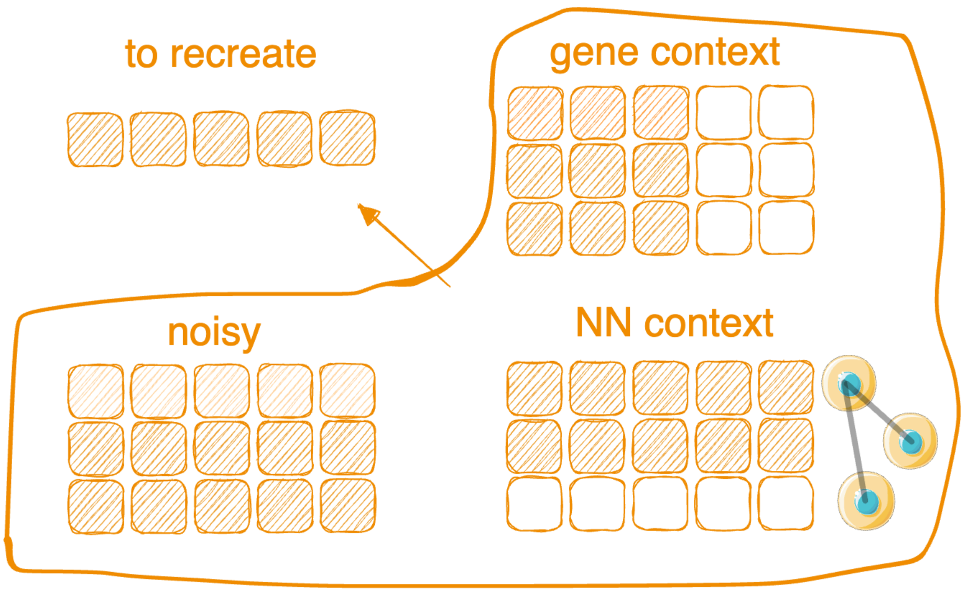


scPRINT can add noise and mask gene expression, modify the number of neighbors, and adjust context lengths.

#### FIG S10: distplot of the non-zero count distribution across cells from the three dataset qualities used

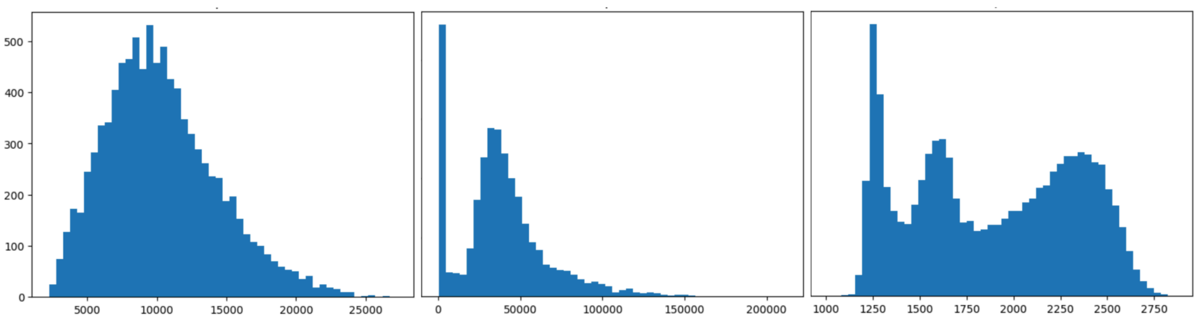


Non-zero count distributions across cells from left: good quality; center: excellent quality; right: poor quality datasets used in our denoising benchmark.

### FIG S11: Umap over scPRINT-2 and PCA embeddings of the Xenium dataset

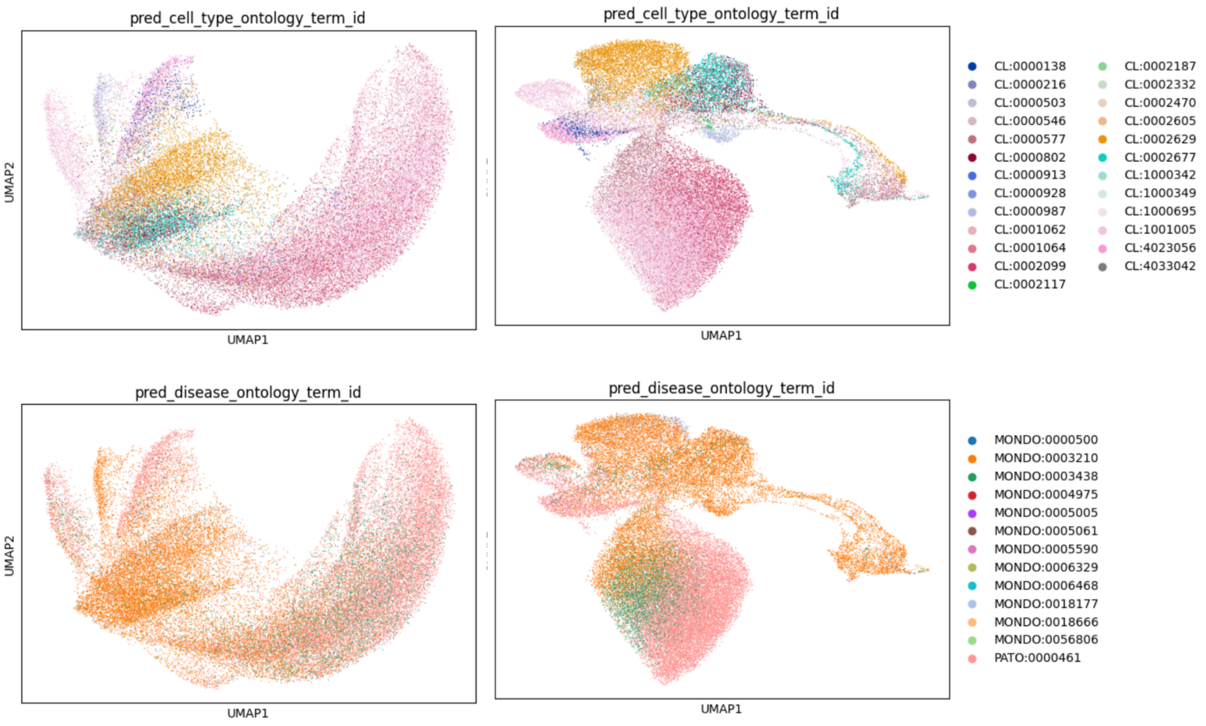


Umap of left: raw PCA expression, right: scPRINT-2 embeddings with scPRINT-2 predicted cell types and diseases.

### FIG S12: Tangram mapping quality plots

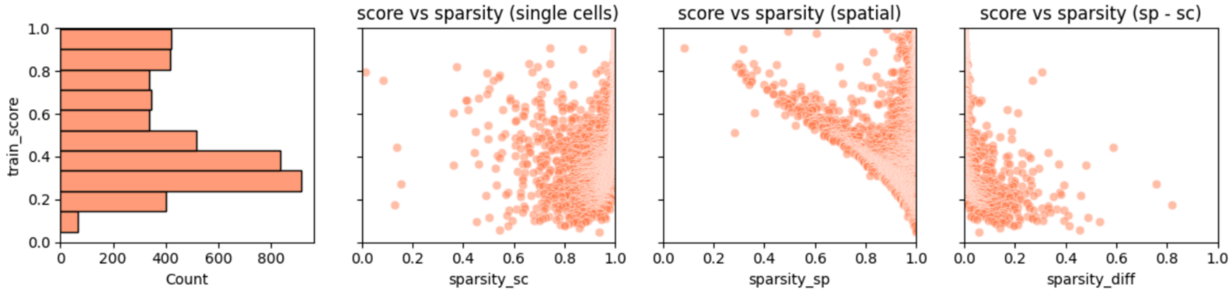


Tangram mapping quality plots on the Xenium skin melanoma datasets and 10v3 skin melanoma datasets.

#### FIG S13: illustration of scPRINT-2's generative imputation mechanism

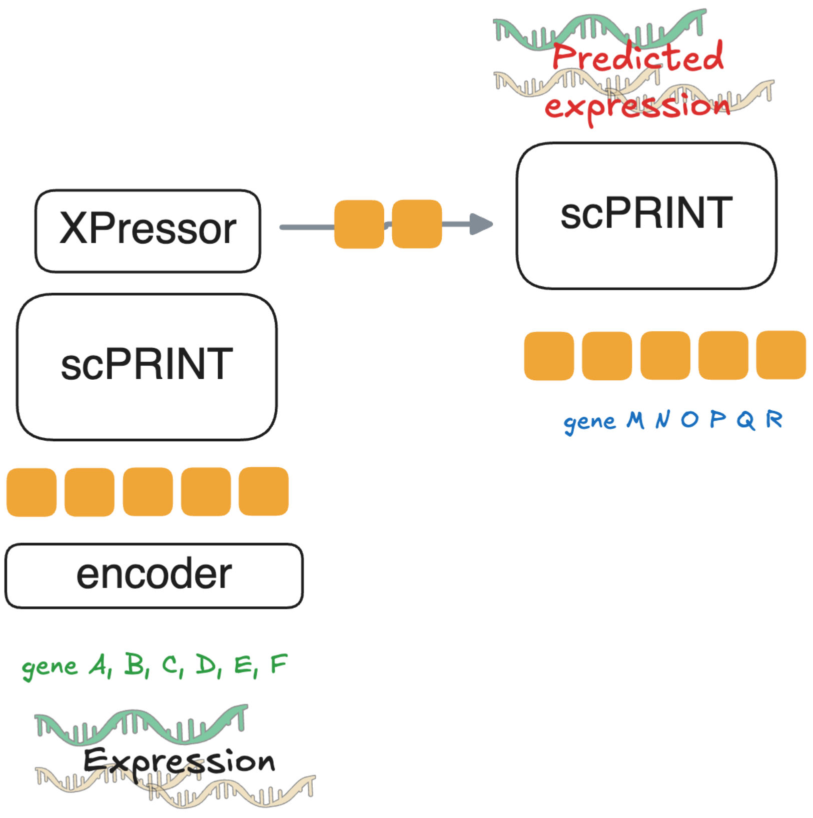


scPRINT encodes all 5000 measured genes into cell embeddings and decodes them on 5000 different unseen gene embeddings.

### FIG S14: spatial plot of the Xenium melanoma dataset with scPRINT-2 predicted cell labels

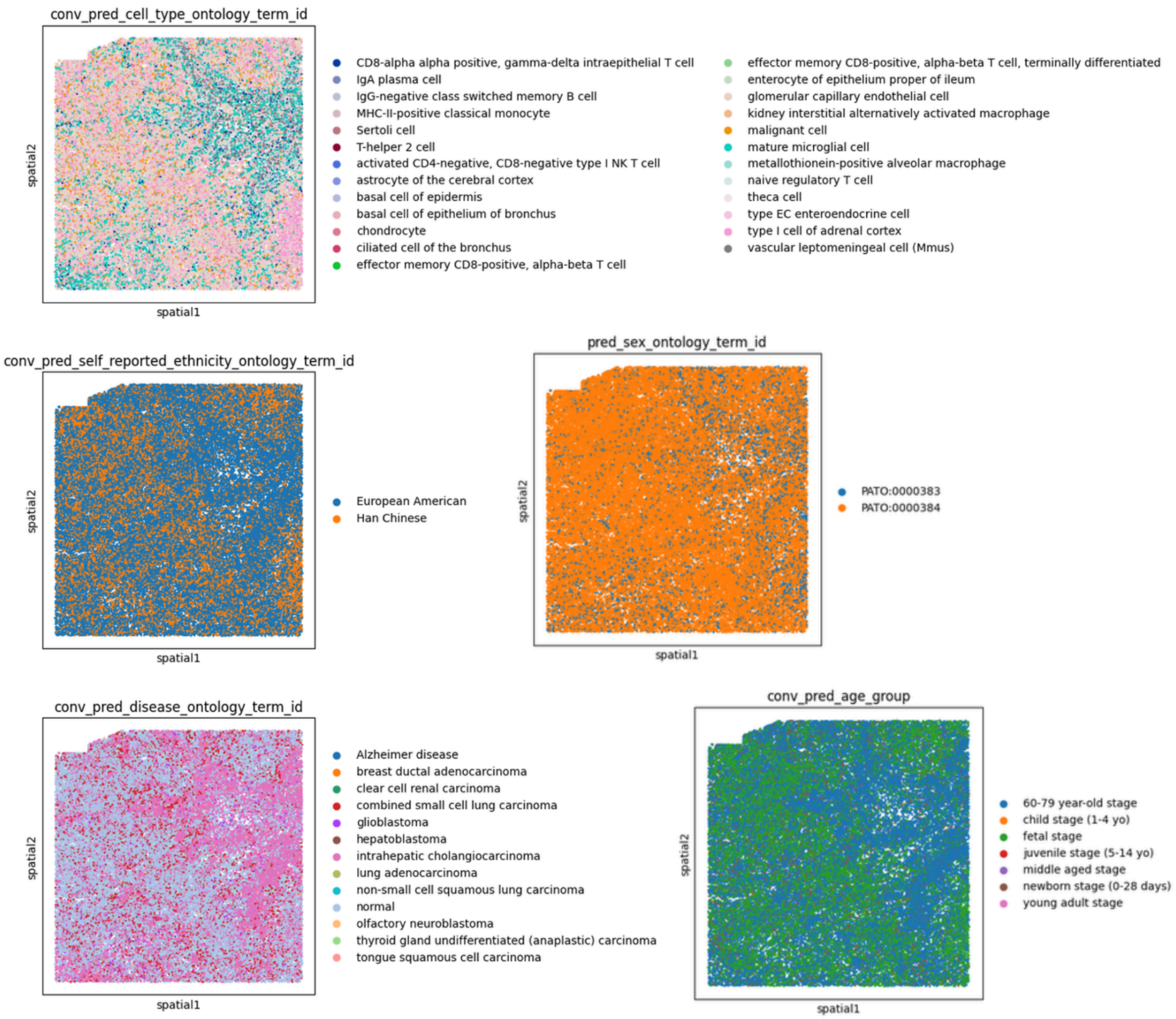


scPRINT-2 predicted cell labels for the disease, age, ethnicity, sex, and cell type labels on top of the selected Xenium skin melanoma patch.

### FIG S15: violin plot comparison of the gene's expression between predicted malignant vs the rest

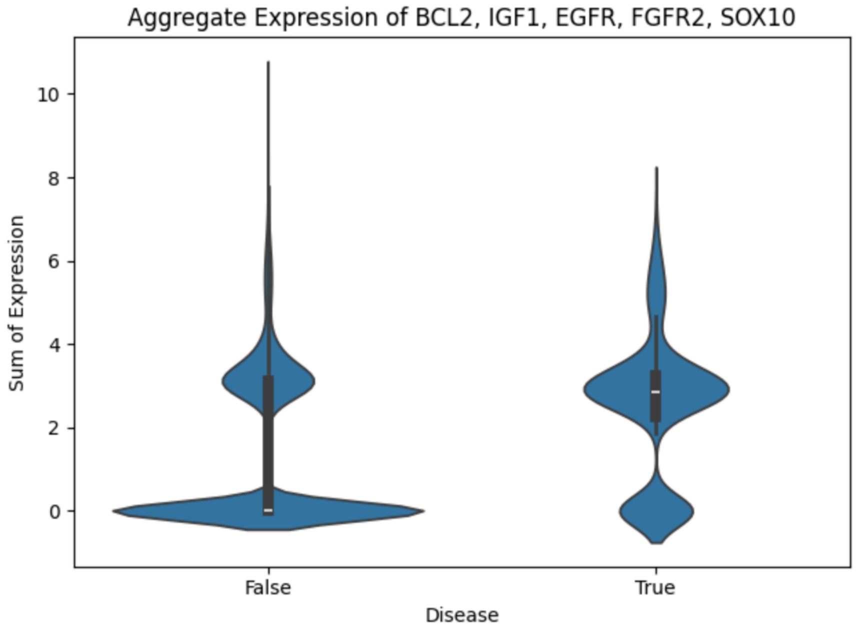


Violin plot showing that BCL2, IGF1, EGFR, FGFR2, SOX10, key melanoma markers are highly expressed in the malignant cell type label group vs the rest, with a p-value of $10^{-234}$

#### FIG S16: differential expression plot of “cancer” disease labelled vs rest in the xenium dataset

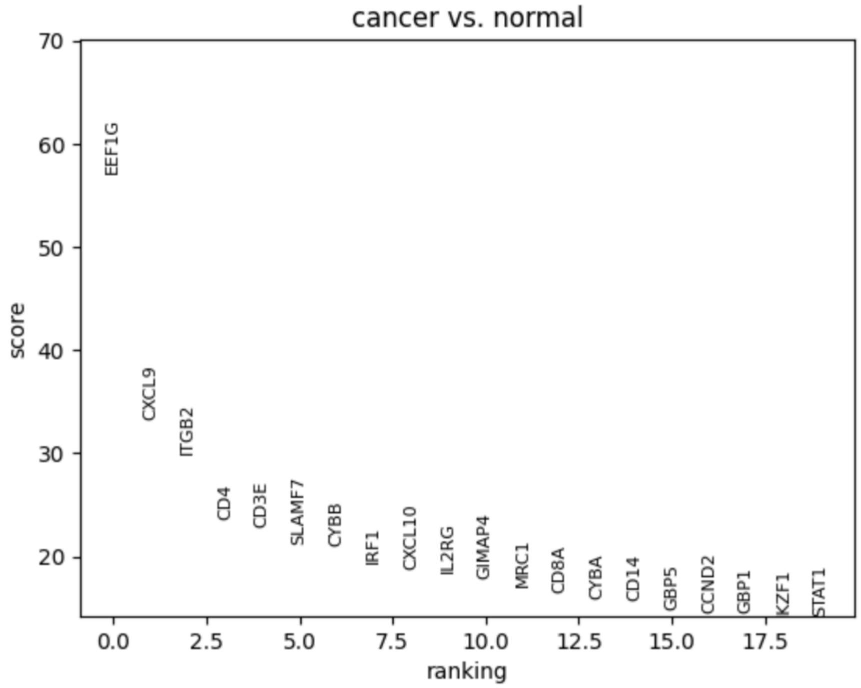


Differential expression plot of cells whose disease label is “cancer” vs the rest in the Xenium skin melanoma dataset

#### FIG S17: Illustration of criss-cross attention

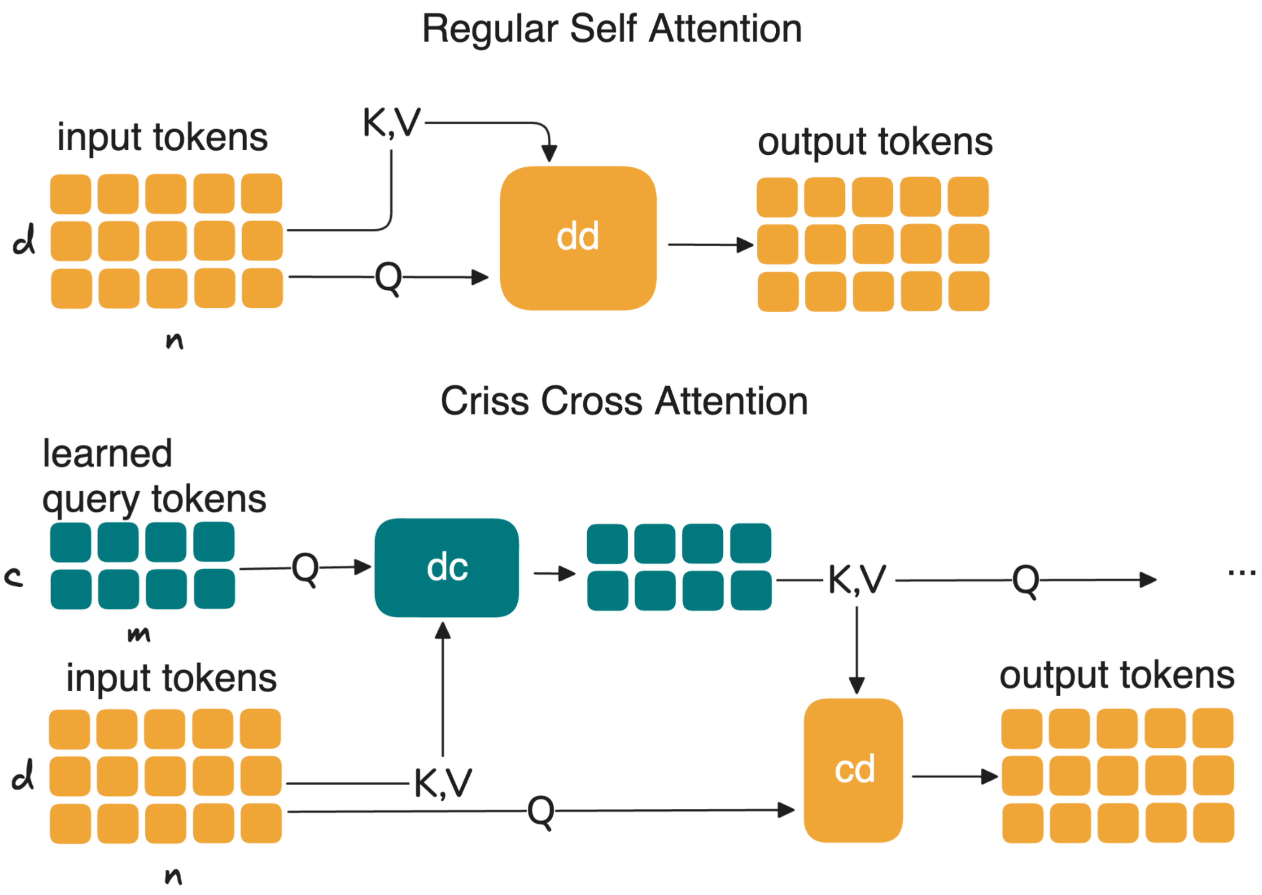


Illustration of our sub-quadratic complexity criss-cross attention mechanism

#### FIG S18: Illustration of the similarity and dissimilarity-based contrastive losses used in scPRINT-2

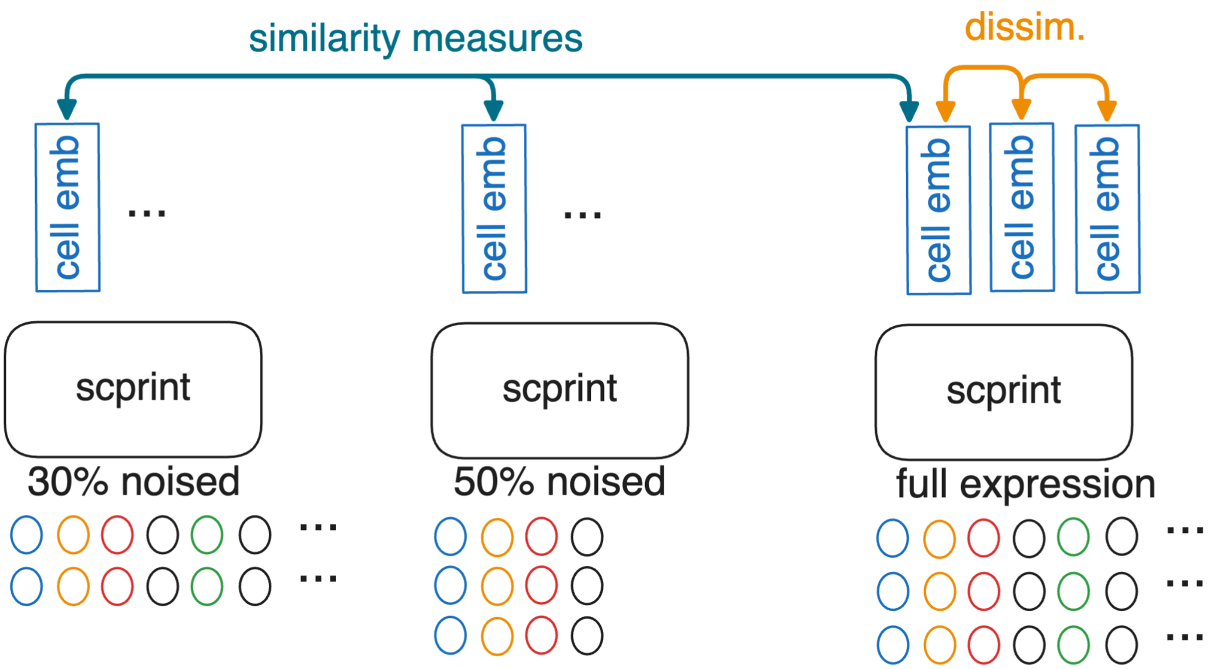


The contrastive losses push embeddings from the same cell at different noise levels to be as similar as possible.

### FIG S19: whisker plot of Open Problems' batch-integration with batch-correction-only scores

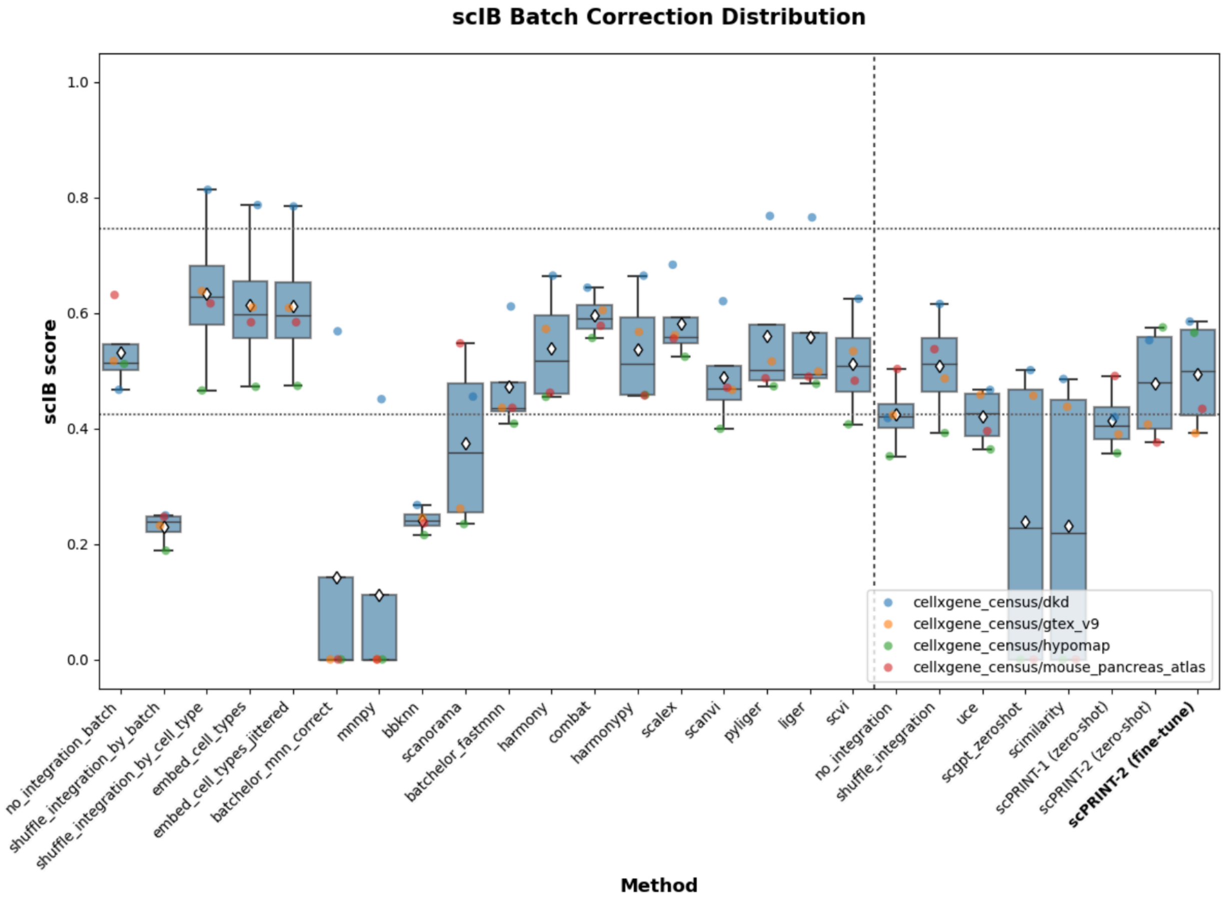


Open Problems' batch-integration with batch-correction-only scores for scPRINT-1 and scPRINT-2 zero-shot, and finetuned, and all other models assessed in open problems.

### FIG S20: whisker plot Open Problems' batch-integration with Bio-conservation-only scores

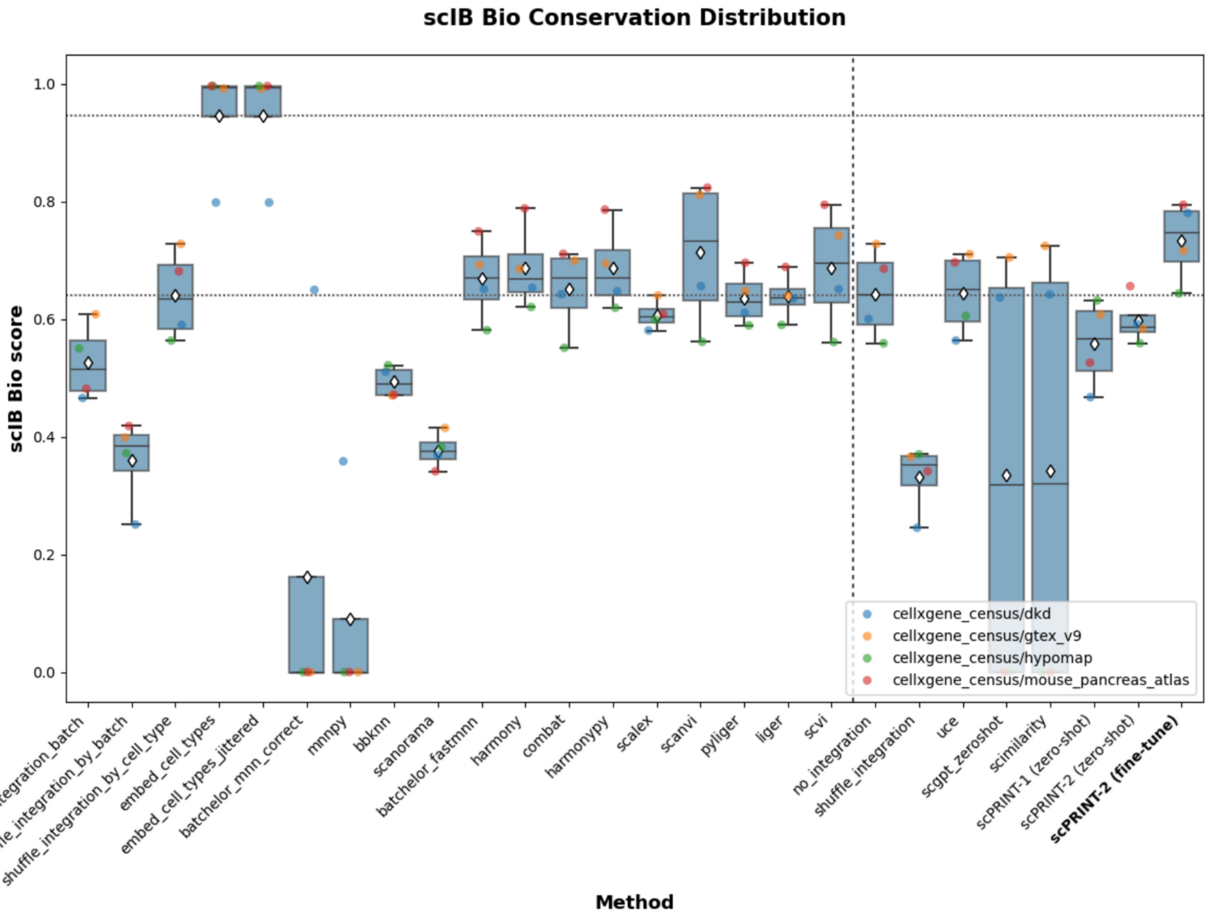


Open Problems' batch-integration with Bio-conservation-only scores for scPRINT-1 and scPRINT-2 zero-shot, and finetuned, and all other models assessed in open problems.

### FIG S21: Umap of scPRINT-2's zero-shot multi-species expression embedding using the full cell-embedding

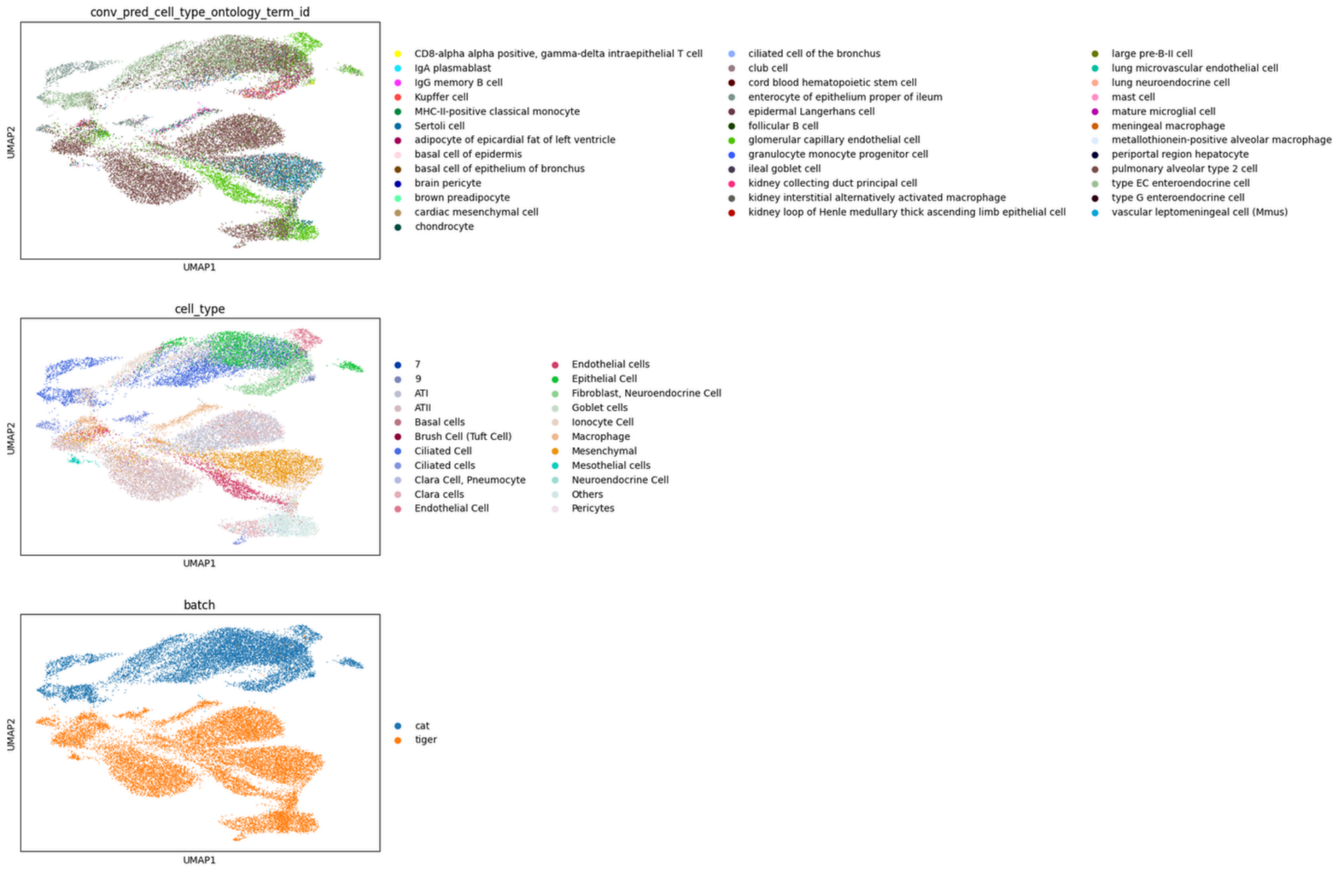


scPRINT-2's zero-shot multi-species expression embedding using the full cell-embedding from top to bottom, scPRINT-2 predicted cell type labels, ground truth cell type labels, and ground truth organism labels.

### FIG S22: barplot of scIB score on scPRINT-2’s multi-species integration

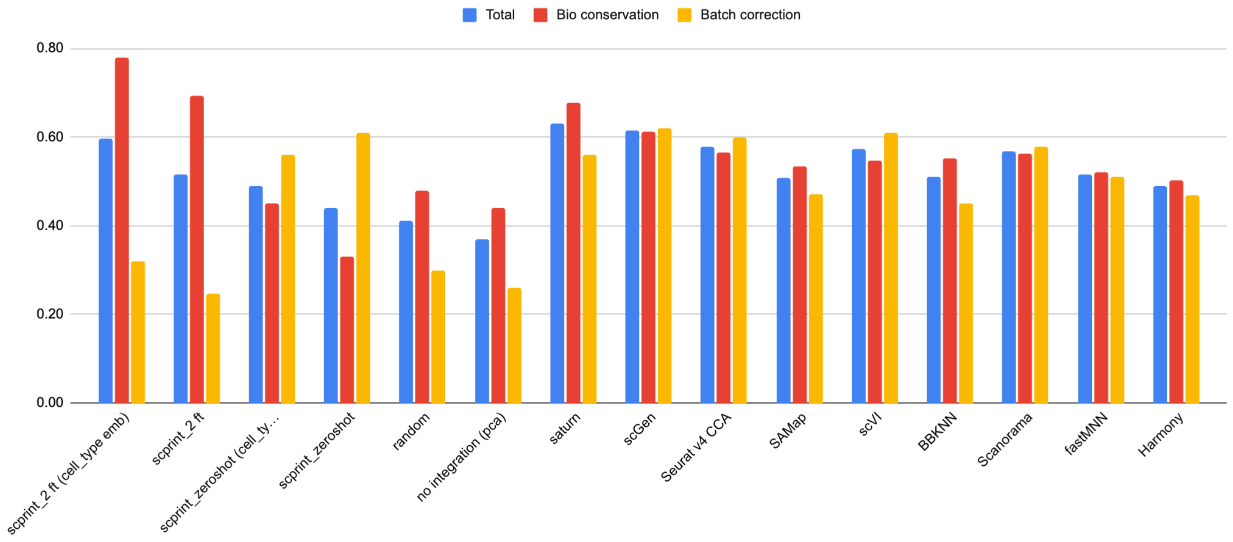


showing total, bio conservation, and batch integration across scPRINT-2 zero-shot, and fine-tuned version using both the full cell-embedding and cell-type-only cell-embedding

### FIG S23: Additional Umaps of the multi-species expression embeddings

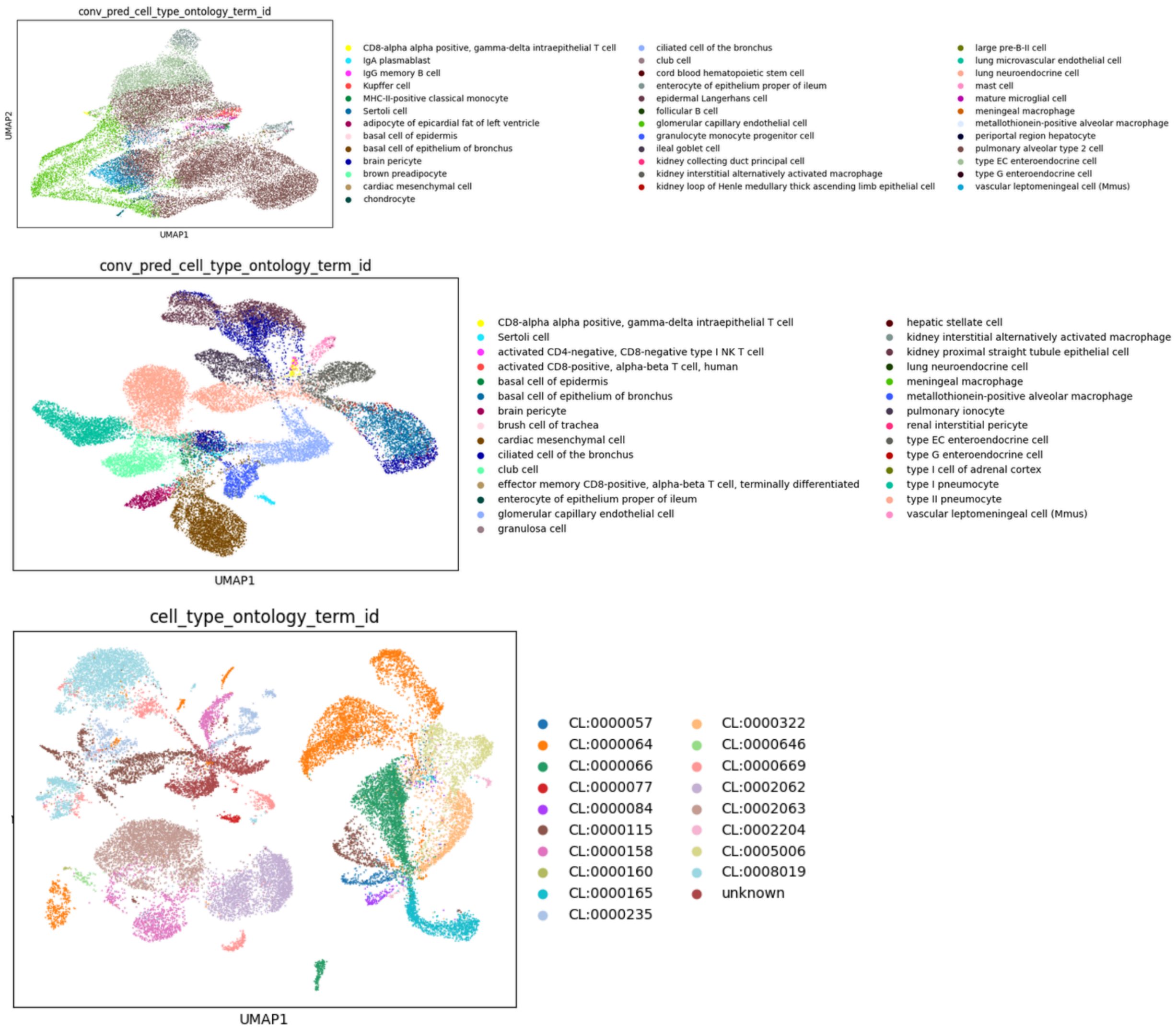


From top to bottom, Umap of scPRINT-2 zero-shot cell-type cell embeddings colored by the cell types predicted by scPRINT-2; of scPRINT-2 fine-tuned cell-type cell embeddings colored by cell types predicted by scPRINT-2; of PCA of the expression data colored by our relabeling of cell types.

### FIG S24: Umap of scPRINT-2's multi-species expression embedding post-finetuning using the full cell-embedding

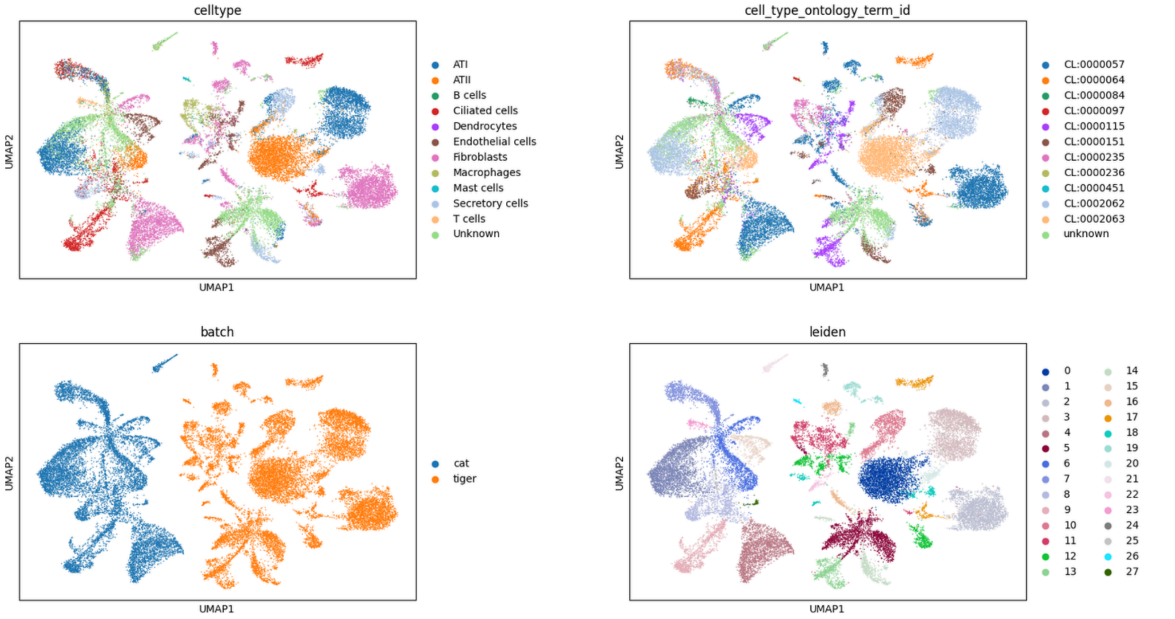


scPRINT-2's multi-species expression embedding post-finetuning using the full cell-embedding from left to right and top to bottom, ground truth cell type, scPRINT-2 predicted cell type labels, ground truth organism labels, and Leiden clusters.

#### FIG S25: Differential expression plot of the human vs mouse dataset from section 4

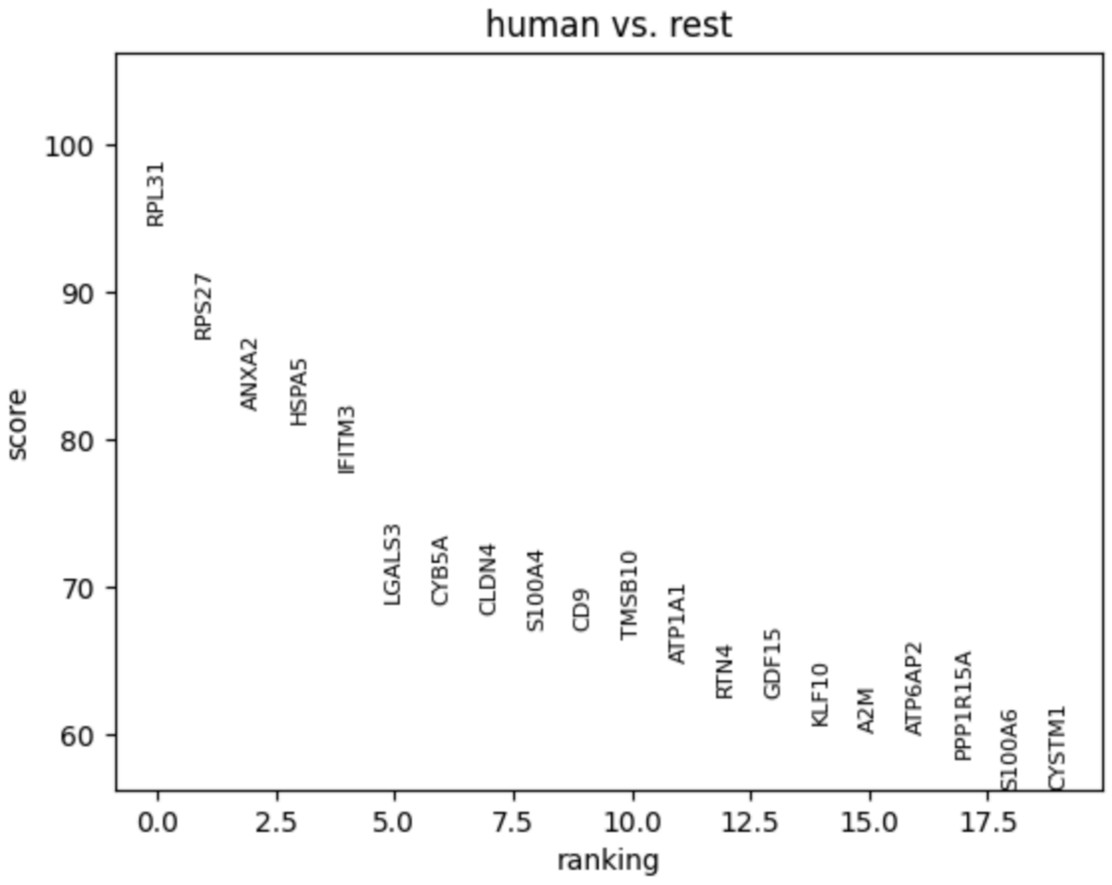


Differential expression plot of the human vs mouse dataset from section 4. Rest is mouse here.

### FIG S26: Over-representation plot of humanized mouse data vs real mouse data compared to human

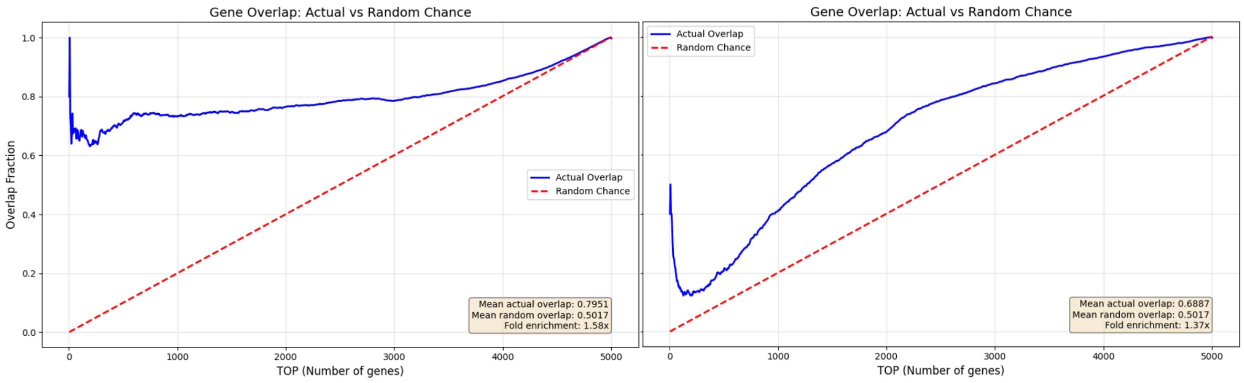


Over-representation plot of differentially expressed genes in scPRINT-2's humanized mouse data vs real mouse data compared to human.

### FIG S27: Over-representation plot of female-like male data vs real female data compared to male

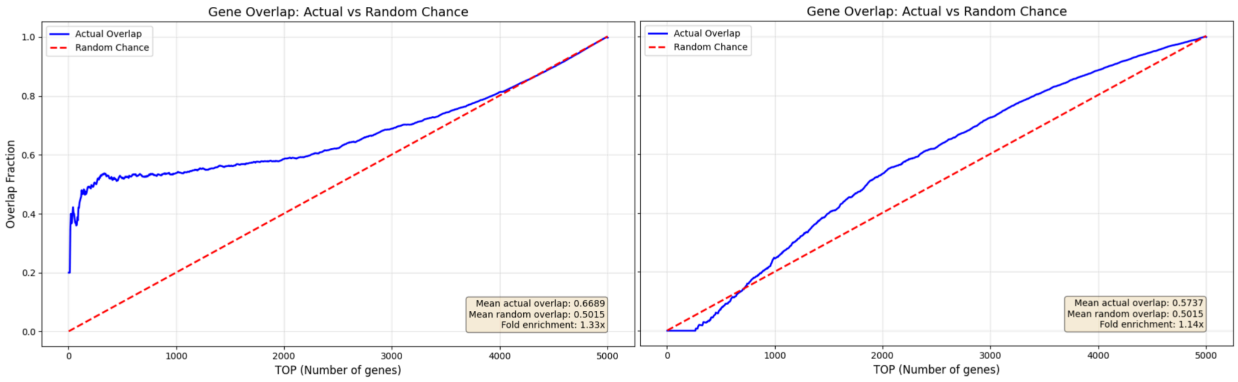


Over-representation plot of top differentially expressed genes in scPRINT-2's female-like male data vs real female data compared to male.

### FIG S28: Dot Plot of Gene-set enrichment analysis over the differential expression analysis of section 4

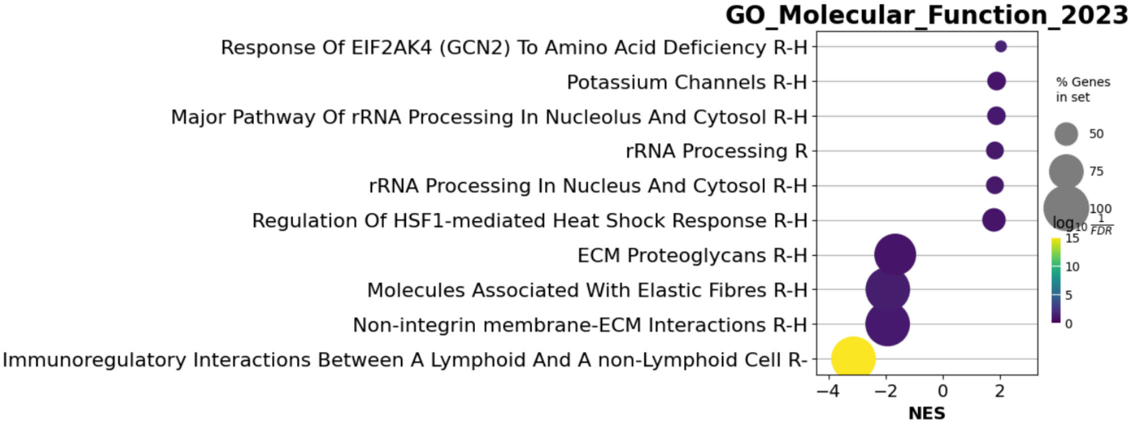


Showing the top 10 most enriched gene sets from the GO molecular function 2023 database.

### FIG S29: Output gene embedding for a non-fully trained model without XPressor architecture

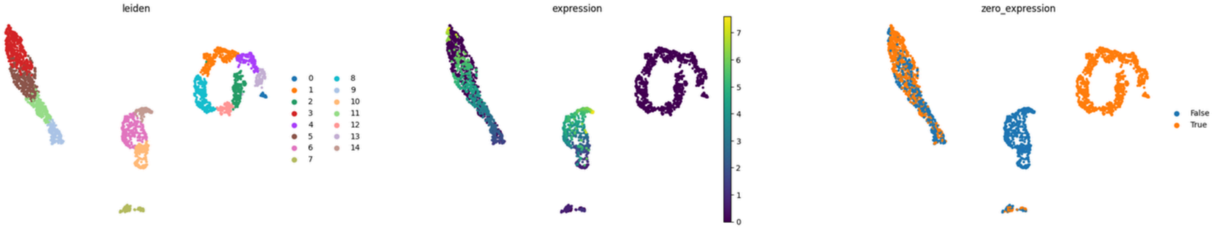


Overlaying in color, from left to right, the Leiden clusters, the expression values, and the zero vs non-zero expression. Despite displaying multiple clusters, the number of enriched pathways in each is still smaller than for a model using XPressor. (see Figure 5)

#### FIG S30: Venn diagram of the different ground truth gene networks

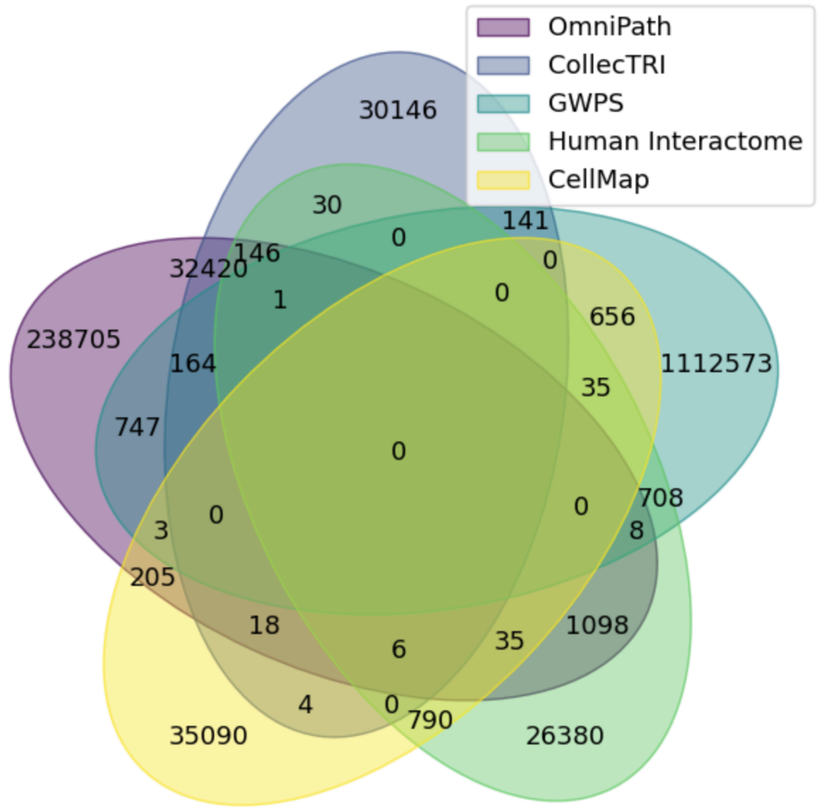


Venn diagram of the different ground truth gene networks showing overlap in the edges using gene symbols over the five ground truths used in our benchmark

### FIG S31: whisker plot of AUPRC-ratio scores for scPRINT-1 and scPRINT-2

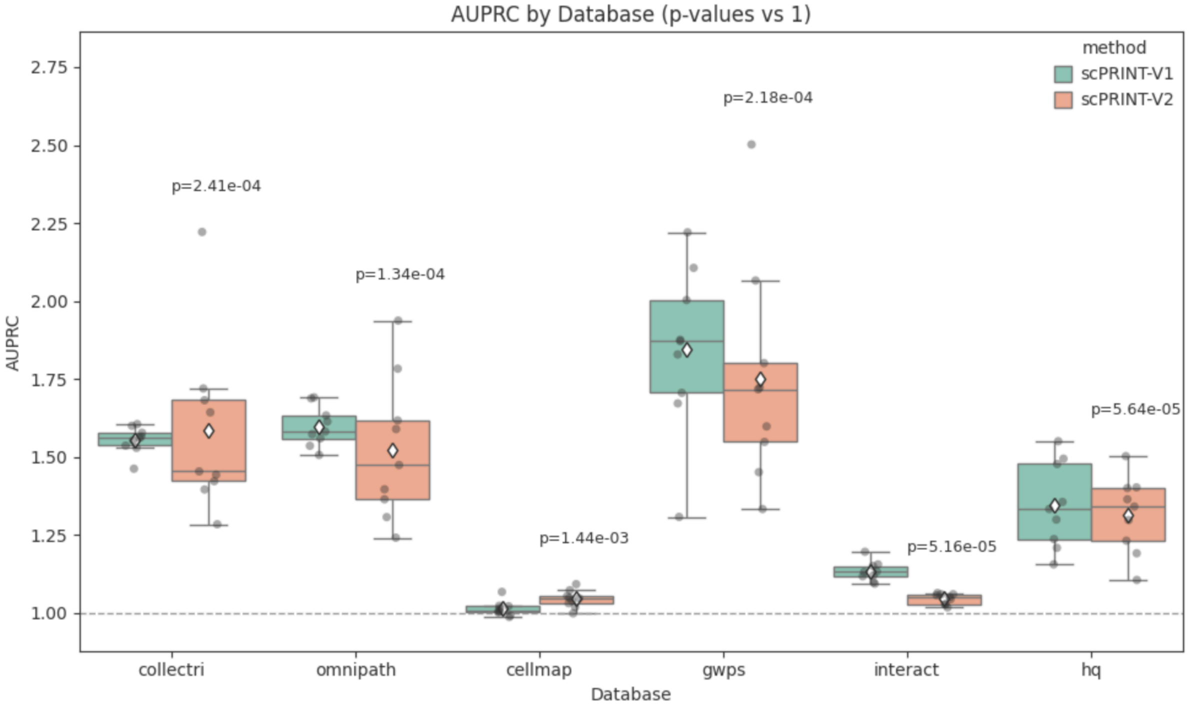


Whisker plot of AUPRC-ratio scores for the benchmark of scPRINT-1 vs scPRINT-2 using their respective GRN-extraction methods, showing that the scPRINT-2 extraction, while highlighting more relevant top connections, remains relatively similar to the scPRINT-1 version on the AUPRC-ratio scores on each of the six ground truth networks.

#### FIG S32: Additional scPRINT-2 generated gene network computed from CDC45

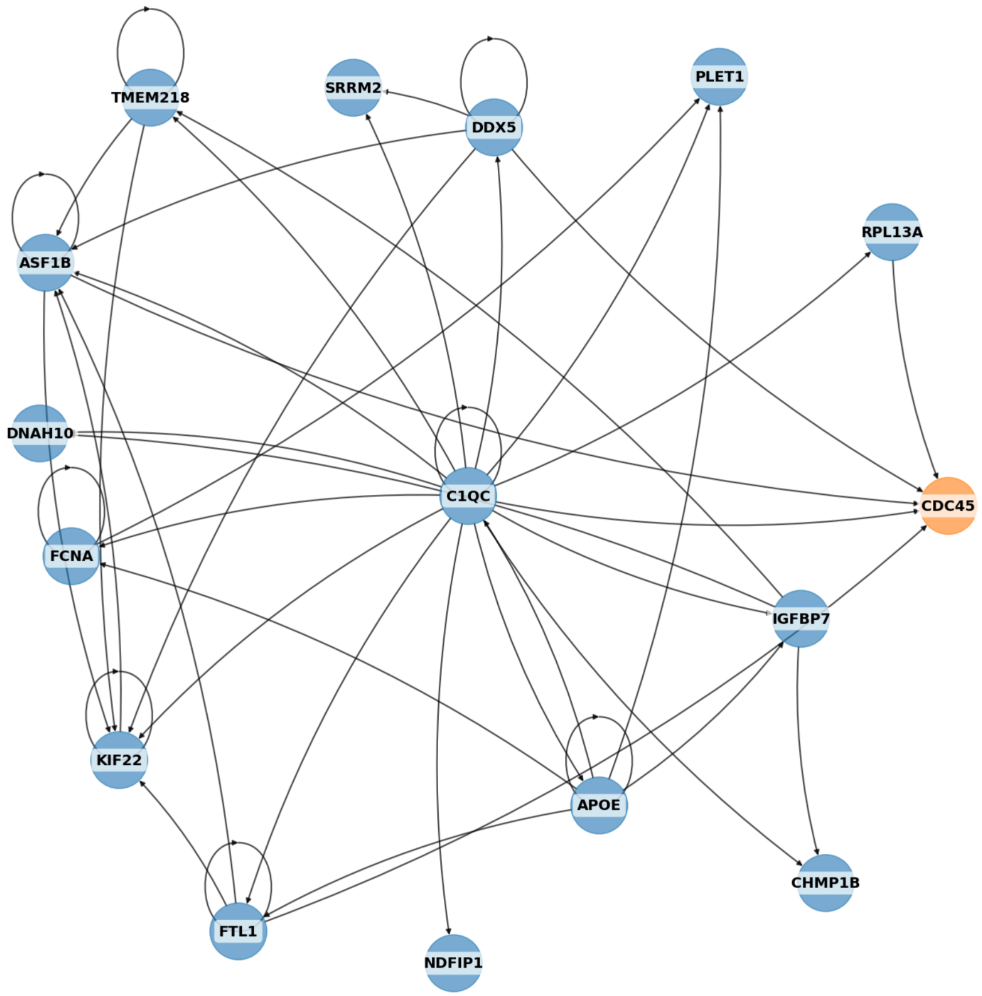


Subpart of the scPRINT-2 generated gene network using CDC45 as a seed gene and computed on 1024 mouse macrophages, showing how these networks can exhibit complex structures.
